# Code for Collagen Folding Deciphered

**DOI:** 10.1101/2024.02.24.581883

**Authors:** Jean-Daniel Malcor, Noelia Ferruz, Sergio Romero-Romero, Surbhi Dhingra, Vamika Sagar, Abhishek A. Jalan

## Abstract

Collagen triple helix folds in two steps: nucleation of three polypeptides at the C-termini followed by zip-chain like propagation. The triple helices found in all domains of life as well as viruses contain upto 6000 amino acids in each polypeptide that are also frequently interrupted with non-helical sequences that disrupt folding and reduce stability. Given the length of polypeptide and the disruptive interruptions, compensating mechanisms that stabilize against local unfolding during propagation and offset the entropic cost of folding the long polypeptides are not fully understood. Here, we show that the information for correct folding of collagen triple helices is encoded in their sequence as interchain electrostatic interactions. In case of humans, disrupting these interactions causes severe to lethal diseases.

**Key Result:** Collagen triple helices found in all the three domains of life as well as viruses have converged on similar mechanism to fold correctly.

## Introduction

Collagens are highly abundant human proteins that provide structure and strength to tissues, and bind cell-surface and secreted proteins to regulate key biological processes, including tissue homeostasis and blood clotting. All collagen proteins contain a characteristic domain called a triple helix composed of three supercoiled polypeptides. In humans, collagen genes encode 44 polypeptides, also called *α*-chains, that self-assemble into 28 distinct collagen types (I-XXVIII).(*1*) The *α*-chains contain a repetitive three amino acid sequence Gly-Xaa-Yaa, where glycine occupies every third position and Xaa and Yaa are non-glycine amino acids. Upon trimerization, the recurrent glycines sequester into a tightly packed core of the triple helix.(*2*) Consequently, their mutation to other amino acids perturbs the structure resulting in delayed folding and reduced stability.(*3, 4*) This is the leading cause of heritable collagen-related diseases.(*5*) Direct study of collagen folding and stability is complicated by the propensity of native collagens to form insoluble aggregates *in vitro*. Fortunately, synthetic peptides containing sufficient Gly-Xaa-Yaa repeats intrinsically self-assemble into a triple helix. Investigations using such peptides suggest that triple helices fold via a nucleation-zipper mechanism, where three peptides nucleate at the C-terminal and then propagate towards the N-terminal like a zip-chain.(*6*)

The *α*-chains of human collagens contain up to 330 Gly-Xaa-Yaa repeats. Intuitively, until the propagating triple helix has reached a critical length sufficient to sustain further folding, the propagation phase is expected to be entropically disrupted due to the substantial length of the *α*-chains. In some collagens, the perfect triplet repeat pattern is also frequently interrupted by non-helical sequences that lower triple helix thermal stability and disrupt folding(*7*). In such cases, the triple helix must renucleate after the interruption and also compensate for the loss in local stability and folding rate. A collagen-specific chaperone heat shock protein (Hsp) 47 resident in the endoplasmic reticulum has been shown to recognize unique sites on the triple-helical domain of some human collagens.(*8*–*10*) This interaction between Hsp47 and collagen triple helices is believed to provide stability against local unfolding while offsetting the entropic loss.(*11*) However, several experimentally observed features of collagen folding are not fully cosistent with the Hsp47-chaperoned folding paradigm. Prominent among these is the observation that Hsp47 does not recognize some types of collagens, or does so only weakly.(*12*) Furthermore, on a structural level, polypeptides within a triple helix adopt a one-amino acid staggered alignment with respect to each other. But, non-canonical polypeptide alignments are not uncommon.(*13*) Hsp47 binding does not explain how polypeptides avoid incorrect alignment during folding. Hsp47 binding also does not explain how triple helices in native collagen re-nucleate after an interruption and compensate for the loss in stability and folding rate. Moreover, in addition to eukaryotic collagens, eukaryotes(*14*), prokaryotes(*15*) and viruses(*16*) also encode collagen-like proteins containing the characteristic Gly-Xaa-Yaa repeats that form stable and functional triple helices(*17*–*19*). While human *α*-chains are ∼330 triplet long for the most abundant types of collagen, collagen-like polypeptides in prokaryotes can reach lengths of upto 2000 triplets and contain a greater number of interruptions.(*15*) How such long polypeptides mitigate local unfolding during the propagation phase while also compensating for the disrupting effect of interruptions is currently not understood.

The information for correct folding of most proteins is encoded in their amino acid sequence. Our results here suggest this to be true for collagens as well. We show that human collagens and collagen-like proteins in the domains archaea, bacteria and eukarya (excluding human collagen orthologues) and viruses contain amino acid motifs capable of forming geometrically specific lysine-glutamate and lysine-aspartate salt bridges. The salt bridges increase kinetic stability of model triple-helical peptides as well as multiple native collagens tested here. We find each collagen subtype to contain an average of 50 salt bridges. By comparison, only a few binding sites for Hsp47 have formally been identified in human type II and III collagens.(*20*) Importantly, incorrect alignment of the chains in the respective triple helices dramatically decreases the number of possible salt bridges in all collagen subtypes. We also find that non-collagenous interruptions are frequently flanked by a higher density of salt bridges. Importantly, salt bridges are found to be distributed throughout the length of the triple-helical domain of human collagens. This combined with their ability to increase kinetic stability suggests that salt bridges act as local electrostatic clamps that lock the folded regions of the triple helices in place and allow the remaining regions to propagate. Therefore, our results provide a general mechanism for how collagen triple helices avoid local unfolding while also precluding unproductive folding intermediates. Furthermore, we find that mutations that disrupt salt bridges are associated with severe or lethal diseases, which has important consequences for understanding why some mutations in human collagens cause more severe phenotypes than others.

## Results

### Salt-bridge-forming triplets anomalously frequent in collagen and collagen-like proteins

Excluding glycine, there are 361 residue pair combinations possible in the Xaa and Yaa positions of a Yaa-Gly-Xaa triplet. Analysis of human fibrillar collagens, representing 9 of the 44 collagen *α*-chains, has previously revealed that several triplets including KGE and KGD occur with a frequency greater than that expected from a random distribution of amino acids.(*21*) Such anomalous frequency suggests selection pressure and thus a prominent biological role in folding and/or function. These triplets possess an ammonium group (on the side chain of lysines) and a carboxylate group (on the side chains of aspartate or glutamate) in close proximity, making them likely to participate in electrostatic salt bridge interactions. We revisited the analysis of KDG or KGE triplets frequency within all 44 human collagen *α*-chains. Additionally, we also analyzed the triplet frequency in collagen-like proteins from archaea, bacteria, eukarya and viruses to understand how these compare to the human collagens.

We defined triple-helical domains as 6 or more contiguous repeats of Yaa-Gly-Xaa triplets. Humans contained 468 collagen domains while archaea, bacteria, eukarya and viruses accounted for 58002 collagen-like protein domains (**supplementary table S1 and supplementary data 1**). For sequence analysis, the occurrence of each Yaa-Gly-Xaa triplets was first predicted based on the observed individual frequency of amino acid residues in the Xaa and Yaa positions within the sequences of human collagen as reported previously(*21*) (see methods for details). This prediction was then compared to the observed occurence of Xaa and Yaa pair combinations in Yaa-Gly-Xaa triplets.

Triplets with Z-scores greater than 3 or less than -3 (more than three standard deviations away from the mean of the difference between observed and predicted values) were considered as anomalously frequent. Z-scores were plotted against relative abundance, which is defined as the percent observed instance of a triplet against total number of triplets in collagens or collagen-like proteins. This was done to understand if triplets with anomalously high frequency also have high relative abundance. As shown in **figure 1a**, KGE and/or KGD triplets are Z-score outliers and thus anomalously frequent not only in the *α*-chains of human collagens but also in collagen-like proteins from the three domains of life and viruses. The KGE and KGD triplets also have high relative abundance ranging between 3 and 6% in both collagens and collagen-like proteins. In the case of viral collagen-like proteins, KGD accounts for a stupendous 13% of all observed triplets. In eukayotes, the PGP triplet, or OGP following the post-translational modification of proline into 4(*R*)-hydroxyproline (noted O), is by far the most abundant. A succession of OGP triplets assembles into the most thermally stable triple helix, with any individual mutation at the Xaa or Yaa position resulting in destabilization. In particular, in model triple-helical peptides, a hydroxyproline to lysine mutation decreases the triple helix thermal stability by 10°C, and a proline to glutamate or aspartate mutation destabilizes it by 7°C or 4°C respectively.(*22*) Yet, mutating both hydroxyproline and proline to lysine and glutamate results in a moderate thermal stability decrease of 5-8°C, suggesting a compensating stabilization mechanism stemming from salt bridge formations.(*23*)

**Fig. 1.**
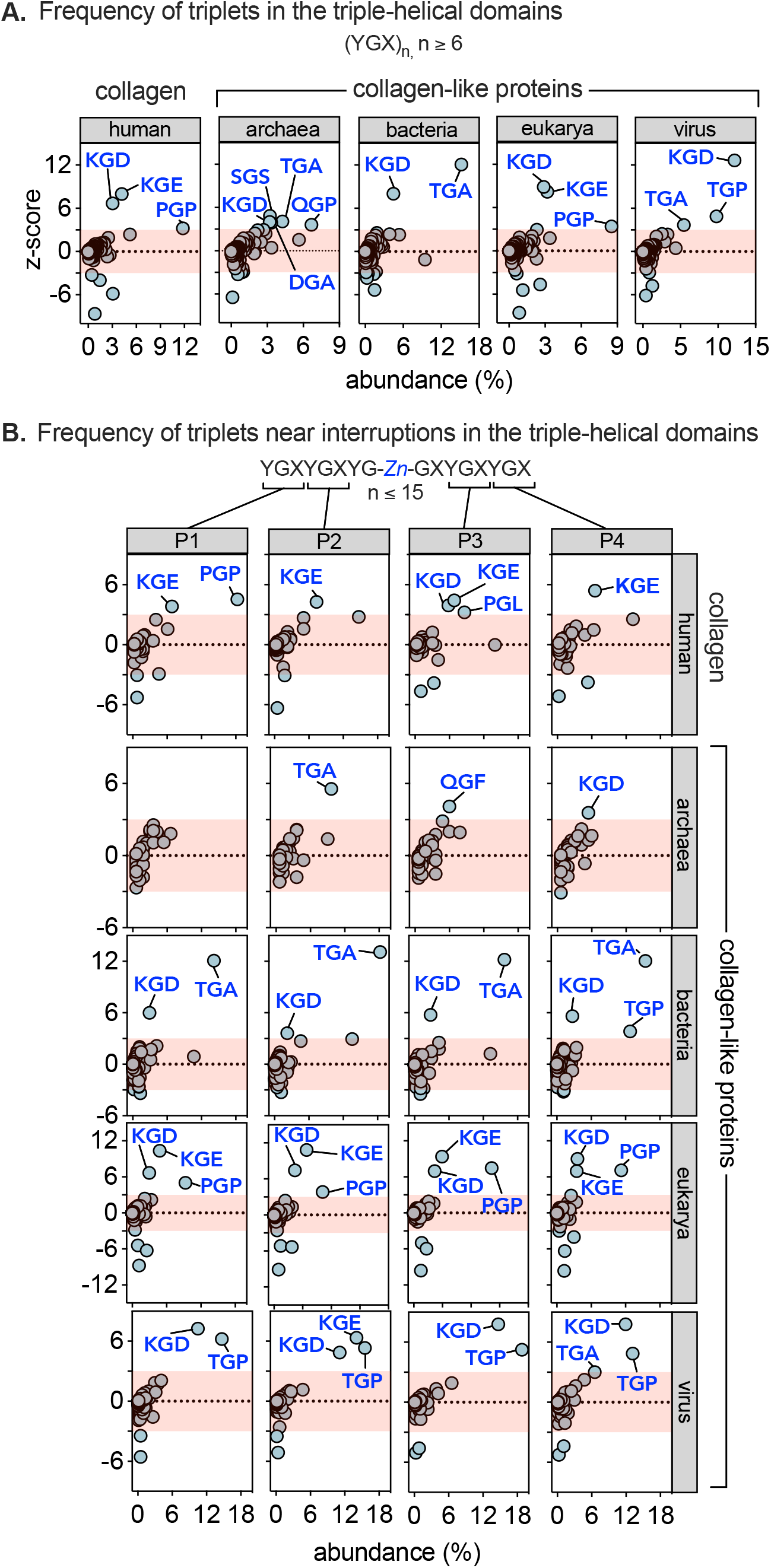
Electrostatically charged triplets are anomalously frequent in collagen and collagen-like proteins. (a) Frequency of Yaa-Gly-Xaa triplets in the uninterrupted triple-helical domains (a) and surrounding the interruption in the 44 known human collagen *α*-chains and in the collagen-like proteins from humans, viruses and the three superkingdoms. In the case of interruptions, the triplet frequency was determined in the four triplets (labelled P1-P4 surrounding the interruptions. *Z* denotes the interruption of length 0 ≥ *n* ≤ 15.

Threonine containing triplets TGA and TGP are also found to be anomalously frequent with high relative abundance in archaea, bacteria and viruses but not in human collagen or collagen-like proteins in eukarya. Bacteria lack the enzyme prolyl hydroxylase required for post-translational modification of hydroxyproline. The high abundnace of threonine in place of hydroxyproline has been rationalized based on the ability of bacteria to O-glycosylate threonines(*24*), which presumably increase triple-helical stability via water-mediate hydrogen bonds(*25*). abundance (%)

We next analyzed the frequency of Yaa-Gly-Xaa triplets that surround interruptions in human collagens and collagen-like proteins. Bella et al. recently proposed a simple and elegant nomenclature for the many different and complex interruptions possible in collagen sequence. In this nomenclature, the interruptions are denoted according to sequence length. For example, a deletion of one amino acid from the triplet repeat sequence is denoted as G1G and a deletion of three or more amino acids is denoted G3G to G(Z)_n_G, where Z denotes interruption and subscript *n* its length up to 15 amino acids. Following this nomenclature, we identified 394 interruptions in human collagens and 33341 interruptions in collagen-like proteins (**supplementary table S1 and supplementary data 2**). Next, we determined the frequency of Yaa-Gly-Xaa triplets focussing only on the sequence surrounding the interruptions. Starting from the N-termini, the two triplets on either side of the interruptions were labelled P1-P4. As shown in **figure 1b**, KGE and KGD are anomalously frequent in P1-P4 not only in human collagens but also in collagen-like proteins from bacteria, eukarya and viruses. Archaeal collagen-like proteins show anomalous frequency of KGD in only the P4 position. The relative abundance of the KGE and KGD triplets surrounding interruptions in collagen-like proteins of bacteria and eukarya range between 2 and 6%. Among all analyzed groups, viruses show the highest relative abundance of these triplets at 10-12% in all four positions surrounding the interruptions. The consistent observation that KGE and KGD triplets are anomalously frequent in the triple-helical domains of human collagens and collage-like proteins in the four groups and that they are also enriched in the triplets surrounding interruption sites suggests a prominent and likely common role in stability, folding and/or function.

### KGE and KGD triplet mediated salt bridges are widely distributed in all human collagens

We established the anomalously high frequency of KGE and KGD triplets in human collagens via analysis of single collagen *α*-chains. However, which of these triplets are capable of forming salt bridges upon trimerization, how many such salt bridges are feasible in a given collagen subtype, what is their distribution within the triple-helical domain and how they are located vis-à-vis interruption is currently not known. To this end, we aligned the 44 collagen *α*-chains into the triple-helical stoichiometries observed in humans(*1*) and determined the number and location of salt bridges in the triple-helical domains and also surrounding the interruptions (**supplementary figure S1**).

The lysine and aspartate/glutamate residues capable of forming a salt bridge in a triple helix are constrained by sequence. In order to understand these constraints, we refer to the three staggered chains of the triple-helix as leading (staggered towards the N-terminal end), middle or trailing (staggered towards the C-terminal end). In this scheme, the *i*^*th*^ lysine of the leading and middle chain form salt bridges with *i+2* aspartate or glutamate of the middle and trailing chain, respectively (**figure 2A**). The remaining lysine on the trailing chain can also form a salt bridge with the aspartate or glutamate occupying the *i+5* position in the leading chain. Salt bridges that conform to the three sequence constraints are identical with respect to the spatial condition for interaction. However, the *i → i+5* salt bridge is formed by residues that may or may not be present within a KGE or KGD triplet. A search of the aligned triple-helical domains of human collagens shown in **supplementary figure S1** for pairs of lysine and aspartate/glutamate residues that satisfy any of the three sequence constraints revealed 1553 salt bridges (**figure 2A and supplementary table S2**). Of these, 21% are *i → i+5* salt bridges that do not originate from the KGE or KGD triplets. This emphasizes the need to search salt bridges in aligned triple-helical sequences rather than relying solely on instances of KGE and KGD triplets.

**Fig. 2.**
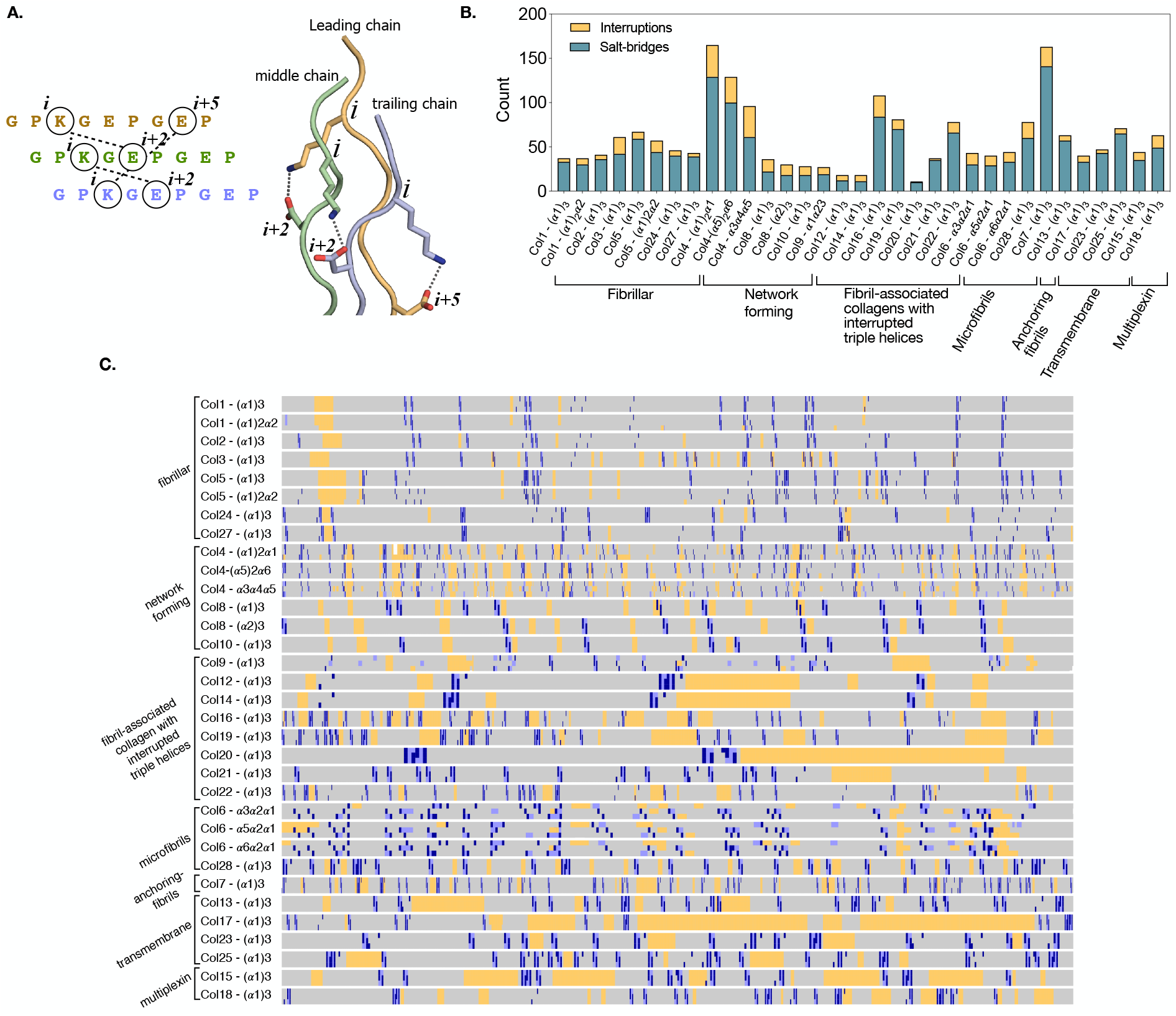
Salt bridges are abundant across all 28 human collagens. (a) Schematic depiction of the sequence constraint for salt bridge formation between the leading (orange), middle (green) and trailing (blue) chains of a triple helix (pdb: 6q3p(*26*)). The number of salt bridges (blue), KGE or KGD triplets (light blue) and interruptions (yellow) in the triple-helical domains of human collagens (b) and a visual representation of their distribution (c) in the triple-helical domains of human collagens.

The observed 1553 salt bridges are distributed across the 28 human collagens with an average of ∼50 in each subtype. A visual inspection of their distribution shown in **figure 2B** suggests that they are present throughout the length of the triple-helical domain. In order to further understand this distribution, we divided the triple-helical domains of aligned collagen sequences into four equal parts and counted the number of salt bridges in each quarter. As shown in **supplementary figure S2**, the C-terminal quarter accounts for ∼50% (757 out of 1553) of all salt bridges. This highly skewed distribution of salt bridges assumes significance in view of the C-terminal nucleation-propagation model currently accepted for the folding of collagen triple helices.

### Misalignment of *α*-chains reduces the number of possible salt bridges

Polypeptides within a triple helix are staggered by one amino acid with respect to each other. This canonical alignment maximizes interchain hydrogen bonds and allows glycines to sequester into the core of the triple helix.(*2*) In theory, the polypeptides can be staggered by an arithmetic series of 1, 4, 7, 10,… amino acids while retaining the tightly-packed triple-helical structure. However, non-canonical alignments of more than 1 amino acid would result in loss of hydrogen bonds at the termini. Thus, triple helices containing non-canonically aligned polypeptides have not been experimentally observed in short collagen peptides. However, given that the triple-helical domains of human collagens contain up to 1000 amino acids and >900 hydrogen bonds, the free energy difference between the canonical and non-canonically aligned triple helices is expected to be small. Thus, if only hydrogen bond and Van der Waals packing are considered, the canonical and non-canonical staggers of collagens would be separated by low energy barriers, resulting in frequent misfolding via misalignment of chains. It remains to be understood how collagens avoid such local folding traps and find the global minimum.

As noted in the previous section, all human collagens contain an average of 50 salt bridges in each collagen subtype. We find that the number of possible salt bridges decreases dramatically upon intentionally misaligning either the middle or the trailing chain by just 4 residues (**figure 3 and supplementary table S2)**. The average loss of salt bridges upon misalignment of the middle and trailing chain by 4 residues is 40 and 30%, respectively. As a representative example, the α3α4α5 heterotimer of collagen type IV loses 8% of salt bridges, while homotrimeric collagen type X loses 94% of salt bridges upon misalignment of the middle chain. The loss in salt bridges is expected to increase the free energy gap between the native and the competing misfolded states, thus ensuring that only the state with the canonical alignment is populated. Importantly, given that 50% of all salt bridges are concentrated in the C-terminal quarter of the triple-helical domains, the decision for correct alignment is likely made early during the propagation phase.

**Fig. 3.**
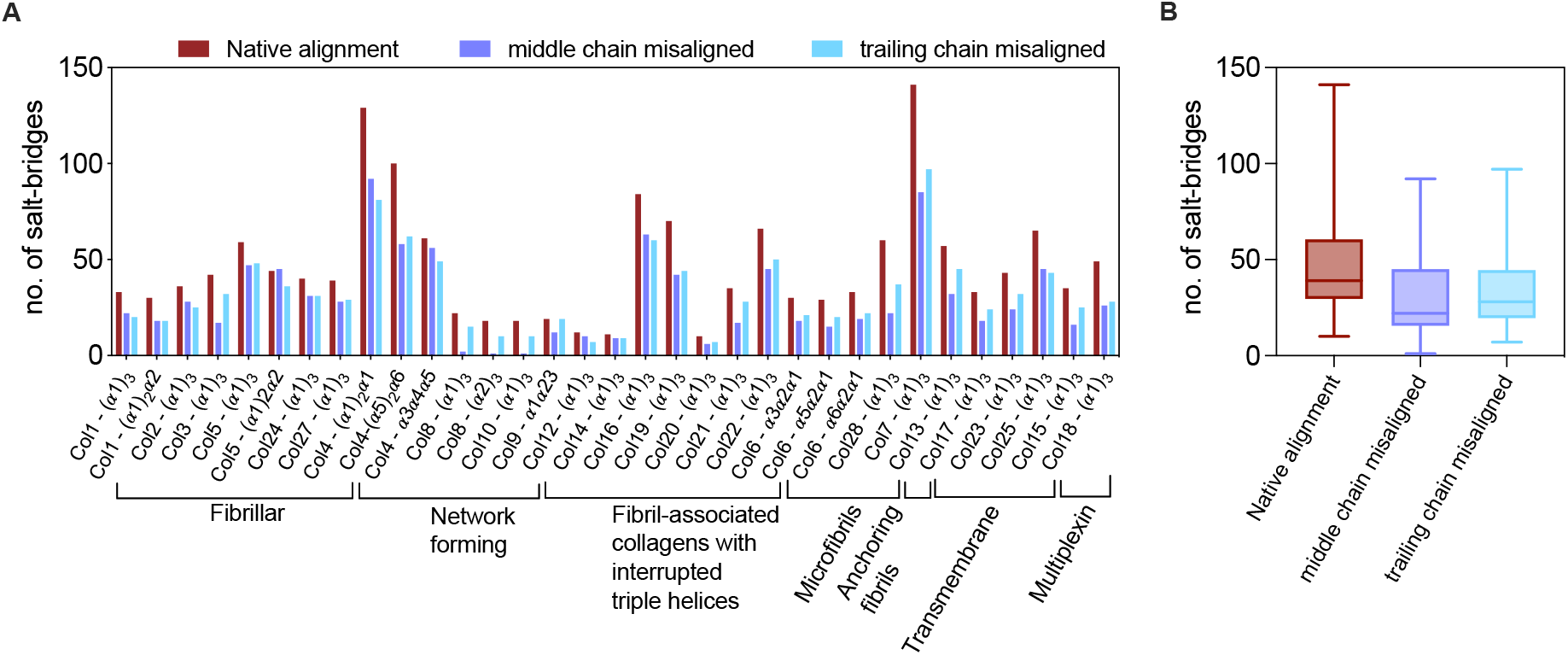
Misalignment of *α* chains reduces the number of salt bridges. Number of salt bridges observed in the correct native alignment (red) and when the middle chain (purple) or trailing chain (cyan) are incorrectly aligned with a four residue offset (A). The box and whiskers plot in (B) shows the aggregate loss in the number of salt bridges upon misalignment.

### Collagens with more interruptions also contain more salt bridges

We find that the number of salt bridges in each collagen subtype is positively correlated to both the number of interruptions and the length of the triple helical domain (**supplementary figure S3**). In physical terms, collagens with more interruptions or longer triple helical domains also contain a greater number of salt bridges. As a representative example, the (α1)_2_α2 heterotrimer of collagen type I with 4 interruptions contains 35 salt bridges but that of collagen type IV with 23 interruptions contains 129 salt bridges. Similarly, collagen type XX with the shortest triple helical domain (141 Gly-Xaa-Yaa triplets) among all collagens contains only 10 salt bridges while collagen type VII with the longest triple-helical domain (1380 Gly-Xaa-Yaa triplets) contains a staggering 141 salt bridges. While the number of salt bridges is strongly correlated to both the number of interruptions (Pearson correlation = 0.69, *P*_*two-tailed*_ < 0.0001, *α*=0.05) and triplets (Pearson correlation = 0.68, *P*_*two-tailed*_ < 0.0001, *α*=0.05), a weaker correlation is observed between the number of triplets and interruptions (Pearson correlation = 0.53, *P*_*two-tailed*_ = 0.0014, *α*=0.05). Interruptions disrupt the triple-helical structure(*27*) causing delayed folding and decreased overall stability(*28*). Similarly, collagens with longer triple helical domains are expected to experience greater entropic disruption of folding during propagation phase. We hypothesize that the increased abundance of salt bridges likely compensates for the disruptive effects of interruptions and also stabilize longer triple-helical domains.

### Human collagen interruptions are flanked by salt bridge”knots”

Of the 394 interruptions identified in the aligned triple-helical domains of human collagens, 58% contain salt bridges on the N- or C-termini or both (**supplementary figure S4**). Close inspection suggests that the salt bridges seldom appear alone. We find that analogous to the cysteine knots in collagens(*29*), where two or more pairs of cysteine residues form disulfide bridges covalently linking all three chains, pairs of two or more interacting lysine and aspartate/glutamate residues also link all three chains. We call these salt bridge knots. As shown in **Figure 4**, we identify knots containing upto eight salt bridges within the sequence space of four triplets.

**Fig. 4.**
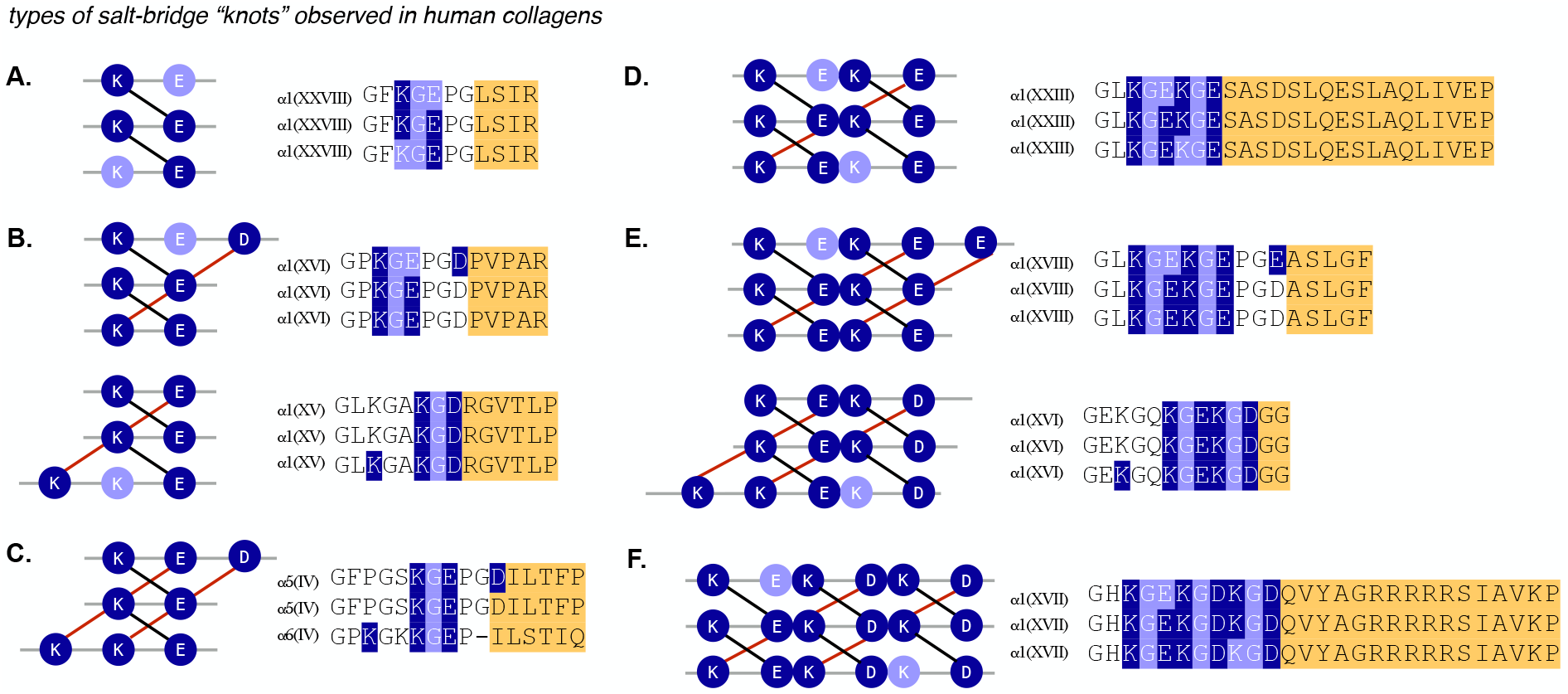
Salt bridges knots flank the interruptions sites in human collagens. Representative examples of a two to six (a-e) and eight (f) salt bridge knot observed in human collagens. Interacting pairs of lysine and aspartate/glutamate residues are shown in dark blue, the KGE and KGD triplets are shown in light purple and interruptions are shown in yellow. The interchain *i → i+2* salt bridges are shown in black line while the *i → i+5* salt bridges are shown in red. Although the representative examples show in this figure contain interruptions only towards the C-termini, examples of salt bridge knots on the N-termini also abound (**supplementary figure S4**).

We purposely distinguish the salt bridge knots from complex salt bridges observed in many globular proteins. As originally defined by Musafia et al(*30*), in a complex salt bridge, a cationic or anionic residue simultaneously interacts with two or more charged residues. In contrast, we define salt bridge knots as pairs of interacting cationic and anionic residues. Complex salt bridges cooperatively increase protein stability i.e. the total change in free energy is more than the sum of parts.(*31, 32*) Due to the extended rod-like topology and the sequence constraint for interaction, complex salt bridges are geometrically not feasible in triple-helices. It is plausible that salt bridge knots in collagens have evolved as an alternative with a role energetically similar to complex salt bridges in globular proteins. Importantly, their predominance close to the structurally disruptive interruptions suggests a key role in compensating for the loss in stability and folding.

### KGE and KGD triplets form geometrically specific salt bridges

Salt bridges can be stabilizing or destabilizing(*33, 34*), depending on the protein context. In a remarkable work, Kumar et al have shown that the relative geometry of interacting cationic (ammonium or guanidinium) and anionic (carboxylate) head group determines whether or not a salt bridge is stabilizing(*35*). This suggests that the stability conferred by a salt bridge is intimiately linked to its geometry. Thus, we analyzed the geometry of salt bridges in the published crystal and solution structures of collagen triple helices using a parameterization previously developed for those in globular protein(*36*). In this parameterization, the geometry of lysine-aspartate/glutamate salt bridges is defined by the angle between carboxylate oxygen and the Cε-Nζ atoms of lysine and the dihedral angle between the carboxylate oxygen and Cδ-Cε-Nζ atoms of lysine. The carboxylate group can adopt one of the three staggered configurations with respect to the tetrahedral ammonium group; gauche plus (*g+*), trans (*t*) and gauche minus (*g-*). Salt bridges in globular proteins are found to favour *g+* or *g-* configurations. Additionally, the carboxylates of glutamate or aspartate can accept hydrogen bond via either of the two nonbonded lone pairs of electrons called the syn and anti. Syn carboxylic acid is a weaker acid than anti and thus its conjugate base is more basic.(*37*) Despite the increased basicity, lysine-aspartate/glutamate salt bridges in globular proteins predominantly form hydrogen bonds via the anti lone pair.

Molecular structures containing lysine-glutamate interactions in collagen triple helices are not available. Thus, in order to understand the geometric preference of salt bridges in collagens, we relied on the published structures containing lysine-aspartate salt bridges. As shown in **supplementary figure S5 and supplementary data 3**, the dihedral angle Cδ-Cε-Nζ---O_carboxylate_ and angle Cε-Nζ---O_carboxylate_ of salt bridges cluster around 180º and 90º, respectively. This suggests an overwhelming preference for the *trans* configuration, in contrast to the *g+*/*g-* preference in globular proteins. The geometric specificity also manifests in whether the syn or anti lone pair interacts with the ammonium moiety. We parameterized this using the angle Oδ2-Oδ2---Nζ_lysine_. Angles between 0 and 120º were classified as *syn* and those above 120º as *anti*. As shown in **supplementary data 3**, the majority of the interactions between the carboxylate and the ammonium group are via the more basic syn lone pair, again in contrast to globular proteins. Thus, our analysis suggests that KGD and KGE triplets also foster geometrically specific salt bridges but, in contrast to globular proteins, these salt bridges strongly favour trans configuration and nearly always interact via the more basic syn lone pair.

### Salt bridges increase kinetic stability of collagens

The anomalously high frequency of KGE and KGD triplets has previously been rationalized based on their ability to foster salt bridges, which presumably increases the thermodynamic stability of collagens(*21*). However, it is well-established that triple-helical peptides containing an OGP triplet unfold at higher temperature than that containing either KGE or KGD.(*23*) We also observe this here (**table 1**). In view of this, the KGE or KGD triplets do not offer any stability advantage compared to an OGP triplet. Thus, the evolutionary implications of retaining KGE/KGD triplets at a frequency comparable to OGP is not clear.

**Table 1.**
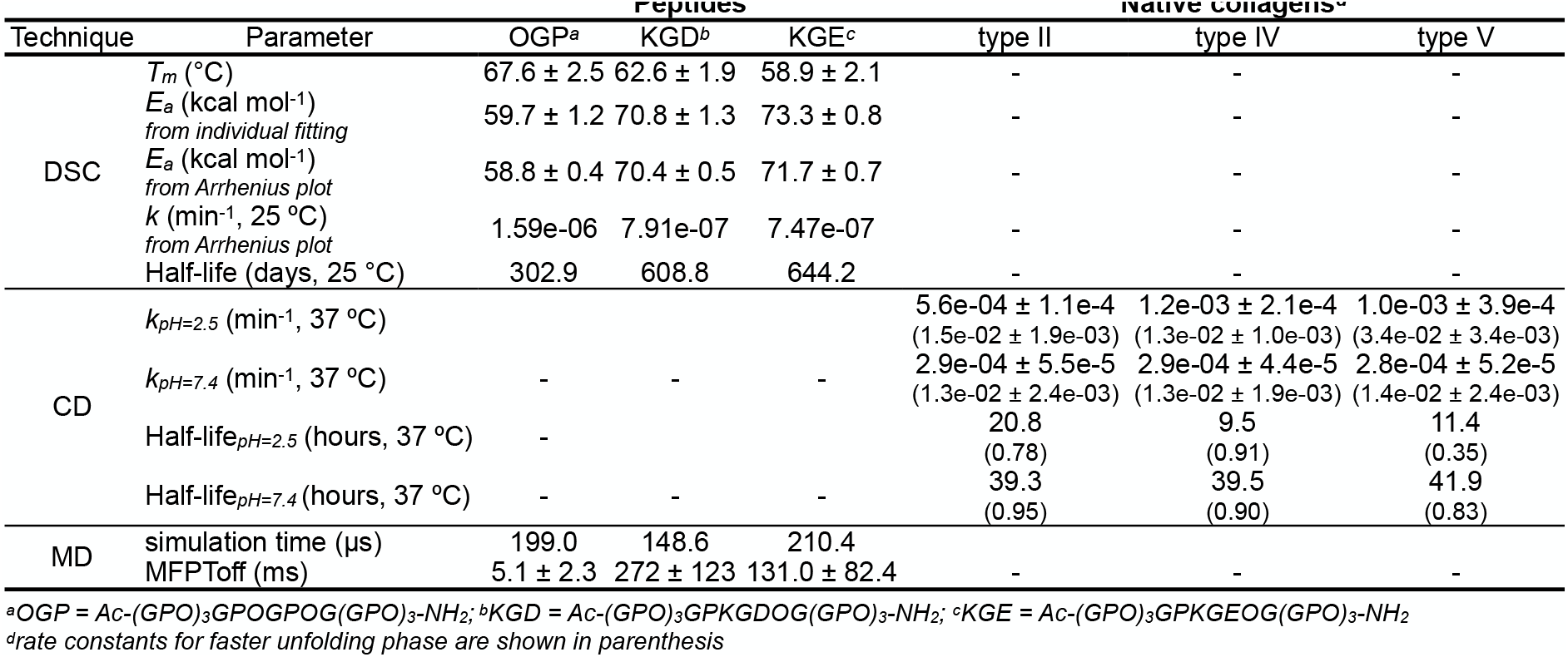
Stability and kinetic parameters for the unfolding of model collagen peptides and native collagens.

Generally, salt bridges increase the thermodynamic stability of proteins.(*35, 38, 39*) However, those with limited(*40, 41*) or even destabilizing effect(*33, 42*) on stability also abound. The destabilization arises due to unfavourable desolvation energy of the interacting electrostatic charges competing against favourable Coulombic attraction. Given the evolutionary pressure to retain even destabilizing salt bridges, it has been postulated that they modulate folding kinetics rather than influence thermodynamic stability.(*38, 43*) This has led to a general appreciation of the role of salt bridges in kinetics irrespective of the thermodynamic component. For example, a surface arginine-glutamate salt bridge in staphylococcal nuclease hinders denaturation by raising the kinetic barrier to unfolding by ∼ 7 kcal/mol.(*44*) Importantly, a geometrically optimized surface salt bridges slows the unfolding rate of *α*-helical peptides while unfavourable geometry has the opposite effect.(*45, 46*) Given that KGE and KGD form geometrically specific salt bridges but do not offer any thermodynamic advantage compared to the OGP triplet, we also suspected a kinetic role.

Due to their trimeric nature, the rate of collagen nucleation is concentration dependent while also limited by the slow rate of proline cis-trans isomerization (*k*_cis*→*trans_ ∼ 0.1 s^-1^).(*47*) Consequently, deconvoluting the effect of salt bridges on refolding rate from the rate-limiting proline isomerization is challenging. Thus, we investigated the unfolding kinetics of collagen peptides containing KGE or KGD triplets via circular dichroism (CD), differential scanning calorimetry (DSC) and molecular dynamic (MD) simulations.

Salt bridge interactions include electrostatic and H-bond components. At pH lower than the pKa of glutamate and aspartate sidechains, the carboxylates are protonated abrogating the electrostatic component. Thus, we monitored the isothermal unfolding of KGE, KGD and OGP peptides at acidic and neutral pH to understand how salt bridges contribute to kinetics (see methods). At pH 2.5, the KGE and KGD peptides unfold to ∼80% of the initial value after incubation at 37 ºC for 8 hours, but no significant change in signal is observed at neutral pH (**figure 5A)**. Importantly, the signal for the OGP peptide is largely independent of pH. We further monitored unfolding kinetics of KGE and KGD peptides. Unfolding of both peptides at neutral pH is extremely slow with less than 10% loss in signal over 8 hours (**figure 5C**). However, at pH 2.5, both peptides unfold quickly with a half-life of 1.6 and 3.5 hours (**table 1**). This pH dependence of unfolding half-life suggest that the increased kinetic stability of KGE and KGD triple helices at neutral pH is primarily due to salt bridge interactions.

**Fig. 5.**
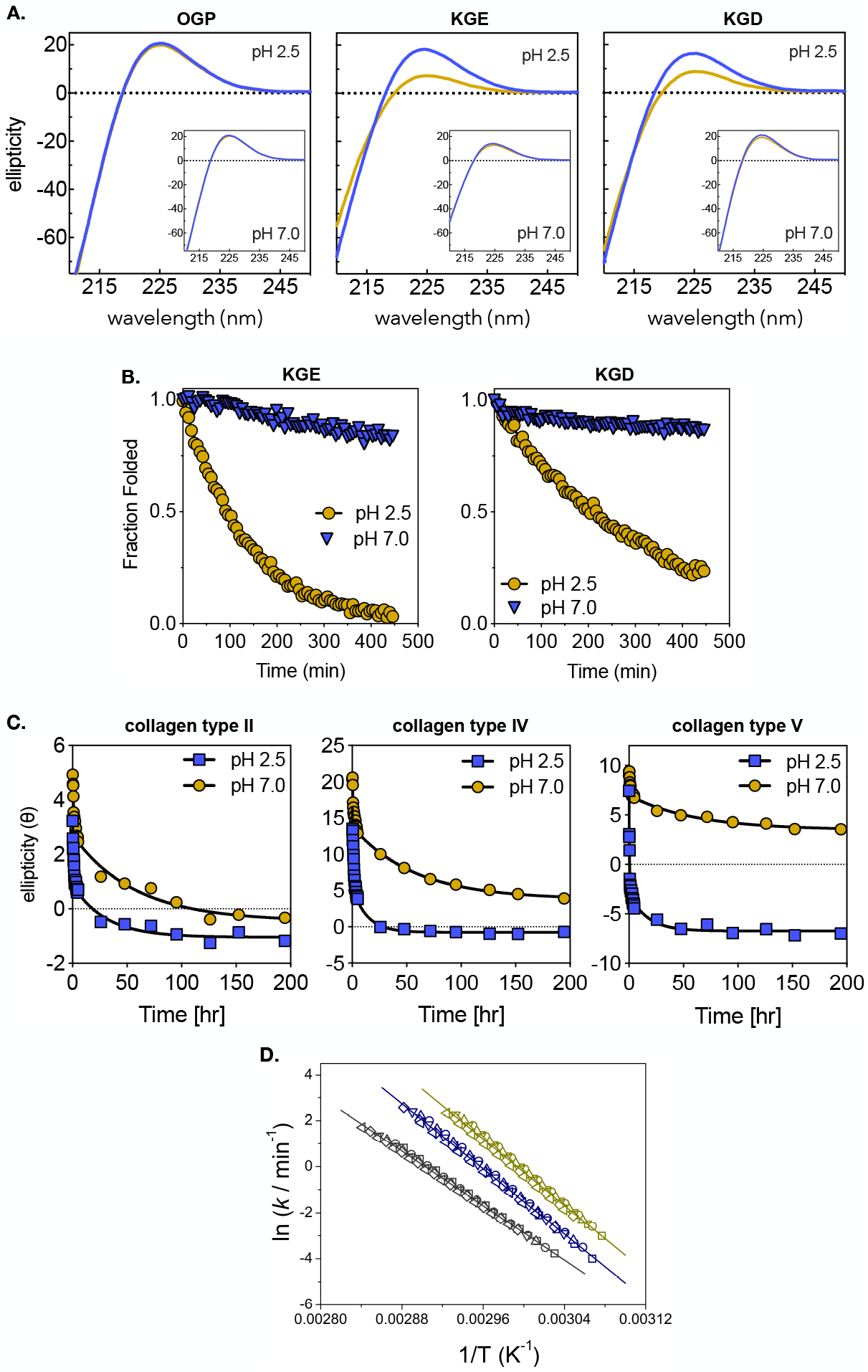
Salt bridges decrease the unfolding rate of collagens. (A) Decrease in the characteristic polyproline type II elipticity signal of OGP, KGE and KGD peptides after isothermal incubation at 37 °C for 8 hours as measured by CD spectroscopy (A), kinetics of unfolding of KGE and KGD peptides (B) and collagens type II, IV and V at pH 2.5 (blue) and 7.0 (orange) (C). Residuals of the monoexponential fit of collagen peptides and biexponential fit of native collagens are shown **in supplementary figure S6**. (D) Kinetic stability comparison of OGP (gray), KGE (blue) and KGD (yellow) peptides analyzed by an Arrhenius plot derived from Differential Scanning Calorimetry experiments performed at different scan rates. The line represents the best fit to the Arrhenius equation. Kinetic parameters are shown in **table 1**. DSC endotherms and fitting to a two-state kinetic model are shown in **supplementary figure S7**.

We further investigated the unfolding kinetics of native fibrillar collagens type II and V, which contain single uninterrupted triple-helical domains and collagen type IV that contains a triple-helical domain with 23 interruptions. As shown in **figure 5C**, all three collagen types show pH dependent unfolding kinetics. The unfolding traces could be fit to a biexponential kinetics with a faster and a slower decay phase. The half-life of the faster unfolding phase is largely independent of pH as well as the type of collagen except type V where a two fold difference is observed. In contrast, the half-life of the slower unfolding phase is pH dependent for all collagen subtypes. Collagen type 4 and 5 unfold ∼ 4-fold slower at pH 7.4 than at pH 2.5 while collagen type 2 unfolds two-fold slower, as judged from their half-lives.

Further, we investigated unfolding of peptides KGD and KGE via DSC to estimate kinetic barriers separating the denatured and folded states. The thermal-unfolding endotherms show a temperature shift on melting transition upon varying the scan rate (**supplementary figure S7**). This suggests that a kinetic process controls unfolding. Kinetic control of collagen unfolding has also been observed by others.(*48*) In our case, fitting of the endotherms to a kinetic model provides an estimate of the unfolding activation energy (**methods**). This parameter has been previously correlated with the magnitude of the protein kinetic stabilty(*49, 50*). In comparison to the OGP peptide, the KGD and KGE peptides have a lower thermal stability but considerably higher activation energy (**table 1**), both in the Arrhenius plot (**figure 6A**) and the endotherm fitting (**supplementary figure S8**). These results indicate a clear difference in kinetic stability as inferred by the one order of magnitude change in the kinetic constant (*k*) and consequently the unfolding half-time of the peptides (**table 1)**. To confirm that the observed differences are indeed due to salt bridges interactions, we performed DSC measurements in presence of physiological concentrations of NaCl (**supplementary figure S8)**. While Tm and activation energy are essentially unchanged for OGP in the presence of salt, a destabilizing effect is observed for both KGE and KGD. This salt effect is reflected on considerable changes on Tm and activation energy (**supplementary table S3**), which further confirms the kinetic role of salt bridges in collagen folding.

The trend in kinetic stability of peptides as observed by CD and DSC experiments is also confirmed by molecular dynamics (MD) simulations. Starting from the previously published crystal structures of collagen peptides containing the KGD and KGE(*23*) or OGP(*51*) triplets, we conducted multiple all-atom simulations in parallel, each capable of sampling various conformational landscapes. Specifically, simulations were conducted for systems containing the triplets KGE, KGD, and OGP, alongside a simulation with the KGE triplet at a NaCl concentration of 150 mM (**methods**). In total, we accumulated over 750 μs of aggregated simulated time (**table 1**). To facilitate unfolding processes within a reasonable timescale, all systems were maintained at 400 K. Each individual simulation ran for 100 ns, which prevented the observation of complete unfolding events but allowed us to sample partial fragments of the process. We therefore employed Markov state models (MSMs) to combine the information from the entire ensembles and analyze their collective behavior. MSMs have proven invaluable for characterizing kinetics, thermodynamics, and trajectories of processes like ligand-binding(*52*), folding(*53*), and protein-protein interactions(*54*) with atomic precision. We computed MSMs for all our systems, which allowed us to obtain approximate transition rates between the folded and unfolded states (**methods, supplementary figure S9**). More specifically, in the context of MSMs, we define the mean first passage time (MFPT) as the average time required to transition from one state to another. We computed MFPTs for our simulations, thereby characterizing the average times required for the different systems to reach the unfolded state. As indicated in **table 1**, KGE and KGD triple-helical peptides unfold approximately two orders of magnitude slowler than those containing hydroxyproline or high salt concentration. To summarize, both theoretical and experimental investigations suggest that salt bridges interactions increase the kinetic stability of collagen triple-helical peptides more than those containing only proline and hydroxyproline. This provides a rational for why Nature has retained the KGE and KGD triplets at anomalously high frequency in human collagens.

### Mutations that disrupt salt bridges are associated with disease

Mutation of amino acids critical for protein folding and function causes disease. We investigated if mutations within or in the periphary of salt bridges in collagen also cause disease. It has previously been shown that Gly *→* Ala mutation within a KGD triplet completely unfolds a region located 9 residues upstream towards the N-termini.(*55*) In a separate study, NMR measurements suggest that triplets N-terminal to the mutation site exhibit unfolded monomer-like dynamics.(*56*) This mutation also disrupts a salt bridge located two triplets towards the C-termini.(*57*) NMR measurements also suggest that mutation of glycine induces a large dynamic motion that propagates at least two triplets towards the C-termini.(*56*) In effect, mutation of the charged residues or any glycine in three triplets on the N-termini and two triplets in the opposite direction can potentially disrupt salt bridges. We refer to this region over which a salt bridge disruption can influence folding as the salt bridge footprint. In order to understand if mutations within the salt bridge footprint are associated with disease, we searched the Clinvar(*58*), Leiden Open Variation Database (LOVD)(*59*) and Alport Syndrome Database(*60*) for pathogenic missense mutations due to single nucleotide change. We identified a total of 2294 mutations, of which 565 (∼25%) occur within the salt bridge footprints (**supplementary table S4**). The most dramatic localization of mutations in the salt bridge footprint is observed in the *α*5 (38%) and *α*6 (52%) chains of collagen type IV and the *α*1 (45%) chain of collagen type VII. The association of salt bridge disruption to disease-causing mutations testifies to their biological function. It is anticipated that mutations would lower the kinetic stability of triple helices causing downstream effects such as change in fibrillogenesis, collagen turnover and transport and secretion to the extracelluar matrix but the exact mechanism remains to be investigated.

### Glycine mutations in the salt bridge footprint of collagen type I are associated with lethal phenotypes

Mutations in the triple-helical region of collagen type I cause osteogenesis imperfecta (OI), also known as the brittle bone disease. Based on the mode of inheritance and clinical symptoms, Sillence et al. classified OI into four broad categories; OI1 presents moderate phenotype, OI2 is lethal causing stillbirth or perinatal death, OI3 causes most-severe but non-lethal phenotype while OI4 phenotype varies between 1 and 3.(*61*) A survey of the LOVD database suggests that 192 missense mutations present in the triple-helical domain of col1*α*1 and 164 in col1*α*2 cause lethal (OI2) or severe (OI3) phenotype. Of these, 54 mutations in the *α*1 chain (28%) and 40 mutations in *α*2 chain (24%) are present within the salt bridge footprint. Importantly, 55% (52 of 94) mutations in the salt bridge footprint of the *α*1 and *α*2 chains are lethal.

Previously, mutational hotspots in collagen type I that cause lethal OI have been mapped to two sequence stretches in the *α*1 chain (residue 869-1001 and 1088-1142) and 8 in *α*2 chain (residues 409-454, 541-592, 637-670, 712-727, 784-796, 847-901, 949-997 and 1027-1084)(*62, 63*). These lethal regions are schematically shown in **supplemenatry figure S10**. It has previously been postulated that lethal regions in collagen sequence correlate with the major ligand binding regions of the triple-helical domain.(*64*) Presumably, mutations disrupt the interaction of collagen fibrils to other proteins, resulting in pathology. However, only 8 of the 54 lethal mutations in the *α*1 chain within the salt bridge footprint co-localize with the known lethal region. Our results suggest that in addition to protein-protein interactions, the molecular effects of mutations that potentially disrupt salt bridges could also be consequential in determining the severity of phenotype.

## Discussion

Collagen is an enigmatic protein from a folding perspective. Unlike globular and other coiled-coils proteins, the triple helix lacks a hydrophobic core and is primarily stabilized by interchain hydrogen bonds formed by the repetitive glycines. Amino acids in the Xaa and Yaa positions determine various aspects of folding and stability. Collagen *α*-chains are generally translated with a globular C-terminal pro-domain(*65*) that self-trimerizes and directs assembly of the correct triple-helical stoichiometry while also facilitating rate-limiting nucleation. Importantly, the pro-domains are cleaved before the nascently folded triple helices undergo further self-assembly into fibers, networks and filaments. In a remarkable work, Leikina et al have shown that collagen type I unfolds at body temperature.(*66*) The implication is that the nascent triple helices have unfavourable free energy and would start unfolding as soon as the pro-domains are cleaved. In order to explain how Nature circumvents this folding problem, it has been proposed that the ER-resident chaperone Hsp47 binds unique sites in triple helices. This interaction is believed to minimize local unfolding, which could be a precursor to global unfolding, and also offset the entropic cost for the stepwise propagation of a ∼1000 amino acid long polypeptide. (*67, 68*) However this folding paragdim does not reconcile with the experimental observation that exogenously added Hsp47 does not significantly influence either the stability or the folding rate of collagen type I *in vitro*.(*69*) Several other biochemical observations are also not fully cosistent with the Hsp47-chaperoned folding paradigm.

Hsp47 binding sites are predominantly located towards the N-terminal half of collagens type I, II and III.(*12, 20, 70*) Given the C- to N-terminal propagation, the implication is that Hsp47 binds these collagens after half the triple-helix has already folded. Hsp47 also does not recognize or does so only weakly collagens type XI, XIV, XXII.(*12*) This is also corroborated by our analysis of the distribution of potential Hsp47 recognition sites across the 28 human collagens (**supplementary figure S11**). We find that collagens type VIII, XIII, XIV, XV, XVIII, XIX, XXI and XXV contain less than 3 Hsp47 binding sites even when low affinity sites are considered. Hsp47 chaperoned folding also runs counter to the general experimental observation that native collagens spontaneously refold *in vitro* in its absence. The role of Hsp47 during late stages of collagen folding such as preventing premature lateral aggregation of folded triple-helices(*71, 72*), transporting pro-domain cleaved triple-helices to the ER-Golgi boundary(*73, 74*) and then sorting into large pro-collagen cargos(*75*) for timely secretion into the extracellular matrix is well-established. The experimental observations that Hsp47 ablation does not influence secretion of collagen type VI microfibrils but only its lateral assembly supports this argument.(*12*) We also find that fibrillar collagens, which are composed of laterally assembled triple helices, contain the highest abundance of Hsp47 binding sites (**supplementary figure S11**). Importantly, collagen-like protein domains of considerable length and interruptions are also found in archaea, bacteria, eukarya (excluding human collagen orthologues) and viruses. To the best of our knowledge, there is currently no evidence for the presence of Hsp47-like proteins in organisms from these groups. Thus, how collagens and collagen-like proteins in these groups fold and then remain folded is an open question.

Salt bridges help globular proteins avoid unproductive folding pathways by stabilizing productive folding intermediates.(*76, 77*) Here, we propose that the correct folding of collagen and collagen-like proteins is also a consequence of the geometric specificity of salt bridges and their ability to decrease the unfolding rate of triple helices. We find that the 28 human collagens contain 1553 lysine – aspartate/glutamate salt bridges with an average of 50 in each collagen subtype. We also find that the interaction between lysine and aspartate/glutamate residues is geometrically specific and that they increase kinetic stability of model triple-helical peptides as well as native collagens. Of the 1553 salt bridges, 50% are localized in the C-terminal quarter of the triple-helical domains. This highly skewed concentration of salt bridges in the C-terminal quarter along with their ability to increase kinetic stability suggests that these act as electrostatic clamps to prevent local unfolding and, in the process, allow triple helices to reach a critical length from which the rest of the propagation can occur. Our observation that collagens with longer triple-helical domains or greater number of structurally and energetically disruptive non-collagenous interruptions also have more salt bridges further corroborates this hypothesis. Additionally, the observation that interruptions are frequently flanked by multiple closely spaced salt bridges, which we call salt bridge knots, adds further weight to it. CD and DSC experiments suggest that the increased kinetic stability is a result of a raised kinetic barrier to unfolding. Importantly, this kinetic barrier is lower for the triple helices containing only proline and hydroxyproline than those containing salt bridges. This provides a rational for why KGE and KGD triplets are anomalously frequent despite a thermal stability contribution comparable to OGP. The salt bridge assisted folding mechanism circumvents the need for a chaperone during the triple-helical propagation phase. This folding paradigm likely also explains how collagen-like proteins from archaea, bacteria, eukarya and viruses, with their stupendously long triple-helical domains and many interruptions, are able to successfully fold and remain folded. The wider implication is that evolution has converged on a similar mechanism to stabilize triple helices across the the three domains of life as well as viruses.

Our results have additional important consequences for collagen folding. The three polypeptides of a collagen triple helices are staggered by one amino acid with respect to each other to optimize interchain hydrogen-bonds and van der Waals packing. Given the enormous length of triple-helical domains and the diversity of interruptions, incorrect staggers of more than one amino acid are plausible. How collagens avoid such misfolded states is currently not understood. Our observation that incorrect stagger of *α*-chains reduces the number of salt bridges by an average of 50% across all collagen subtypes suggests a mechanism for how collagens might ensure correct stagger. Most revealingly, we find that mutation of glycine residue within or in the periphery of a salt bridge are associated with heritable diseases across several collagen types while such mutations in collagen type I frequently result in lethal phenotype. Previously, efforts to rationalize why mutation of some glycine residues in collagen type I cause a more severe phenotype than others using loss in thermal stability showed a poor correlation.(*78*) We can extrapolate from our results that the phenotypic severity is likely correlated to the loss in kinetic rather than thermodynamic stability. The structural consequence of mutation in a salt bridge footprint, how it affects folding kinetics, stability, fibrillogenesis and ultimately transportation into the extracellular matrix and why it overwhelmingly results in lethal phenotype is not clear to us. However, these questions gain significance in light of our observation that mutations in salt bridges generally result in lethal to severe phenotype, even when present outside the aformentioned lethal regions. Given the effect of salt bridges on collagen kinetic stability as demonstrated here, a systematic study to explore correlation between loss in kinetic stabilty and the phenotypic severity due to a mutation is desirable in future.

## Acknowledgements

Part of the work was funded by the Newton International Alumni Funding (2018-2013) awarded to A.A.J jointly by the Royal Society, the British Academy and the Academy of Medical Sciences. J.D.M was partly funded by the French National Agency (CARTEGRIN ANR21-CE19-0017). The authors would like to thank Prof Birte Höcker for access to DSC instrument. The authors gratefully acknowledge the scientific support and HPC resources provided by the Erlangen National High Performance Computing Center (NHR@FAU) of the Friedrich-Alexander-Universität Erlangen-Nürnberg (FAU) under the NHR project b114cb (UID 210235). NHR funding is provided by federal and Bavarian state authorities. NHR@FAU hardware is partially funded by the German Research Foundation (DFG) – 440719683. We also thank Prof Thomas Scheibel for access to the CD spectrometer. The authors specially thank Prof Richard Farndale for insightful discussion and help with the peptide synthesis.

## Contributions

A.A.J. and J.D.M conceived the project and developed the experimental proposal. A.A.J performed sequence analysis, made deductions based on these, made the figures, performed CD experiments and also wrote the manuscript, SRR performed DSC experiments and the related data analysis, NF performed the MD simulations and analyzed the data, SD clustered the collagen sequences using CD-HIT, VS performed the CD kinetic unfolding experiments of the peptides.

## Methods

### Identification of the collagen domains and interruptions

UniProtKB database was queried on 16.03.2023 for proteins containing the collagen or collagen-like sequences by applying taxonomic filters (archaea, bacteria, eukarya or virus) while filtering for sequences annotated by InterPro as containing collagen triple-helical repeats (ipr008160). The search terms used were “(taxonomy_id:2759) AND (xref:interpro-ipr008160)” for eukarya, “(taxonomy_id: 10239) AND (xref:interpro-ipr008160)” for viruses, “(taxonomy_id:2157) AND (xref:interpro-ipr008160)” for archae, “(taxonomy_id:2) AND (xref:interpro-ipr008160)” for eubacteria. The 44 *α* chains of human collagen sequences were also obtained from UniProtKB by applying taxonomic filter 9606 and searching for gene names such as COL1A1, COL2A1 etc. A total of 59226 collagen sequences were downloaded from UniProtKB, which were split into smaller datasets based on the phyla they were recovered from. CD-HIT(*79*), a clustering algorithm to produce non-redundant (nr) dataset was used cluster these collagen domains from each taxa. The sequences were clustered at 70% identity to remove most of the proteins with high sequence similarity and retain enough diversity. After removal of duplicate sequences, collagen or collagen-like domains in the taxa-specific fasta files were identified by searching for contiguous repeats of 6 or more Yaa-Gly-Xaa triplets. The total observed instance of each Yaa-Gly-Xaa triplet in each collagenome was counted and their relative abundance determined using the **eqn. 1** below,

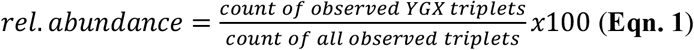

The predicted distribution of the triplets was calculated based on the statistical model described previously.(*21*) Given the 20 canonical amino acids, a total of 361 Yaa-Xaa residue pair combinations (excluding glycine) are theoretically possible in the Yaa-Gly-Xaa triplet. We counted the observed instance of the different triplets and calculated Z-scores from the numerical difference in the observed and predicted count of triplets using the Z-score function in R package.(*80*) Motifs with Z-scores ± 3 were considered outliers and thus anomalously frequent.

Interruptions in the human collagen domains and the collagen-like proteins in the archaea, bacteria, eukarya and viruses were identified essential as described previously(*81*). Briefly, interruptions were defined via their sequence length. For example, a deletion of one amino acid from the triplet repeat sequence was denoted as G1G and a deletion of three or more amino acids is denoted G3G to G(Z)_n_G, where Z denotes interruption and subscript *n* its length up to 15 amino acids. The interruptions along with 7 amino acids on the N- and C-termini were extracted from the human and taxa-specific fasta files containing the collagen and collagen-like domains and Z-scores were calculated as described in the previous paragraph.

### Peptide synthesis

The Ac-(GPO)_3_GPKGEO(GPO)_3_-NH_2_ (KGE), Ac-(GPO)_3_GPKGDO(GPO)_3_-NH_2_ (KGD) and the Ac-(GPO)_8_-NH_2_ (OGP) peptides were synthesized on a TentaGel Rink Amide MBHA resin at a scale of 0.1 mmol using standarad Fmoc-based solid state peptide synthesis chemistry on a CEM microwave peptide synthesizer. The peptides were acetylated at the N-terminal and amidated at the C-terminal and cleaved from the resin using 95:2.5:2.5 volumetric mixture of trifluoroacetic acid (TFA), triisopropylsilane and miliQ-H_2_O. Cleaved peptides were precipitated from TFA solution using dry-ice-cold diethyl ether, filtered under vacuum, re-dissolved in 95:5 water:acetonitrile (0.1% TFA), freeze-dried and stored at -20 ºC. The cleaved peptides were purified using reverse phase HPLC using a gradient of acetonitrile in water with 0.1% TFA. Multiple fraction corresponding to the main eluting peak were collected and analyzed by matrix-assisted laser desorption ionization mass spectroscopy. Fractions containing the desired peptide were pooled, flash frozen in liquid nitrogen and lyophilized to obtain pure peptides used in all further experiments.

### Determination of unfolding rates via Circular Dichroism

The unfolding of peptides were determined to investigate the effect of electrostatic interactions on their kinetic stability. Peptides OGP, KGE or KGD at pH 2.5 (aqueous HCl) or 7.0 (10 mM sodium phosphate buffer) at 0.4 mg/ml total peptide concentration were equilibrated in an Eppendorf Thermomixer at 37 ºC and the characteristic CD maximum of polyproline type II helices at 225 nm monitored as a function of time until at least 80% of the initial signal has decayed. For measuring the spectra, the peptide solutions were transferred to a quartz cuvette (pathlength = 1mm) pre-equilibrated at 37 ºC via a Peltier temperature controller attached to a Jasco-815 CD spectropolarimeter and CD spectrographs recorded with a data pitch of 5 s and response time of 4 s at predetermined intervals. The average deadtime between transfer of the solutions to the cuvette and recording of the spectra was 3 s. All spectra were recorded three times with three independent samples prepared from a common stock solution. The ellipticity of the peptide solutions was plotted as a function of wavelength without further data processing.

The unfolding rates were measured at 37 ºC to match the experimental condition for monitoring unfolding of native collages described later. 0.4 mg/ml peptide solutions at pH 2.5 or 7.0 were incubated at 37 ºC in an Eppendorf Thermomixer and the change in CD signal at 225 nm was monitored as a function of time. For each data point, the signal at 225 nm was averaged for 2 minutes. The samples were stored at 37 ºC in an eppendorf tube between measurements. Unfolding spectra were recorded thrice with samples independently prepared from a common stock solution. The raw data for samples were fitted to a one-phase decay model in GraphPad Prism according to **eqn. 2** below and the data normalized using **eqn. 3**.

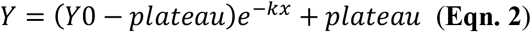

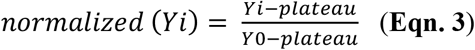

In order to understand how electrostatic interactions influence kinetic stability of native collagens, we monitored unfolding of kinetics of native collagens type II (bovine tracheal cartilage; Sigma-Aldrich C1188), type IV (human placenta; Sigma-Aldrich C5533) and type V (human placenta; Sigma-Aldrich C3657) in aqueous buffer at pH 7.4 (200 mM sodium phosphate, pH 7.4 containing 0.5 M glycerol to prevent fibrillogenesis) and aqueous HCl at pH 2.5. 1 mg of each collagen was suspended in 1 ml of neutral or low pH buffer precooled in a ice bath and vortexed for 10 minutes. After orbital shaking overnight in a cold room, the suspensions were centrifuged at 5000 rpm for 10 minutes at 5 ºC and the supernatants were used for further experiments. In a typical kinetic experiment, the native collagen samples stored at 5 ºC were equilibrated to 37 ºC for 10 minutes in an Eppendorf Thermomixer and then transferred to a quartz cuvette (pathlength=1mm) also maintained at 37 ºC in the spectrpolarimeter. The remaining experimental conditions for data acquisition were identical to those used for the collagen triple-helical peptides.

### Determination of activation energy via DSC

Differential Scanning Calorimetry (DSC) measurements were collected in a VP-Capillary DSC (Malvern Panalytical). Samples were assayed at 0.4 mg/mL in buffer 10 mM sodium phosphate pH 7.0, and varying the scan rate from 0.5 °C/min to 3 °C/min. For experiments to determine the effect of salt on the kinetic stability, thermal unfolding was measured in buffer 10 mM sodium phosphate pH 7.0 and 154 mM NaCl. All scans were collected after exhaustive dialysis and buffer degassing. In all cases, proper instrument equilibration was reached by running at least 2 buffer-buffer scans before sample-buffer experiments. The last buffer-buffer scan was then used to subtract the signal from each peptide-buffer scan in order to perform all thermodynamic analysis.

Calorimetric transitions were adequately described by the two-state kinetics model (N →F) where N is the native peptide and F is the final state.(*82*) The kinetic conversion from N to F is described by a first-order rate constant (k) changing with temperature according to the Arrhenius equation (**eqn. 4**):

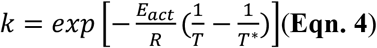

where *T*^*\**^ is the temperature at which the kinetic constant *k*= 1 min^−1^ and *E*_*act*_ is the activation energy between the native and the transition states that describes the unfolding process.(*50*) Here, *E*_*act*_ was used to compare the kinetic stability among peptides. Then, the apparent heat capacity which describes the endotherm is given by (**eqn. 5**):

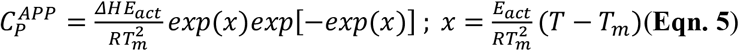

where *T* is the temperature and Δ*H* is the unfolding enthalpy. The *E*_*act*_ was also obtained from the slope of Arrhenius plots, i.e. ln *k* vs. 1/T as described before.(*83*)

### Determination of unfolding times via MD simulations

The simulation systems were prepared taking the X-ray structures as protein templates (PDB code 3T4F and 3U29 for KGE and KGD, respectively). The molecules were modelled to 15 residues, with four residues at the N-terminus before the triplet and 8 after. Systems were solvated and ions were added to neutralization, except for the KGE system which was modeled at high concentration (150 mM NaCl). The systems had ca. 34,000 atoms and ran at 400K. The simulations ran as in our previous work, following an adaptive sampling scheme after the equilibration protocol(*52*). The Markov model analysis ran for all the systems using a lag time of 40, 25, 25 and 40 ns for systems KGE, KGD, GPO and KGE at high concentration, respectively. The MSMs were computed with the software HTMD(*84*). To provide statistics, we bootstrap the data, remove 20% of the trajectories at each iteraction, and re-compute the MSM each time.

## References

1. S. Ricard-Blum, The Collagen Family. Cold Spring Harb. Perspect. Biol. 3, 1–19 (2011).

2. J. Bella, M. Eaton, B. Brodsky, H. M. Berman, Crystal and molecular structure, of a collagen-like peptide at 1.9 Å resolution. Science (80-.). 266, 75–81 (1994).

3. S. Park, T. E. Klein, V. S. Pande, Folding and misfolding of the collagen triple helix: Markov analysis of molecular dynamics simulations. Biophys. J. 93, 4108–15 (2007).

4. K. Beck et al., Destabilization of osteogenesis imperfecta collagen-like model peptides correlates with the identity of the residue replacing glycine. Proc. Natl. Acad. Sci. U. S. A. 97, 4273–4278 (2000).

5. J. Myllyharju, K. I. Kivirikko, Collagens and collagen-related diseases. Ann. Med. 33, 7–21 (2001).

6. J. Engel, D. J. Prockop, The zipper-like folding of collagen triple helices and the effects of mutations that disrupt the zipper. Annu. Rev. Biophys. Biophys. Chem. 20, 137–152 (1991).

7. X. Sun et al., A Natural Interruption Displays Higher Global Stability and Local Conformational Flexibility than a Similar Gly Mutation Sequence in Collagen Mimic Peptides. Biochemistry. 54, 6106–6113 (2015).

8. M. Kurkinen, A. Taylor, J. I. Garrels, B. L. M. Hogan, Cell surface-associated proteins which bind native type IV collagen or gelatin. J. Biol. Chem. 259, 5915–5922 (1984).

9. M. Yagi-Utsumi et al., NMR and Mutational Identification of the Collagen-Binding Site of the Chaperone Hsp47. PLoS One. 7, 5–10 (2012).

10. C. Widmera et al., Molecular basis for the action of the collagen-specific chaperone Hsp47/SERPINH1 and its structure-specific client recognition. Proc. Natl. Acad. Sci. 109, 13243–13247 (2012).

11. T. Koide et al., Specific recognition of the collagen triple helix by chaperone HSP47: II. The HSP47-binding structural motif in collagens and related proteins. J. Biol. Chem. 281, 11177–11185 (2006).

12. A. Köhler et al., New specific HSP47 functions in collagen subfamily chaperoning. FASEB J. 34, 12040–12052 (2020).

13. A. A. Jalan, K. A. Jochim, J. D. Hartgerink, Rational design of a non-canonical “sticky-ended” collagen triple helix. J. Am. Chem. Soc. 136, 7535–7538 (2014).

14. T. A. Linden, N. King, bioRxiv, in press (available at https://www.biorxiv.org/content/10.1101/2021.10.08.463732v1%0Ahttps://www.biorxiv.org/content/10.1101/2021.10.08.463732v1.abstract).

15. Y. Qiu, C. Zhai, L. Chen, X. Liu, J. Yeo, Current Insights on the Diverse Structures and Functions in Bacterial Collagen-like Proteins. ACS Biomater. Sci. Eng. (2021), doi:10.1021/acsbiomaterials.1c00018.

16. M. Rasmussen, M. Jacobsson, L. Björck, Genome-based identification and analysis of collagen-related structural motifs in bacterial and viral proteins. J. Biol. Chem. 278, 32313–6 (2003).

17. M. Zairi, A. C. Stiege, N. Nhiri, E. Jacquet, P. Tavares, The collagen-like protein gp12 is a temperature-dependent reversible binder of SPP1 viral capsids. J. Biol. Chem. 289, 27169–27181 (2014).

18. N. Ghosh et al., Collagen-like proteins in pathogenic E. coli strains. PLoS One. 7 (2012), doi:10.1371/journal.pone.0037872.

19. Y. Xu, D. R. Keene, J. M. Bujnicki, M. Höök, S. Lukomski, Streptococcal Scl1 and Scl2 proteins form collagen-like triple helices. J. Biol. Chem. 277, 27312–8 (2002).

20. H. Cai et al., Identification of HSP47 Binding Site on Native Collagen and Its Implications for the Development of HSP47 Inhibitors. Biomolecules. 11, 983 (2021).

21. T. Stability, A. V. Persikov, J. A. M. M. Ramshaw, A. Kirkpatrick, B. Brodsky, Electrostatic interactions involving lysine make major contributions to collagen triple-helix stability. Biochemistry. 44, 1414–1422 (2005).

22. A. V. Persikov, J. A. M. Ramshaw, A. Kirkpatrick, B. Brodsky, Amino acid propensities for the collagen triple-helix. Biochemistry. 39, 14960–14967 (2000).

23. J. A. Fallas, J. Dong, Y. J. Tao, J. D. Hartgerink, Structural Insights into Charge Pair Interactions in Triple Helical Collagen-like Proteins. J. Biol. Chem. 287, 8039–47 (2012).

24. J. M. Daubenspeck et al., Novel oligosaccharide side chains of the collagen-like region of BclA, the major glycoprotein of the Bacillus anthracis exosporium. J. Biol. Chem. 279, 30945–30953 (2004).

25. K. Mann et al., Glycosylated threonine but not 4-hydroxyproline dominates the triple helix stabilizing positions in the sequence of a hydrothermal vent worm cuticle collagen. J. Mol. Biol. 261, 255–66 (1996).

26. A. A. Jalan et al., Chain alignment of collagen I deciphered using computationally designed heterotrimers. Nat. Chem. Biol. 16, 423–429 (2020).

27. T. Xu, C. Z. Zhou, J. Xiao, J. Liu, Unique Conformation in a Natural Interruption Sequence of Type XIX Collagen Revealed by Its High-Resolution Crystal Structure. Biochemistry. 57, 1087–1095 (2018).

28. E. S. Hwang, B. Brodsky, Folding delay and structural perturbations caused by type IV collagen natural interruptions and nearby Gly missense mutations. J. Biol. Chem. 287, 4368–4375 (2012).

29. A. S. DiChiara et al., A cysteine-based molecular code informs collagen C-propeptide assembly. Nat. Commun. 9 (2018), doi:10.1038/s41467-018-06185-2.

30. B. Musafia, V. Buchner, D. Arad, Complex salt bridges in proteins: Statistical analysis of structure and function. J. Mol. Biol. 254, 761–770 (1995).

31. C. a Olson, E. J. Spek, Z. Shi, a Vologodskii, N. R. Kallenbach, Cooperative helix stabilization by complex Arg-Glu salt bridges. Proteins. 44, 123–32 (2001).

32. A. G. Gvritishvili, A. V. Gribenko, G. I. Makhatadze, Cooperativity of complex salt bridges. Protein Sci. 17, 1285–1290 (2008).

33. P. Phelan et al., Salt bridges destabilize a leucine zipper designed for maximized ion pairing between helices. Biochemistry. 41, 2998–3008 (2002).

34. Z. S. Hendsch, B. Tidor, Do salt bridges stabilize proteins? A continuum electrostatic analysis. Protein Sci. 3, 211–226 (1994).

35. S. Kumar, R. Nussinov, Salt bridge stability in monomeric proteins. J. Mol. Biol. 293, 1241–55 (1999).

36. J. E. Donald, D. W. Kulp, W. F. DeGrado, Salt bridges: Geometrically specific, designable interactions. Proteins Struct. Funct. Bioinforma. 79, 898–915 (2011).

37. R. Pal et al., Syn vs Anti Carboxylic Acids in Hybrid Peptides: Experimental and Theoretical Charge Density and Chemical Bonding Analysis. J. Phys. Chem. A. 122, 3665–3679 (2018).

38. I. Kursula, S. Partanen, A. M. Lambeir, R. K. Wierenga, The importance of the conserved Arg191-Asp227 salt bridge of triosephosphate isomerase for folding, stability, and catalysis. FEBS Lett. 518, 39–42 (2002).

39. D. E. Anderson, W. J. Becktel, F. W. Dahlquist, pH-Induced Denaturation of Proteins: A Single Salt Bridge Contributes 3-5 kcal/mol to the Free Energy of Folding of T4 Lysozyme. Biochemistry. 29, 2403–2408 (1990).

40. D. Sali, M. Bycroft, A. R. Fersht, Surface electrostatic interactions contribute little to stability of barnase. J. Mol. Biol. 220, 779–788 (1991).

41. Z. Hong, Z. Ahmed, S. A. Asher, Circular dichroism and ultraviolet resonance raman indicate little Arg-Glu side chain α-helix peptide stabilization. J. Phys. Chem. B. 115, 4234–4243 (2011).

42. D. W. Carey, Joel F Schildbach, Robert T. Sauer, Are buried salt bridges important for protein stability and conformational specificity? Nat. Struct. Biol. 2, 122–128 (1995).

43. H. S. Andersson et al., The α-defensin salt-bridge induces backbone stability to facilitate folding and confer proteolytic resistance. Amino Acids. 43, 1471–1483 (2012).

44. A. D. Gruia, S. Fischer, J. C. Smith, Molecular dynamics simulation reveals a surface salt bridge forming a kinetic trap in unfolding of truncated Staphylococcal nuclease. Proteins Struct. Funct. Genet. 50, 507–515 (2003).

45. H. Meuzelaar et al., Solvent-exposed salt bridges influence the kinetics of α-helix folding and unfolding. J. Phys. Chem. Lett. 5, 900–904 (2014).

46. H. Meuzelaar, J. Vreede, S. Woutersen, Influence of Glu/Arg, Asp/Arg, and Glu/Lys Salt Bridges on α-Helical Stability and Folding Kinetics. Biophys. J. 110, 2328–2341 (2016).

47. a V Buevich, Q. H. Dai, X. Liu, B. Brodsky, J. Baum, Site-specific NMR monitoring of cistrans isomerization in the folding of the proline-rich collagen triple helix. Biochemistry. 39, 4299–308 (2000).

48. K. Mizuno et al., Kinetic hysteresis in collagen folding. Biophys. J. 98, 3004–3014 (2010).

49. J. M. Sanchez-Ruiz, Protein kinetic stability. Biophys. Chem. 148, 1–15 (2010).

50. S. Romero-Romero, M. Costas, A. Rodríguez-Romero, D. Alejandro Fernández-Velasco, Reversibility and two state behaviour in the thermal unfolding of oligomeric TIM barrel proteins. Phys. Chem. Chem. Phys. 17, 20699–20714 (2015).

51. K. Okuyama, K. Miyama, K. Mizuno, H. P. Bächinger, Crystal structure of (Gly-Pro-Hyp)9: Implications for the collagen molecular model. Biopolymers. 97, 607–616 (2012).

52. P. Kröger, S. Shanmugaratnam, N. Ferruz, K. Schweimer, B. Höcker, A comprehensive binding study illustrates ligand recognition in the periplasmic binding protein PotF. Structure. 29, 433-443.e4 (2021).

53. J. D. Chodera, W. C. Swope, J. W. Pitera, K. A. Dill, Long-time protein folding dynamics from short-time molecular dynamics simulations. Multiscale Model. Simul. 5, 1214–1226 (2006).

54. N. Plattner, S. Doerr, G. De Fabritiis, F. Noé, Complete protein-protein association kinetics in atomic detail revealed by molecular dynamics simulations and Markov modelling. Nat. Chem. 9, 1005–1011 (2017).

55. J. Xiao, H. Cheng, T. Silva, J. Baum, B. Brodsky, Osteogenesis imperfecta missense mutations in collagen: Structural consequences of a glycine to alanine replacement at a highly charged site. Biochemistry. 50, 10771–10780 (2011).

56. X. Liu, S. Kim, Q. H. Dai, B. Brodsky, J. Baum, Nuclear magnetic resonance shows asymmetric loss of triple helix in peptides modeling a collagen mutation in brittle bone disease. Biochemistry. 37, 15528–33 (1998).

57. K. Xu, I. Nowak, M. Kirchner, Y. Xu, Recombinant collagen studies link the severe conformational changes induced by osteogenesis imperfecta mutations to the disruption of a set of interchain salt bridges. J. Biol. Chem. 283, 34337–44 (2008).

58. M. J. Landrum et al., ClinVar: improving access to variant interpretations and supporting evidence. Nucleic Acids Res. 46, D1062–D1067 (2018).

59. I. F. A. C. Fokkema et al., LOVD v.2.0: The next generation in gene variant databases. Hum. Mutat. 32, 557–563 (2011).

60. D. K. Crockett et al., The Alport syndrome COL4A5 variant database. Hum. Mutat. 31, E1652–7 (2010).

61. D. O. Sillence, A. Senn, D. M. Danks, Genetic heterogeneity in osteogenesis imperfecta. J. Med. Genet. 16, 101–116 (1979).

62. J. C. Marini et al., Consortium for osteogenesis imperfecta mutations in the helical domain of type I collagen: Regions rich in lethal mutations align with collagen binding sites for integrins and proteoglycans. Hum. Mutat. 28, 209–221 (2007).

63. K. Salacinska et al., Novel Mutations Within Collagen Alpha1(I) and Alpha2(I) Ligand-Binding Sites, Broadening the Spectrum of Osteogenesis Imperfecta – Current Insights Into Collagen Type I Lethal Regions. Front. Genet. 12 (2021), doi:10.3389/fgene.2021.692978.

64. J. Des Parkin et al., Mapping structural landmarks, ligand binding sites, and missense mutations to the collagen IV heterotrimers predicts major functional domains, novel interactions, and variation in phenotypes in inherited diseases affecting basement membranes. Hum. Mutat. 32, 127–143 (2011).

65. G. Bellamy, P. Bornstein, Evidence for procollagen, a biosynthetic precursors of collagen. Proc. Natl. Acad. Sci. U. S. A. 68, 1138–1142 (1971).

66. E. Leikina, M. V. Mertts, N. Kuznetsova, S. Leikin, Type I collagen is thermally unstable at body temperature. Proc. Natl. Acad. Sci. 99, 1314–1318 (2002).

67. E. Makareeva, S. Leikin, Procollagen triple helix assembly: An unconventional chaperone-assisted polding paradigm. PLoS One. 2 (2007), doi:10.1371/journal.pone.0001029.

68. T. Koide et al., Specific recognition of the collagen triple helix by chaperone HSP47: Minimal structural requirement and spatial molecular orientation. J. Biol. Chem. 281, 3432–3438 (2006).

69. J. R. Macdonald, H. P. Bächinger, HSP47 Binds Cooperatively to Triple Helical Type I Collagen but Has Little Effect on the Thermal Stability or Rate of Refolding. J. Biol. Chem. 276, 25399–25403 (2001).

70. E. T. Abraham et al., Collagen’s primary structure determines collagen:HSP47 complex stoichiometry. J. Biol. Chem. 297, 101169 (2021).

71. Y. Ishida et al., Type I collagen in Hsp47-null cells is aggregated in endoplasmic reticulum and deficient in N-propeptide processing and fibrillogenesis. Mol. Biol. Cell. 17, 2346–2355 (2006).

72. Y. Matsuoka et al., Insufficient folding of type IV collagen and formation of abnormal basement membrane-like structure in embryoid bodies derived from Hsp47-null embryonic stem cells. Mol. Biol. Cell. 15, 4467–4475 (2004).

73. A. Nakai, M. Satoh, K. Hirayoshi, K. Nagata, Involvement of the stress protein HSP47 in procollagen processing in the endoplasmic reticulum. J. Cell Biol. 117, 903–914 (1992).

74. M. Satoh, K. Hirayoshi, S. I. Yokota, N. Hosokawa, K. Nagata, Intracellular interaction of collagen-specific stress protein HSP47 with newly synthesized procollagen. J. Cell Biol. 133, 469–483 (1996).

75. Y. Ishikawa, S. Ito, K. Nagata, L. Y. Sakai, H. P. Bächinger, Intracellular mechanisms of molecular recognition and sorting for transport of large extracellular matrix molecules. Proc. Natl. Acad. Sci. U. S. A. 113, E6036–E6044 (2016).

76. B. Ibarra-Molero, J. A. Zitzewitz, C. R. Matthews, Salt-bridges can Stabilize but do not Accelerate the Folding of the Homodimeric Coiled-coil Peptide GCN4-p1. J. Mol. Biol. 336, 989–996 (2004).

77. A. D. Stoycheva, J. N. Onuchic, C. L. Brooks, Effect of gatekeepers on the early folding kinetics of a model β-barrel protein. J. Chem. Phys. 119, 5722–5729 (2003).

78. E. Makareeva et al., Structural heterogeneity of type I collagen triple helix and its role in osteogenesis imperfecta. J. Biol. Chem. 283, 4787–98 (2008).

79. W. Li, L. Jaroszewski, A. Godzik, Clustering of highly homologous sequences to reduce the size of large protein database. Bioinformatics. 17, 282–283 (2011).

80. R. C. Team, R: A Language and Environment for Statistical Computing (2023).

81. J. Bella, A first census of collagen interruptions: Collagen’s own stutters and stammers. J. Struct. Biol. 186, 438–450 (2014).

82. J. M. Sanchez-Ruiz, Theoretical analysis of Lumry-Eyring models in differential scanning calorimetry. Biophys. J. 61, 921–935 (1992).

83. S. Romero-Romero et al., The Stability Landscape of de novo TIM Barrels Explored by a Modular Design Approach. J. Mol. Biol. 433 (2021), doi:10.1016/j.jmb.2021.167153.

84. S. Doerr, M. J. Harvey, F. Noé, G. De Fabritiis, HTMD: High-Throughput Molecular Dynamics for Molecular Discovery. J. Chem. Theory Comput. 12, 1845–1852 (2016).

