## Supplementary Material for "Code for Collagen Folding Deciphered"

**Supplementary table S1.** Frequency of collagen and collagen-like protein domains and their interruptions in humans, the three domains of life and viruses.

| | human | archaea | bacteria | eukarya | virus |
|---|---|---|---|---|---|
| Proteins from UniProtKb | 44 | 524 | 17728 | 39511 | 1963 |
| Proteins after clustering with CD-HIT* | - | 276 | 8314 | 11163 | 898 |
| Total collagen domains | 468 | - | - | - | - |
| Total collagen-like protein domains | - | 490 | 15302 | 39538 | 2672 |
| Total triplets | 10456 | 12581 | 446143 | 675106 | 60156 |
| Total interruptions | 394 | 165 | 5517 | 26220 | 1435 |

**human collagen alpha chains were not clustered*

### fibrillar collagens

#### collagen type I – $(\alpha1)_3$

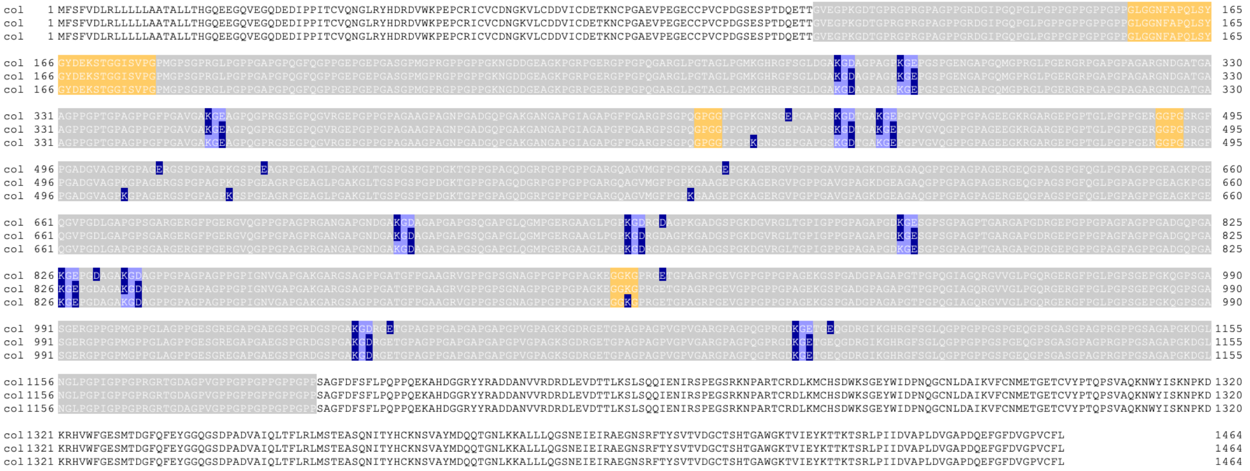

#### collagen type I – $(\alpha1)_2\alpha2$

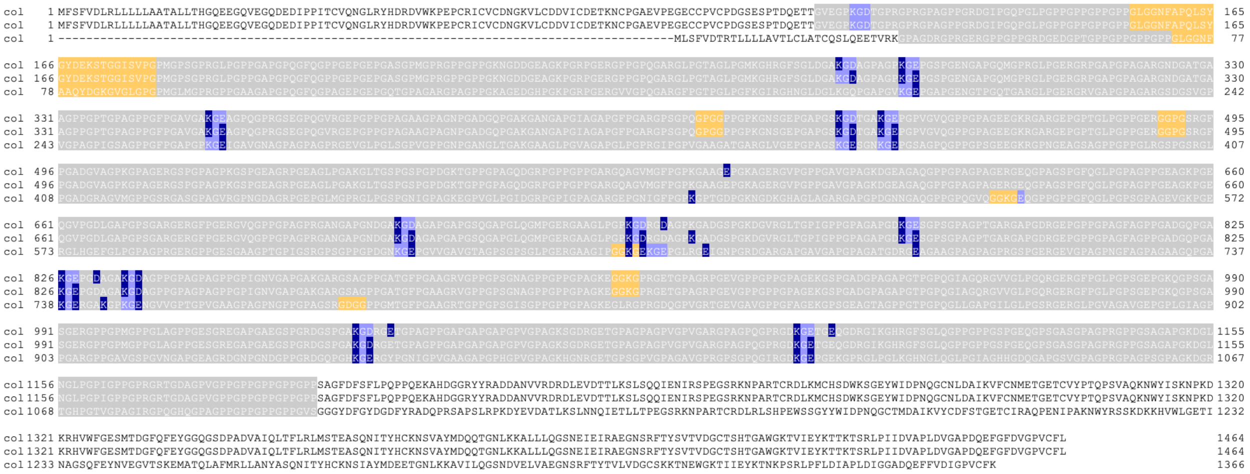

#### collagen type II – $(\alpha1)_3$

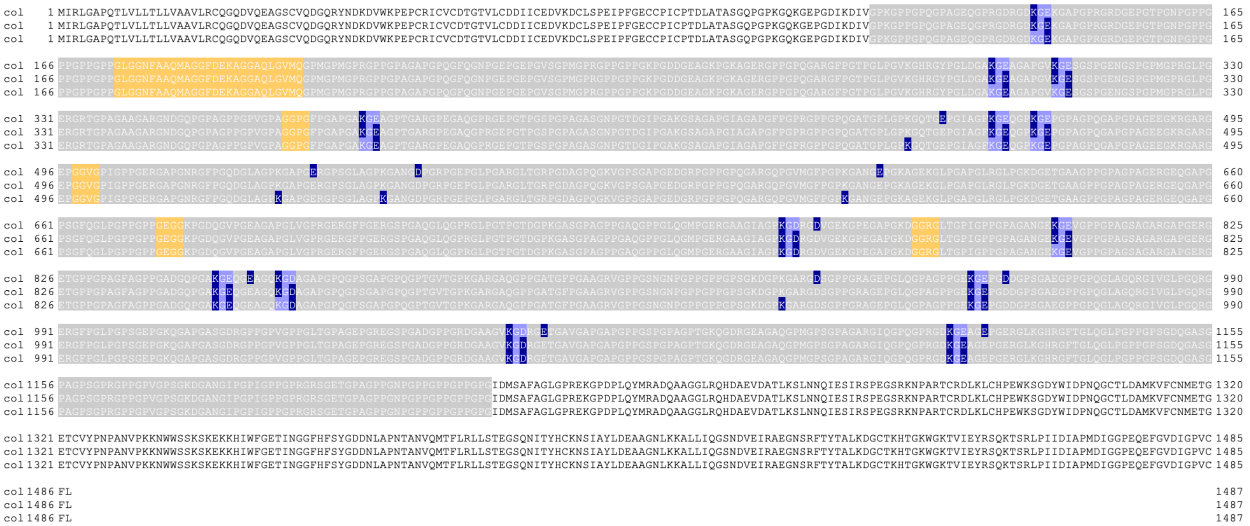

collagen type III – $(\alpha 1)_3$

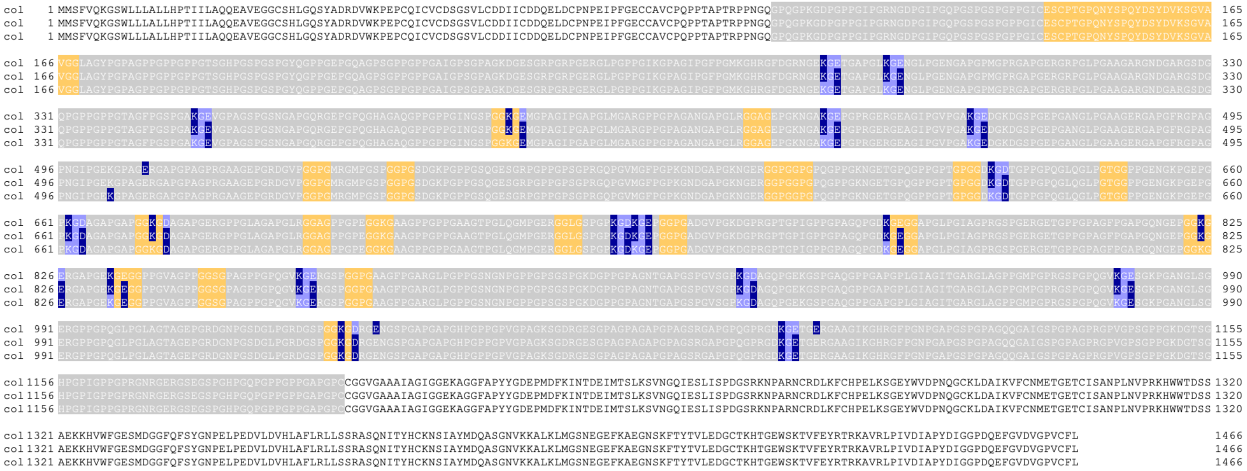

#### collagen type V – $(\alpha1)_3$

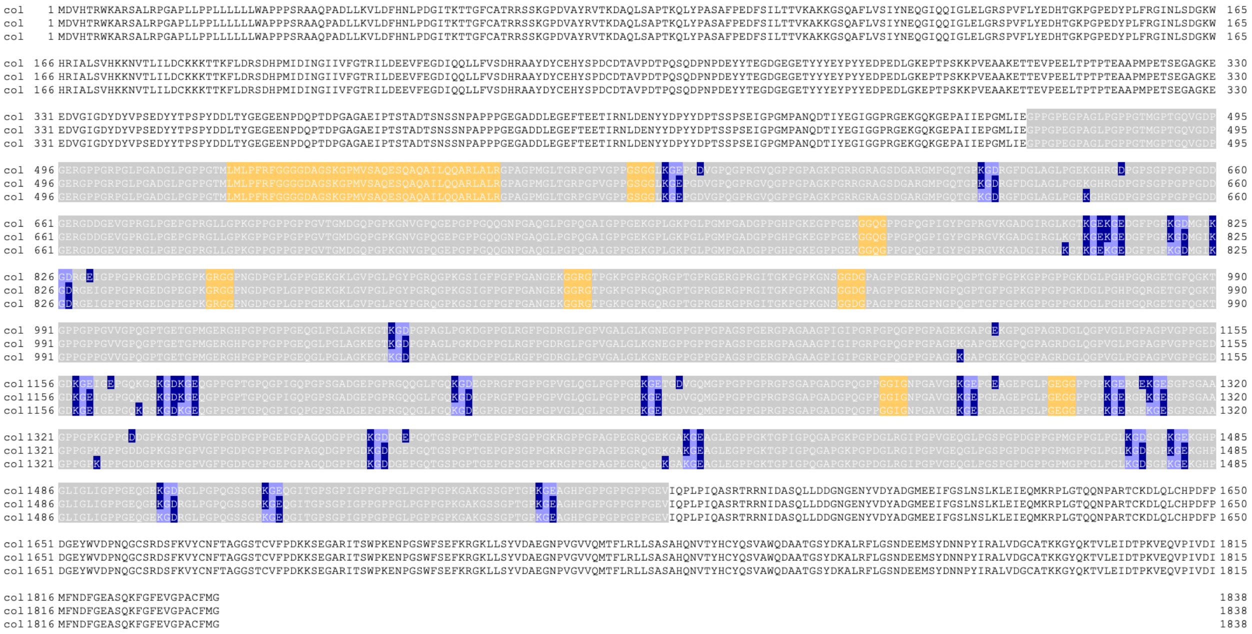

#### collagen type V – $(\alpha1)_2\alpha2$

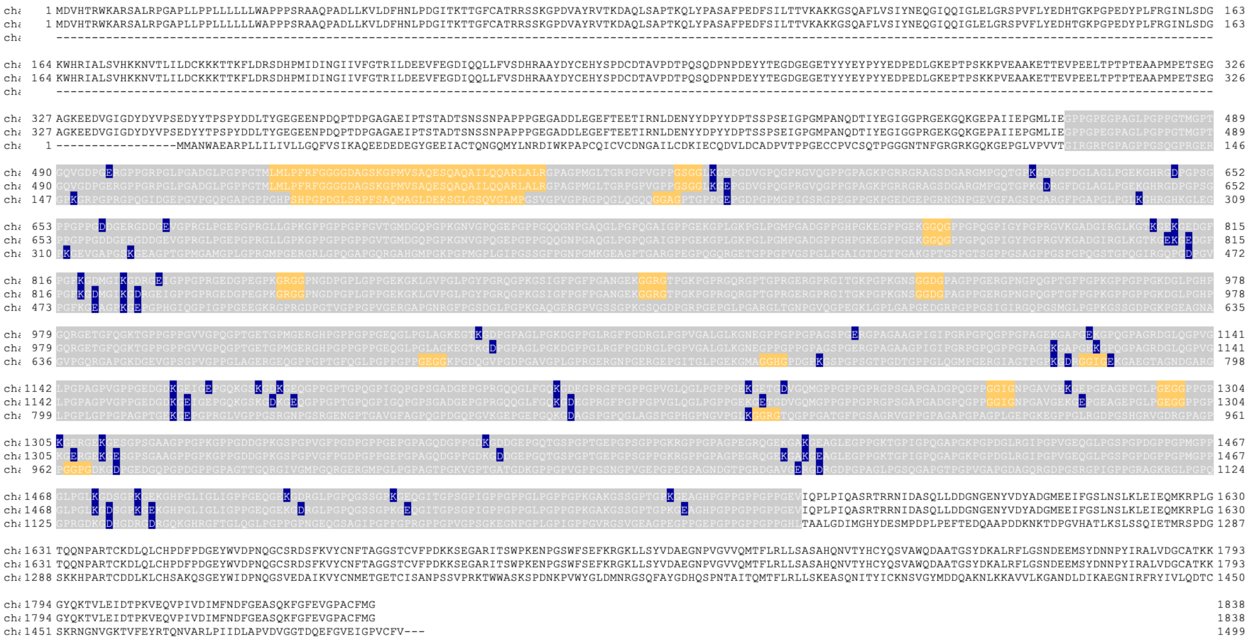

#### collagen type XXIV – (α1)$_3$

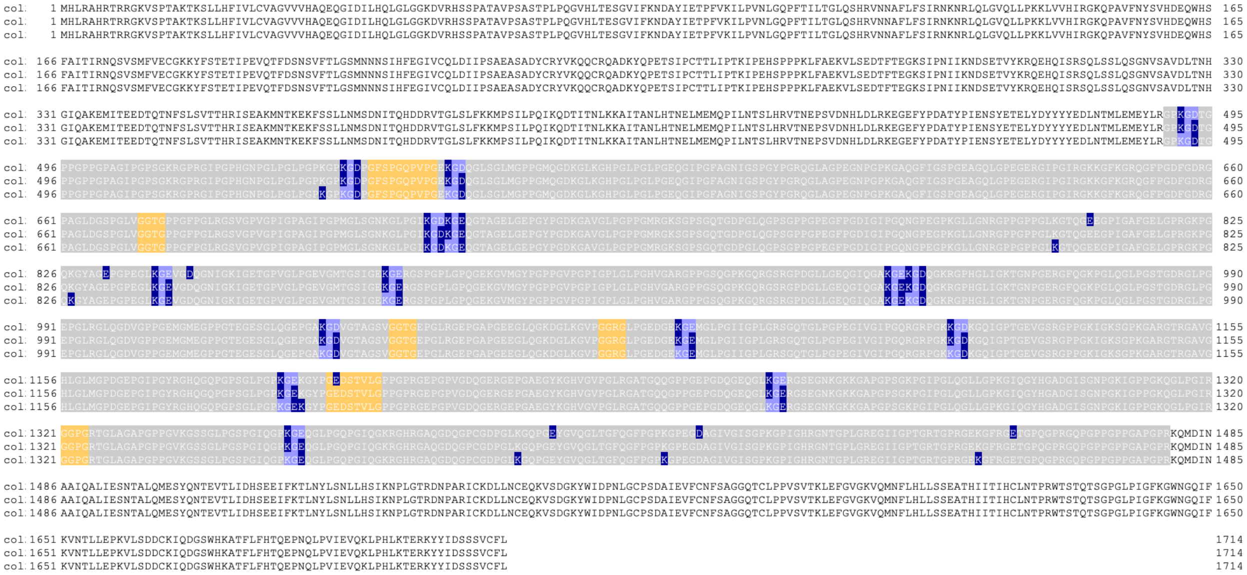

#### collagen type XXVII – (α1)$_3$

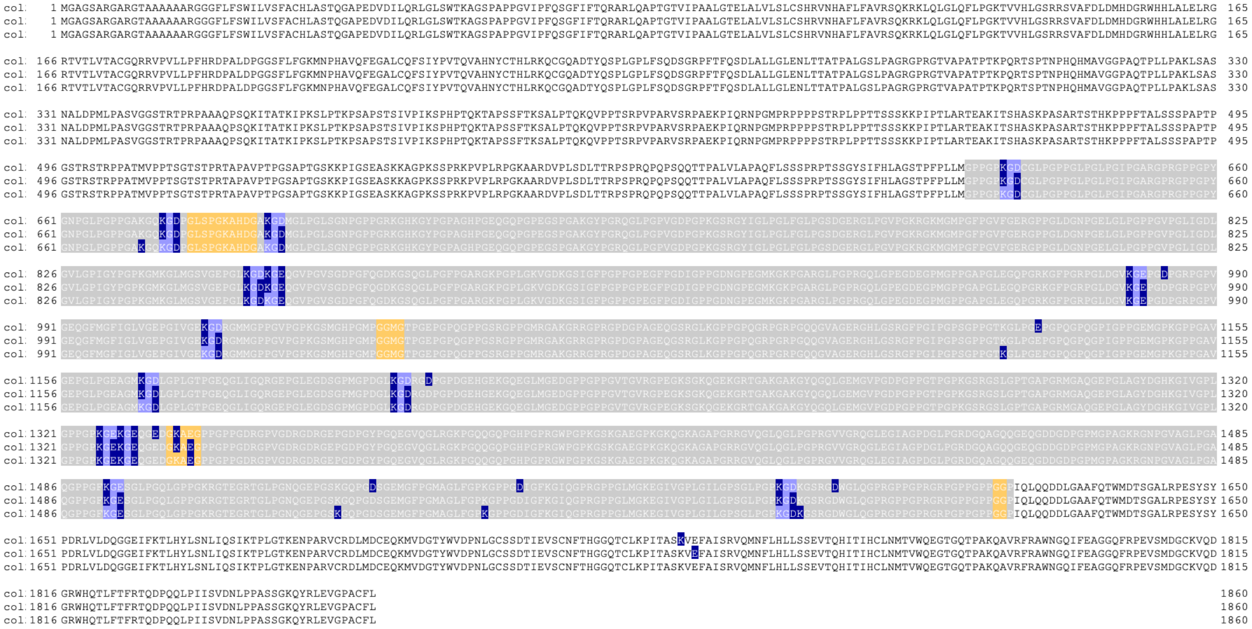

<u>network-forming collagens</u>

collagen type IV – (α1)₂α2

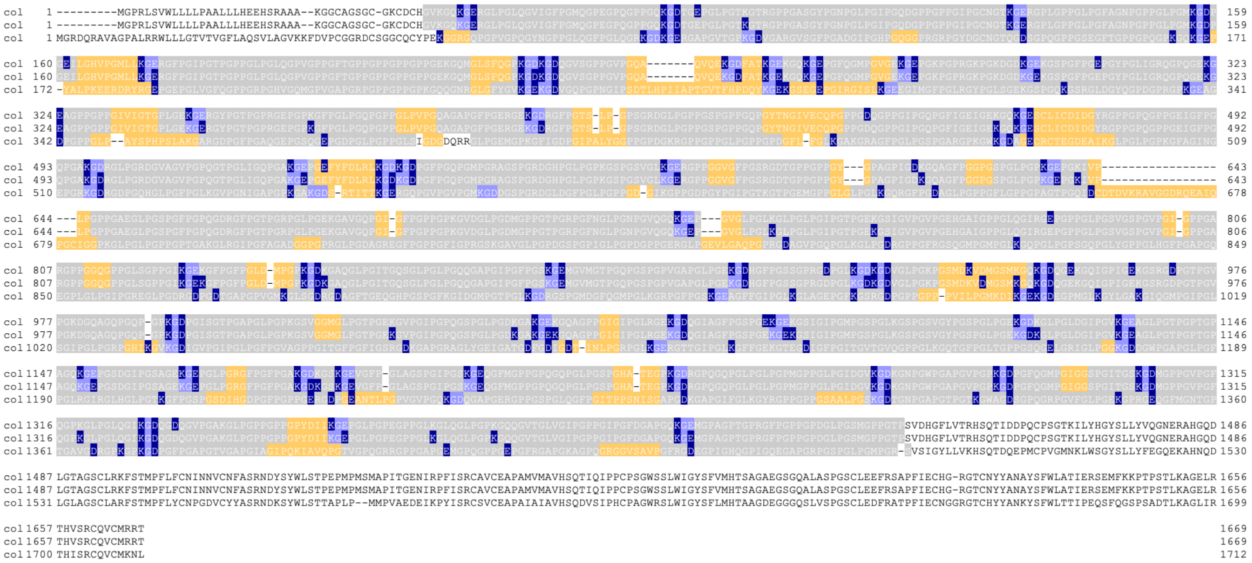

collagen type IV – (α5)₂α6

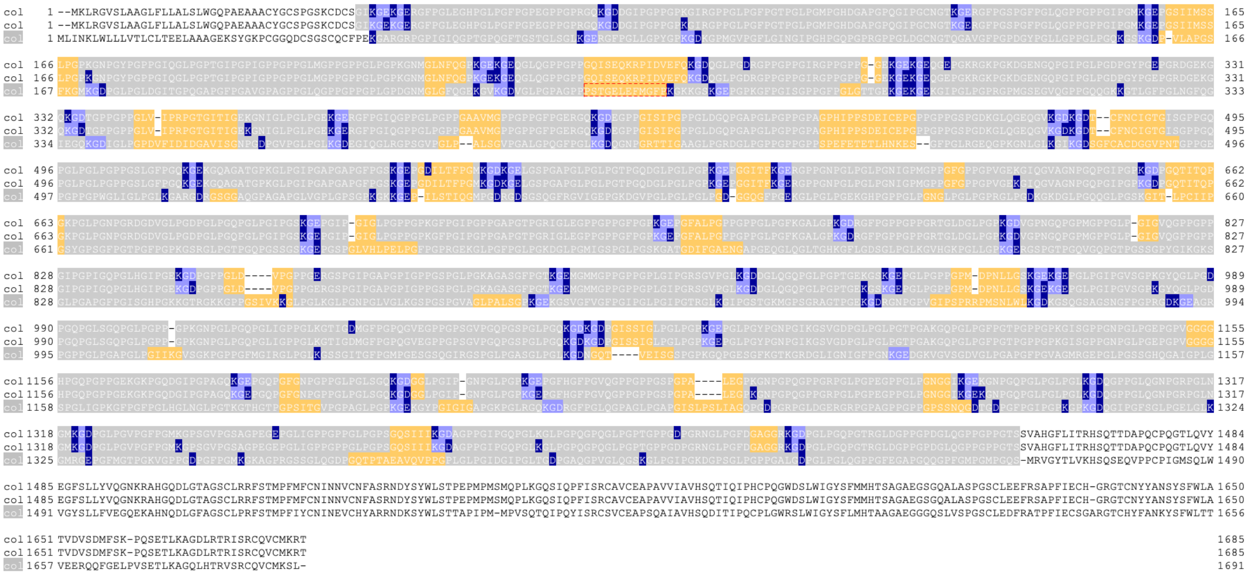

#### collagen type IV – α3α4α5

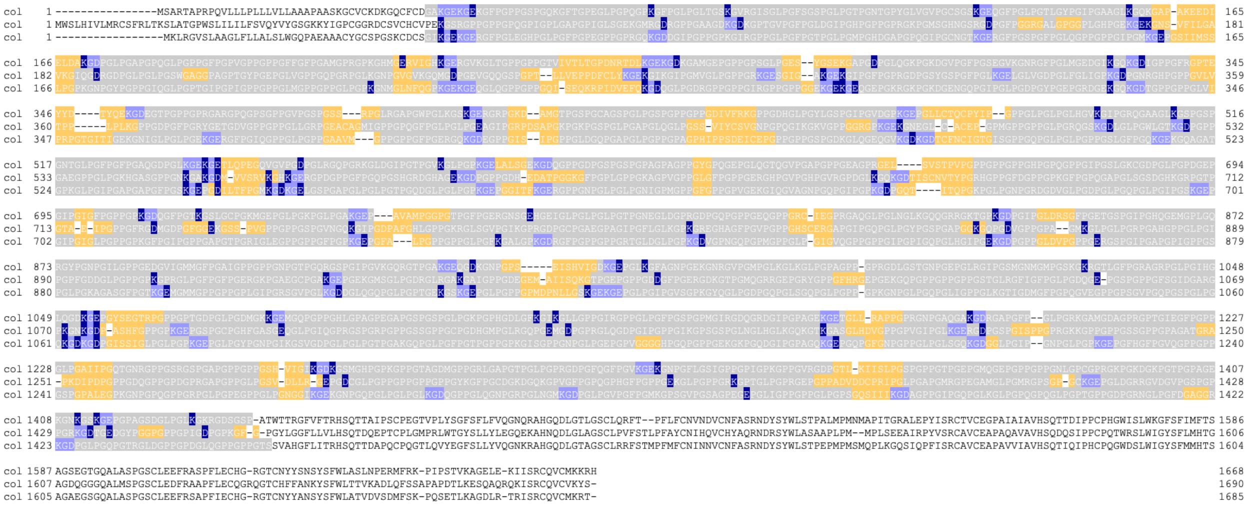

#### collagen type VIII – $(\alpha1)_3$

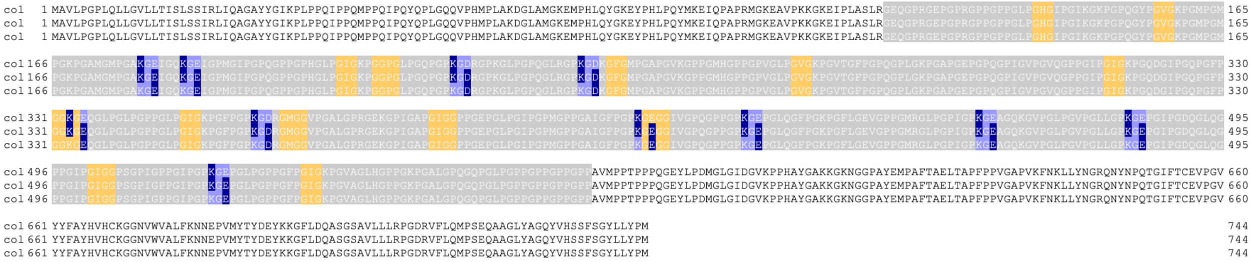

#### collagen type VIII – $(\alpha2)_3$

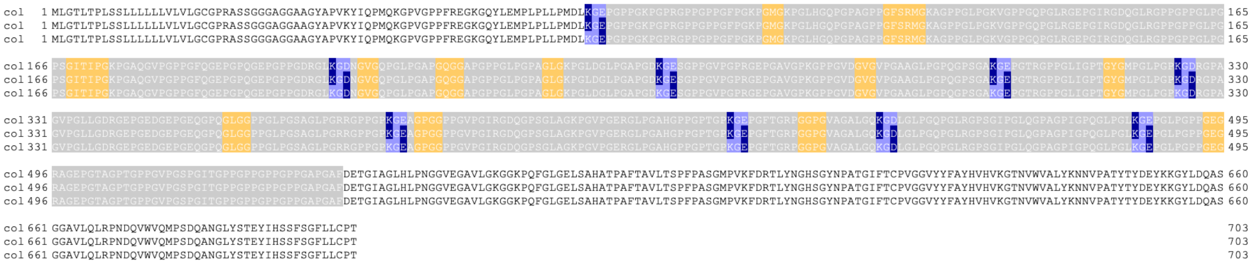

#### collagen type X – $(\alpha1)_3$

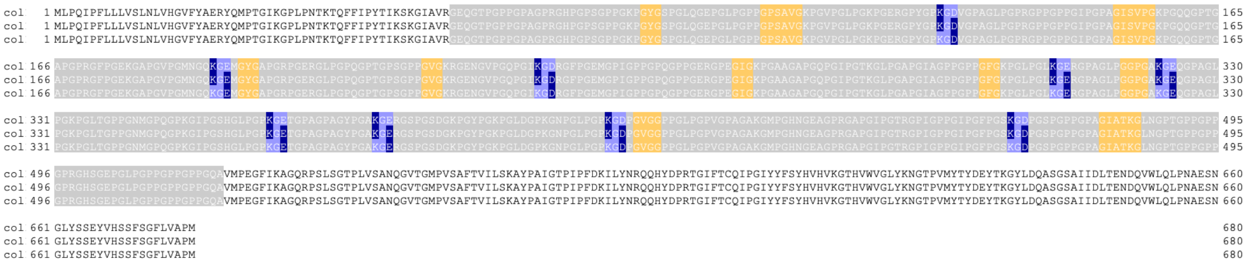

#### fibril-associated collagen with interrupted triple helices

##### collagen type IX – α1α2α3

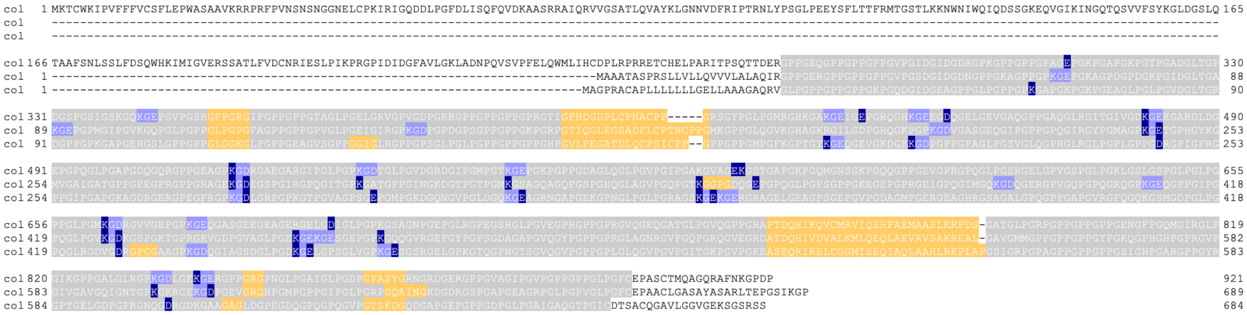

##### collagen type XII – (α1)₃

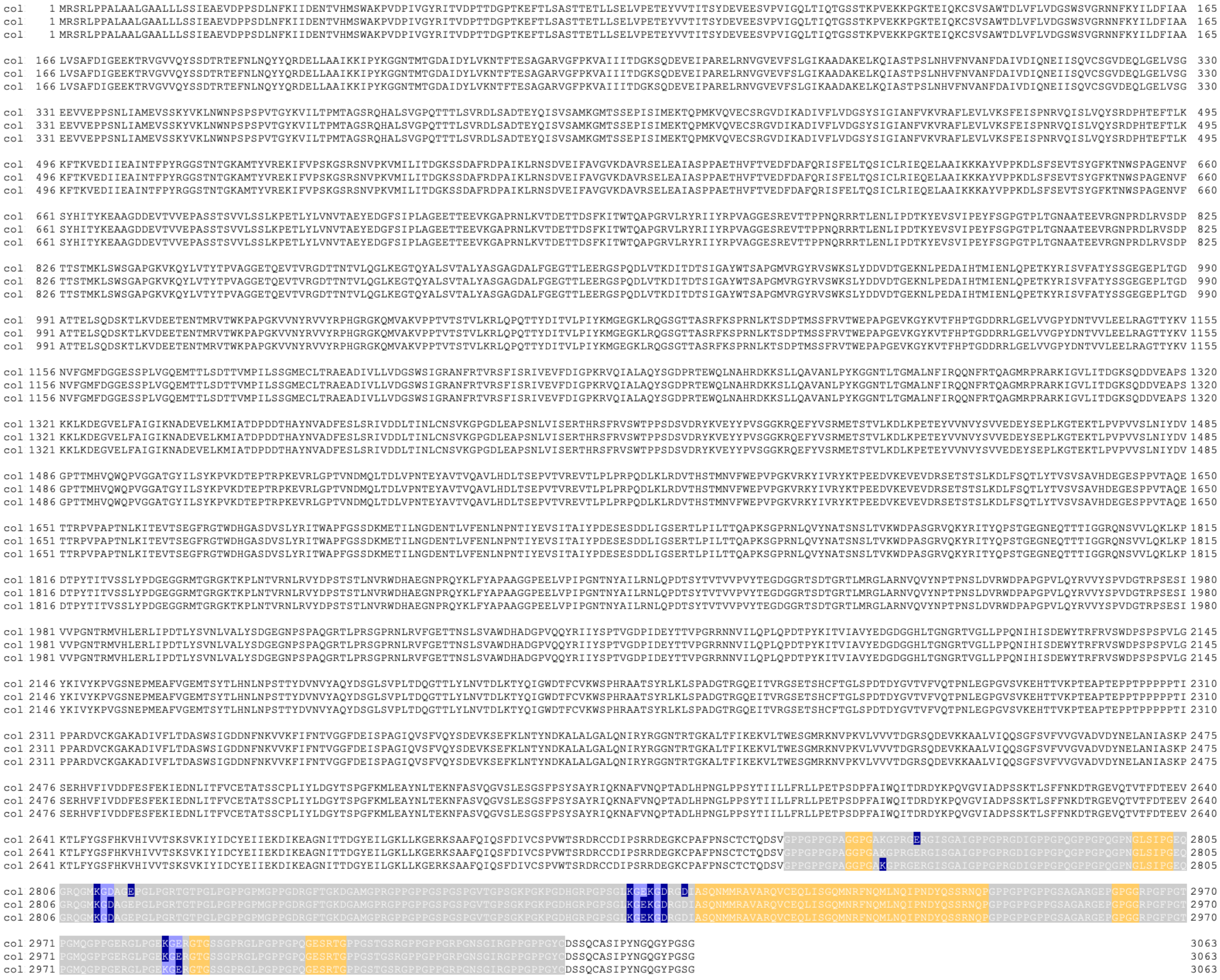

#### collagen type XIV – (α1)$_3$

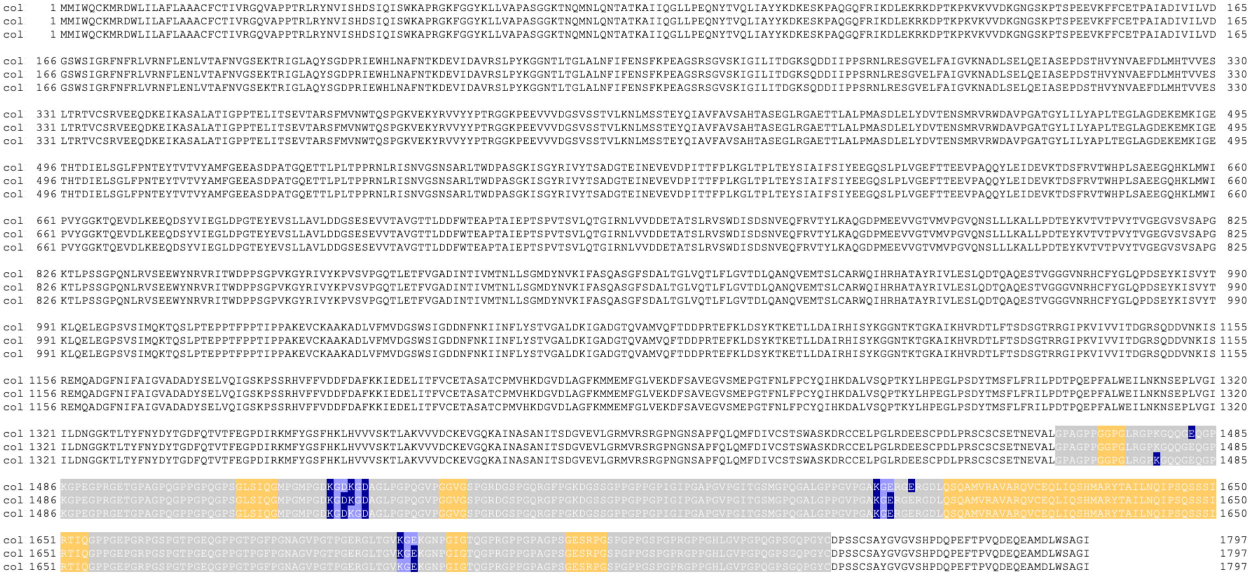

#### collagen type XVI – (α1)$_3$

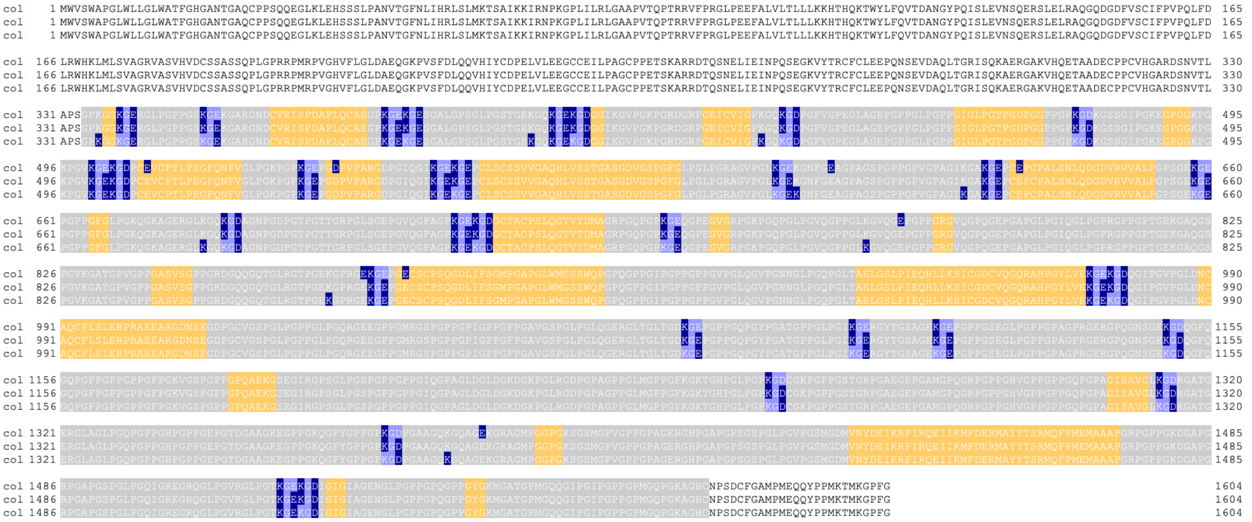

#### collagen type XIX – (α1)$_3$

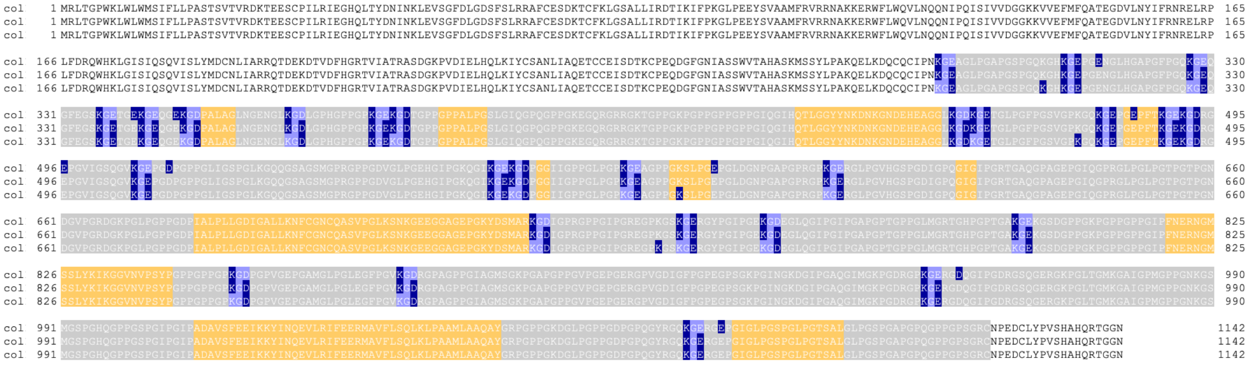

#### collagen type XX – (α1)$_3$

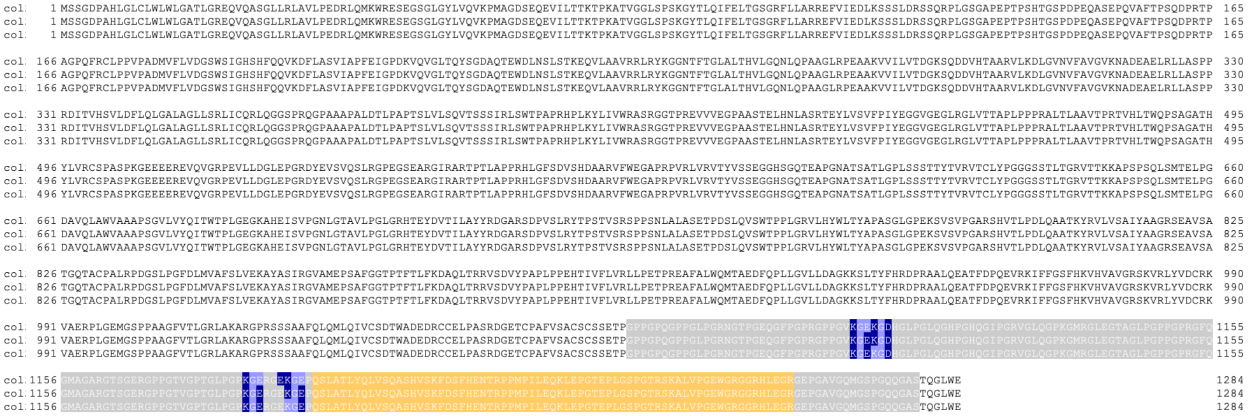

#### collagen type XXI – (α1)$_3$

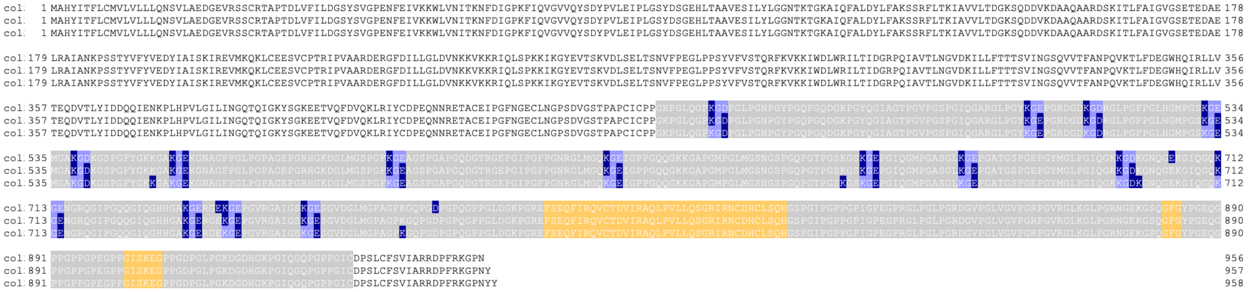

#### collagen type XXII – (α1)$_3$

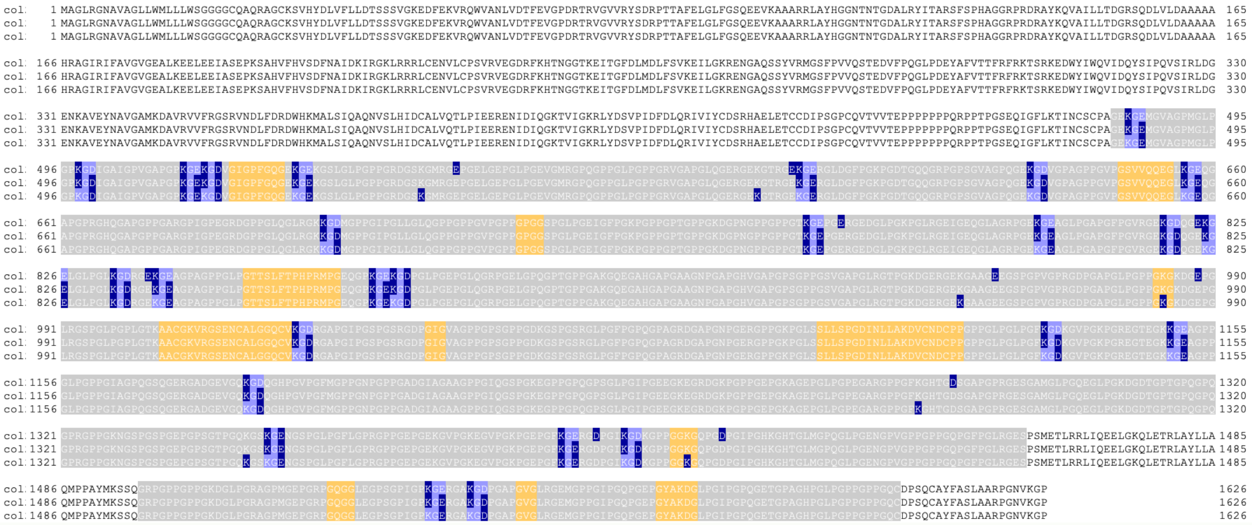

#### microfibrillar collagens

#### collagen type VI – α3α2α1

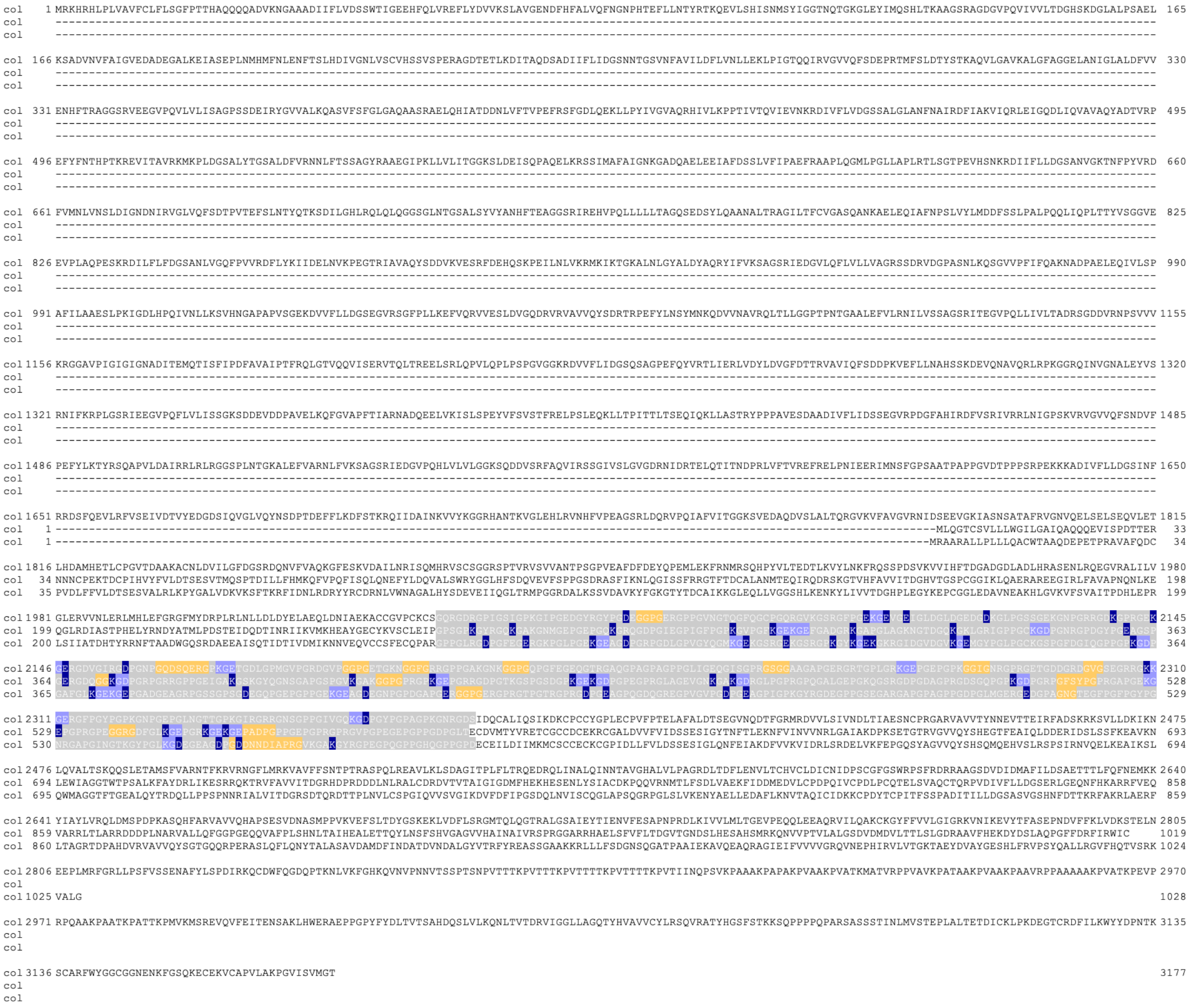

#### collagen type VI – α5α2α1

```
col    1 MKILLIIFVLIIWTETLADQSPGPGPVYADVVFLVDSSDHLGPKSFPFVKTFINKMINSLPIEANKYRVALAQYSDEFHSEFHLSTFKGRSPMLNHLKKNFQFIGGSLQIGKALQEAHRTYFSAPINGRDRKQFPPILVVLASAESEDEVEEASKALQKDGVKII  165
col      ---------------------------------------------------------------------------------------------------------------------------------------------------------------------
col      ---------------------------------------------------------------------------------------------------------------------------------------------------------------------

col  166 SVGVQKASEENLKAMATSHFHFNLRTIRDLSTFSQNMTQIIKDVTKYKEGAVDADMQVHFPISCQKDSLADLVFLVDESLGTGGNLRHLQTFLENITSSMDVKENCMRLGLMSYSNSAKTISFLKSSTTQSEFQQQIKNLSIQVGKSNTGAAIDQMRRDGFSESY  330
col      ---------------------------------------------------------------------------------------------------------------------------------------------------------------------
col      ---------------------------------------------------------------------------------------------------------------------------------------------------------------------

col  331 GSRRAQGVPQIAVLVTHRPSDDEVHDAALNLRLEDVNVFALSIQGANNTQLEEIVSYPPEQTISTLKSYADLETYSTKFLKKLQNEIWSQISTYAEQRNLDKTGCVDTKEADIHFLIDGSSSIQEKQFEQIKRFMLEVTEMFSIGPDKVRVGVVQYSDDTEVEFY  495
col      ---------------------------------------------------------------------------------------------------------------------------------------------------------------------
col      ---------------------------------------------------------------------------------------------------------------------------------------------------------------------

col  496 ITDYSNDIDLRKAIFNIKQLTGGTYTGKALDYILQIIKNGMKDRMSKVPCYLIVLTDGMSTDRVVEPAKRLRAEQITVHAVGIGAANKIELQEIAGKEERVSFGQNFDALKSIKNEVVREICAEKGCEDMKADIMFLVDSSWSIGNENFRKMKIFMKNLLTKIQI  660
col      ---------------------------------------------------------------------------------------------------------------------------------------------------------------------
col      ---------------------------------------------------------------------------------------------------------------------------------------------------------------------

col  661 GADKTQIGVVQFSDKTKEEFQLNRYFTQQEISDAIDRMSLINEGTLTGKALNFVGQYFTHSKGARLGAKKFLILITDGVAQDDVRDPARILRGKDVTIFSVGVYNANRSQLEEISGDSSLVFHVENFDHLKALERKLIFRVCALHDCKRITLLDVVFVLDHSGSI  825
col      ---------------------------------------------------------------------------------------------------------------------------------------------------------------------
col      ---------------------------------------------------------------------------------------------------------------------------------------------------------------------

col  826 KKQYQDHMINLTIHLVKKADVGRDRVQFGALKYSDQPNILFYLNTYSNRSAIIENLRKRRDTGGNTYTAKALKHANALFTEEHGSRIKQNVKQMLIVITDGESHDHDQLNDTALELRNKGITIFAVGVGKANQKELEGMAGNKNNTIYVDNFDKLKDVFTLVQER  990
col      ---------------------------------------------------------------------------------------------------------------------------------------------------------------------
col      ---------------------------------------------------------------------------------------------------------------------------------------------------------------------

col  991 MCTEAPEVCHLQEADVIFLCDGSDRVSNSDFVTMTTFLSDLIDNFDIQSQRMKIGMAQFGSNYQSIIELKNSLTKTQWKTQIQNVSKSGGFPRIDFALKKVSNMFNLHAGGRRNAGVPQTLVVITSGDPRYDVADAVKTLKDLGICVLVLGIGDVYKEHLLPITG 1155
col      ---------------------------------------------------------------------------------------------------------------------------------------------------------------------
col      ---------------------------------------------------------------------------------------------------------------------------------------------------------------------

col 1156 NSEKIITFQDFDKLKNVDVKKRIIREICQSCGKTNCFMDIVVGFDISTHVQGQPLFQGHPQLESYLPGILEDISSIKGVSCGAGTEAQVSLAFKVNSDQGFPAKFQIYQKAVFDSLLQVNVSGPTHLNAQFLRSLWDTFKDKSASRGQVLLIFSDGLQSESNIML 1320
col      ---------------------------------------------------------------------------------------------------------------------------------------------------------------------
col      ---------------------------------------------------------------------------------------------------------------------------------------------------------------------

col 1321 ENQSDRLREAGLDALLVVSLNTTAHHEFSSFEFGKRFDYRTHLTIGMRELGKKLSQYLGNIAERTCCCTFCKCPGIPGPHGTRGLQAMKGSQGLKGSRGHRGEDGNPGVRDTGPQGDKGIAGCPGAWGQKGLKGFSGPKGGHGDDGIDGLDGEEGCHGFPGIKG 1485
col    1 ----------------------------------------------------------------CYKVSCLEIPGPSGPKGYRGQKGAKGNMGEPGEPGQKGRQGDPGIEGPIGFPGPKGVPGPKGEKGEFGADGRKGAPGLAGKNGTDGQKGKLGRIGPPGCKG  101
col    1 ----------------------------------------------------------------CCSFECQPARGPPGLRGDPGFEGERGKPGLPGEKGEAGDPGRPGDLPGVGYQGMKGEKGSRGEKGSRGPKGYKGEKGKRGIDGVDGVKGEMGYPGLPGCKG  101

col 1486 EKGDPGSQGSPGSRGAPGQYGEKGFPGDPGNPGQNNNIKGQKGSKGEQGRQGRSGQKGVQGSPSSRGSRGREGQRGLRGVSGEPGNPGPTGTLGAEGLQGPQGSQGNPGRKGEKGSQGQKGPQGSPGLMGAKGSTGRPGLLGKKGEPGLPGDLGPVGQTGQRGRQ 1650
col  102 DPGNRGPDGYPGEAGSPGERGDQGGKGDPGRPGR----RGPPGEIGAKGSKGYQGNSGAPGSPGVKGAKGGPGPRGPKGEPGRRGDPGTKGSPGSDGPKGEKGDPGPEGPRGLAGEVGNKGAKGDRGLPGPRGPQGALGEPGKQGSRGDPGDAGPRGDSGQPGPK  262
col  102 SPGFDGIQGPPGPKGDPGAFGLKGEKGEPGADGE----AGRPGSSGPSGDEGQPGEPGPPGEKGEAGDEGNPGPDGAPGERGGPGERGPRGTPGTRGPRGDPGEAGPQGDQGREGPVGVPGDPGEAGPIGPKGYRGDEGPPGSEGARGAPGPAGPPGDPGLMGER  262

col 1651 GDSGIPGYGQMGRKGVKGPRGFPGDAGQKGDIGNPGIPGGPGPKGFRGLALTVGLK-GEEGSRGLPGPPGQRGIKGMAGQPVYSQCDLIRFLREHSPCWKEKCPAYPTELVFALDNSYDVTEESFNKTRDIITSIVNDLNIRENNCPVGARVAMVSYNSGTSYLI 1814
col  263 GDPGRPGFSYPGPRGAPGEKGEPGPRGPEGGRGDFGLKGEPGRKGEKGEPADPGPP-GEPGPRGPRGVPGPEGEPGPPGDP                                                                                     342
col  263 GEDGPAGNGTEGFPGFPGYPGNRGAPGINGTKGYPGLKGDEGEAGDPGDDNNDIAPRGVKGAKGYRGPEGPQGPPGHQGPP                                                                                     343

col 1815 RWSDYNRKKQLLQQLSQIKYQDTTEPRDVGNAMRFVTRNVFKRTYAGANVRRVAVFFSNGQTASRSSIITATMEFSALDISPTVFAFDERVFLEAFGFDNTGTFQVIPVPPNGENQTLERLRRCALCYDKCFPNACIREAFLPEDSYMDVVFLIDNSRNIAKDEF 1979
col
col

col 1980 KAVKALVSSVIDNFNIASDPLISDSGDRIALLSYSPWESSRRKMGTVKTEFDFITYDNQLLMKNHIQTSFQQLNGEATIGRALLWTTENLFPETPYLRKHKVIFVVSAGENYERKEFVKMMALRAKCQGYVIFVISLGSTRKDDMEELASYPLDQHLIQLGRIHK 2144
col
col

col 2145 PDLNYIAKFLKPFLYSVRRGFNQYPPPMLEDACRLINLGGENIQNDGFQFVTELQEDFLGGNGFIGQELNSGRESPFVKTEDNGSDYLVYLPSQMFEPQKLMINYEKDQKSAEIASLTSGHENYGRKEEPDHTYEPGDVSLQEYYMDVAFLIDASQRVGSDEFKE 2309
col
col

col 2310 VKAFITSVLDYFHIAPTPLTSTLGDRVAVLSYSPPGYMPNTEECPVYLEFDLVTYNSIHQMKHHLQDSQQLNGDVFIGHALQWTIDNVFVGTPNLRKNKVIFVISAGETNSLDKDVLRNVSLRAKCQGYSIFVFSFGPKHNDKELEELASHPLDHHLVQLGRTHK 2474
col
col

col 2475 PDWNYIIKFVKPFVHLIRRAINKYPTEDMKATCVNMTSPNPENGGTENTVLLLPGIYEIKTENGDLFDEFDSQAQHLLVLGNNHSSGSETATDLMQKLYLLFSTEKLAMKDKEKAHLEEISALVVDKQQEKEDKEMEATDI                        2615
col
col
```

#### collagen type VI – α6α2α1

```
col    1 MMLLILFLVIICSHISVNQDSGPEYADVVFLVDSSDRLGSKSFPFVKMFITKMISSLPIEADKYRVALAQYSDKLHSEFHLSTFKGRSPMLNHLRKNFGFIGGSLQIGKALQEAHRTYFSAPANGRDKKQFPPILVVLASSESEDNVEEASKALRKDGVKIISVG  165
col      ---------------------------------------------------------------------------------------------------------------------------------------------------------------------
col      ---------------------------------------------------------------------------------------------------------------------------------------------------------------------

col  166 VQKASEENLKAMATSQFHFNLRTVRDLSMFSQNMTHIIKDVIKYKEGAVDDIFVEACQGPSMADVVFLLDMSINGSEENFDYLKGFLEESVSALDIKENCMRVGLVAYSNETKVINSLSMGINKSEVLQHIQNLSPRTGKAYTGAAIKKLRKEVFSARNGSRKNQ  330
col      ---------------------------------------------------------------------------------------------------------------------------------------------------------------------
col      ---------------------------------------------------------------------------------------------------------------------------------------------------------------------

col  331 GVPQIAVLVTHRDSEDNVTKAAVNLRREGVTIFTLGIEGASDTQLEKIASHPAEQYVSKLKTFADLAAHNQTFLKKLRNQITHTVSVFSERTETLKSGCVDTEEADIYLLIDGSGSTQATDFHEMKTFLSEVVGMFNIAPHKVRVGAVQYADSWDLEFEINKYSN  495
col      ---------------------------------------------------------------------------------------------------------------------------------------------------------------------
col      ---------------------------------------------------------------------------------------------------------------------------------------------------------------------

col  496 KQDLGKAIENIRQMGGNTNTGAALNFTLSLLQKAKKQRGNKVPCHLVVLTNGMSKDSILEPANRLREEHIRVYAIGIKEANQTQLREIAGEEKRVYYVHDFDALKDIRNQVVQEICTEEACKEMKADIMFLVDSSGSIGPENFSKMKTFMKNLVSKSQIGPDRVQ  660
col      ---------------------------------------------------------------------------------------------------------------------------------------------------------------------
col      ---------------------------------------------------------------------------------------------------------------------------------------------------------------------

col  661 IGVVQFSDINKEEFQLNRFMSQSDISNAIDQMAHIGQTTLTGSALSFVSQYFSPTKGARPNIRKFLILITDGEAQDIVKEPAVVLRQEGVIIYSVGVFGSNVTQLEEISGRPEMVFYVENFDILQRIEDDLVFGICSPREECKRIEVLDVVFVIDSSGSIDYDEY  825
col      ---------------------------------------------------------------------------------------------------------------------------------------------------------------------
col      ---------------------------------------------------------------------------------------------------------------------------------------------------------------------

col  826 NIMKDFMIGLVKKADVGKNQVRFGALKYADDPEVLFYLDDFGTKLEVISVLQNDQAMGGSTYTAEALGFSDHMFTEARGSRLNKGVPQVLIVITDGESHDADKLNATAKALRDKGILVLAVGIDGANPVELLAMAGSSDKYFFVETFGGLKGIFSDVTASVCNSS  990
col      ---------------------------------------------------------------------------------------------------------------------------------------------------------------------
col      ---------------------------------------------------------------------------------------------------------------------------------------------------------------------

col  991 KVDCEIDKVDLVFLMDGSTSIQPNDFKKMKEFLASVVQDFDVSLNRVRIGAAQFSDTYHPEFPLGTFIGEKEISFQIENIKQIFGNTHIGAALREVEHYFRPDMGSRINTGTPQVLLVLTDGQSQDEVAQAAEALRHRGIDIYSVGIGDVDDQQLIQITGTAEKK 1155
col      ---------------------------------------------------------------------------------------------------------------------------------------------------------------------
col      ---------------------------------------------------------------------------------------------------------------------------------------------------------------------

col 1156 LTVHNFDELKKVNKRIVRNICTTAGESNCFVDVVVGFDVSTQEKGQTLLEGQPWMETYLQDILRAISSLNGVSCEVGTETQVSVAFQVTNAMEKYSPKFEIYSENILNSLKDITVKGPSLLNANLLDSLWDTFQNKSAARGKVVLLFSDGLDDDVEKLEQKSDEL 1320
col      ---------------------------------------------------------------------------------------------------------------------------------------------------------------------
col      ---------------------------------------------------------------------------------------------------------------------------------------------------------------------

col 1321 RKEGLNALITVALDGPADSSDLADLPYIEFGKGFEYRTQLSIGMRELGSRLSKQLVNVAERTCCCLFCKCIGGDGTMGDPGPPGKRGPPGPKGSEGYLGEEGIAGERGAPGPVGEQGTKGCYGTKGPKGNRGLNGQEGEVGENGIDGLNGEQGDNGLPGRKGEKG 1485
col    1 -------------------------------------------------------------CYKVSCLEIPGPSGPKGYRGQKGAKGNMGEPGEPGQKGRQGDPGIEGPIGFPGPKGVPGPKGEKGEFGADGRKGAPGLAGKNGTDGQKGKLGRIGPPGCKGDPG  104
col    1 -------------------------------------------------------------CCSFECQPARGPPGLRGDPGFEGERGKPGLPGEKGEAGDPGRPGDLPGVGYQGMKGEKGSRGEKGSRGPKGYKGEKGKRGIDGVDGVKGEMGYPGLPGCKGSPG  104

col 1486 DEGSQGSPGKRGTPGDRGAKGLRGDPGAPGVDSSIEGPTGLKGERGRQGRRGWPGPPGTPGSRRKTAAHGRRGHTGPQGTAGIPGPDGLEGSLGLKGPQGPRGEAGVKGEKGGVGSKGPQGPPGPGGEAGNQGRLGSQGNKGEPGDLGEKGAVGFPGPRGLQGND 1650
col  105 NRGPDGYPGEAGSPGERGDQGGKGDPGRPGR----RGPPGEIGAKGSKGYQGNSGAPGSPGVKGAKGGPGPRGPKGEPGRRGDPGTKGSPGSDGPKGEKGDPGPEGPRGLAGEVGNKGAKGDRGLPGPRGPQGALGEPGKQGSRGDPGDAGPRGDSGQPGPKGDP  265
col  105 FDGIQGPPGPKGDPGAFGLKGEKGEPGADGE----AGRPGSSGPSGDEGQPGEPGPPGEKGEAGDEGNPGPDGAPGERGGPGERGPRGTPGTRGPRGDPGEAGPQGDQGREGPVGVPGDPGEAGPIGPKGYRGDEGPPGSEGARGAPGPAGPPGDPGLMGERGED  265

col 1651 GSPGYGSVGRKGAKGQEGFPGESGPKGEIGDPGGPGETGLKGARGKMISAGLPGEMGSPGEPGPPGRKGVKGAKGLASFSTCELIQYVRDRSPGRHGKPECPVHPTELVFALDHSRDVTEQEFERMKEMMAFLVRDIKVRENSCPVGAHIAILSYNSHARHLVRF 1815
col  266 GRPGFSYPGPRGAPGEKGEPGPRGPEGGRGDFGLKGEPGRKGEKGEPADPGPP-GEPGPRGPRGVPGPEGEPGPPGDP---------------------------------------------------------------------------------------  342
col  266 GPAGNGTEGFPGFPGYPGNRGAPGINGTKGYPGLKGDEGEAGDPGDDNNDIAPRGVKGAKGYRGPEGPQGPPGHQGPP---------------------------------------------------------------------------------------  343

col 1816 SDAYKKSQLLREIETIPYERSSASREIGRAMRFISRNVFKRTLPGAHTRKIATFFSSGQSADAHSITTAAMEFGALEIIPVVITFSNVPSVRRAFAIDDTGTFQVIVVPSGADYIPALERLQRCTFCYDVCKPDASCDQARPPPVQSYMDAAFLLDASRNMGSAE 1980
col      ---------------------------------------------------------------------------------------------------------------------------------------------------------------------
col      ------------------------------------------------------------------------------------------------------------------------------------

col 1981 FEDIRAFLGALLDHFEITPEPETSVTGDRVALLSHAPPDFLPNTQKSPVRAEFNLTTYRSKRLMKRHVHESVKQLNGDAFIGHALQWTLDNVFLSTPNLRRNKVIFVISAGETSHLDGEILKKESLRAKCQGYALFVFSLGPIWDDKELEDLASHPLDHHLVQLG 2145
col      ---------------------------------------------------------------------------------------------------------------------------------------------------------------------
col

col 2146 RIHKPDHSYGVKFVKSFINSIRRAINKYPPINLKIKCNRLNSIDPKQPPRPFRSFVPGPLKATLKEDVLQKAKFFQDKKYLSRVARSGRDDAIQNFMRSTSHTFKNGRMIESAPKQHD                                               2263
col      ----------------------------------------------------------------------------------------------------------------------
col
```

#### collagen type XXVIII – (α1)$_3$

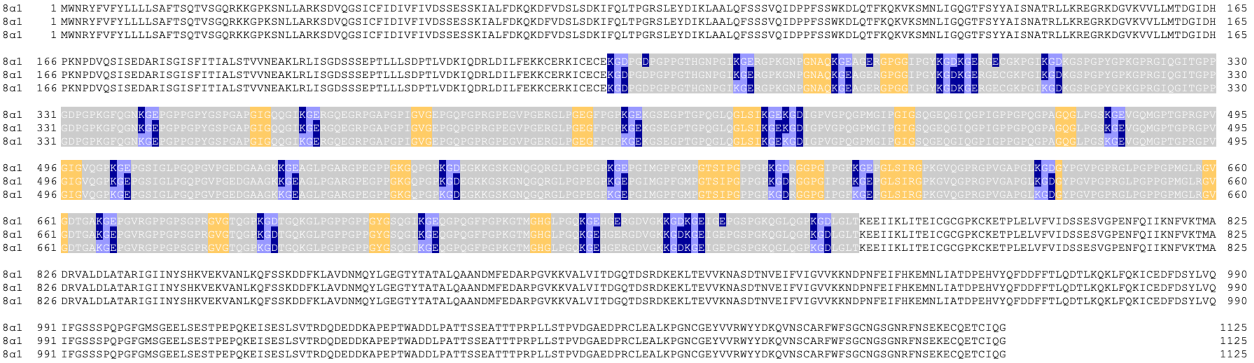

#### anchoring-fibril collagens

#### collagen type VII – (α1)$_3$

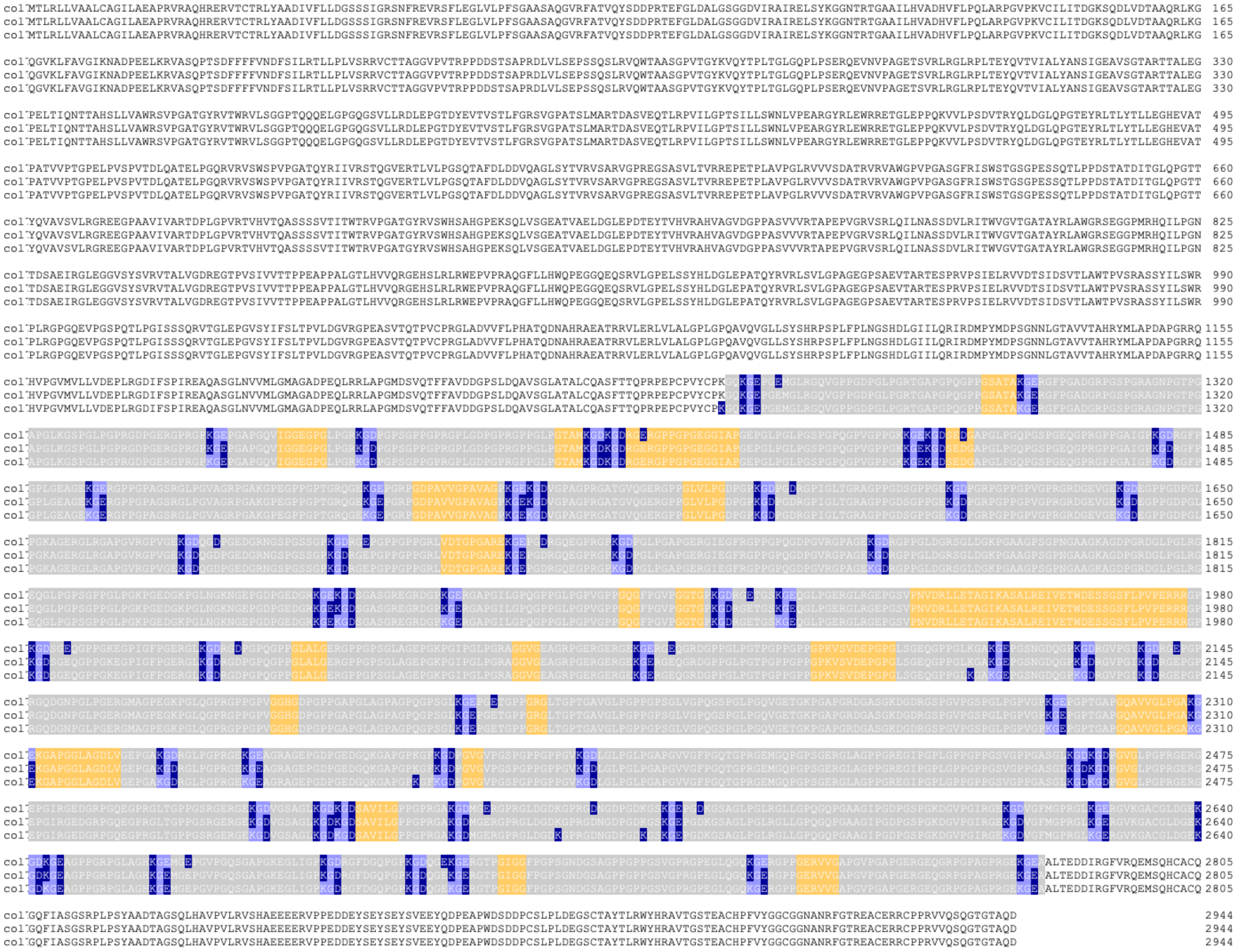

<u>transmembrane collagens</u>

#### collagen type XIII – $(\alpha 1)_3$

```
col    1 MVAERTHKAAATGARGPGELGAPGTVALVAARAERGARLPSPGSCGLLTLALCSLALSLLAHFRTAELQARVLRLEAERGEQQMETAILGRVNQLLDEKWKLHSRRRREAPKTSPGCNCPPGPPGPTGRPGLPGDKGAIGMPGRVGSPGDAGLSIIGPRGPPGQP 165
col    1 MVAERTHKAAATGARGPGELGAPGTVALVAARAERGARLPSPGSCGLLTLALCSLALSLLAHFRTAELQARVLRLEAERGEQQMETAILGRVNQLLDEKWKLHSRRRREAPKTSPGCNCPPGPPGPTGRPGLPGDKGAIGMPGRVGSPGDAGLSIIGPRGPPGQP 165
col    1 MVAERTHKAAATGARGPGELGAPGTVALVAARAERGARLPSPGSCGLLTLALCSLALSLLAHFRTAELQARVLRLEAERGEQQMETAILGRVNQLLDEKWKLHSRRRREAPKTSPGCNCPPGPPGPTGRPGLPGDKGAIGMPGRVGSPGDAGLSIIGPRGPPGQP 165

col  166 GTRGFPGFPGPIGLDGKPGHPGPKGDMGLTGPPGQPGPQGQKGEKGQCGEYPHRECLSSMPAALRSSQIIALKLLPLLNSVRLAPPPVIKRRTFQGEQSQASIQGPPGPPGPPGPSGPLGHPGLPGPMGPPGLPGPPGPKGDPGIQGYHGRKGERGMPGMPGKHG 330
col  166 GTRGFPGFPGPIGLDGKPGHPGPKGDMGLTGPPGQPGPQGQKGEKGQCGEYPHRECLSSMPAALRSSQIIALKLLPLLNSVRLAPPPVIKRRTFQGEQSQASIQGPPGPPGPPGPSGPLGHPGLPGPMGPPGLPGPPGPKGDPGIQGYHGRKGERGMPGMPGKHG 330
col  166 GTRGFPGFPGPIGLDGKPGHPGPKGDMGLTGPPGQPGPQGQKGEKGQCGEYPHRECLSSMPAALRSSQIIALKLLPLLNSVRLAPPPVIKRRTFQGEQSQASIQGPPGPPGPPGPSGPLGHPGLPGPMGPPGLPGPPGPKGDPGIQGYHGRKGERGMPGMPGKHG 330

col  331 AKGAPGIAVAGMKGEPGIPGTKGEKGAEGSPGLPGLLGQKGEKGDAGNSIGGGRGEPGPPGLPGPPGPKGEAGVDGQVGPPGQPGDKGERGAAGEQGPDGPKGSKGEPGKGEMVDYNGNINEALQEIRTLALMGPPGLPGQIGPPGAPGIPGQKGEIGLPGPPGH 495
col  331 AKGAPGIAVAGMKGEPGIPGTKGEKGAEGSPGLPGLLGQKGEKGDAGNSIGGGRGEPGPPGLPGPPGPKGEAGVDGQVGPPGQPGDKGERGAAGEQGPDGPKGSKGEPGKGEMVDYNGNINEALQEIRTLALMGPPGLPGQIGPPGAPGIPGQKGEIGLPGPPGH 495
col  331 AKGAPGIAVAGMKGEPGIPGTKGEKGAEGSPGLPGLLGQKGEKGDAGNSIGGGRGEPGPPGLPGPPGPKGEAGVDGQVGPPGQPGDKGERGAAGEQGPDGPKGSKGEPGKGEMVDYNGNINEALQEIRTLALMGPPGLPGQIGPPGAPGIPGQKGEIGLPGPPGH 495

col  496 DGEKGPRGKPGDMGPPGPQGPPGKDGPPGVKGENGHPGSPGEKGEKGETGQAGSPGEKGEAGEKGNPGAEVPGLPGPEGPPGPPGLQGVPGPKGEAGLDGAKGEKGFQGEKGDRGPLGLPGASGLDGRPGPPGTPGPIGVPGPAGPKGERGSKGDPGMTGPTGAA 660
col  496 DGEKGPRGKPGDMGPPGPQGPPGKDGPPGVKGENGHPGSPGEKGEKGETGQAGSPGEKGEAGEKGNPGAEVPGLPGPEGPPGPPGLQGVPGPKGEAGLDGAKGEKGFQGEKGDRGPLGLPGASGLDGRPGPPGTPGPIGVPGPAGPKGERGSKGDPGMTGPTGAA 660
col  496 DGEKGPRGKPGDMGPPGPQGPPGKDGPPGVKGENGHPGSPGEKGEKGETGQAGSPGEKGEAGEKGNPGAEVPGLPGPEGPPGPPGLQGVPGPKGEAGLDGAKGEKGFQGEKGDRGPLGLPGASGLDGRPGPPGTPGPIGVPGPAGPKGERGSKGDPGMTGPTGAA 660

col  661 GLPGLHGPPGDKGNRGERGKKGSRGPKGDKGDQGAPGLDAPCPLGEDGLPVQGCWN 716
col  661 GLPGLHGPPGDKGNRGERGKKGSRGPKGDKGDQGAPGLDAPCPLGEDGLPVQGCWNK 717
col  661 GLPGLHGPPGDKGNRGERGKKGSRGPKGDKGDQGAPGLDAPCPLGEDGLPVQGCWNKK 718
```

#### collagen type XVII – $(\alpha 1)_3$

```
col    1 MDVTKKNKRDGTEVTERIVTETVTTRLTSLPPKGGTSNGYAKTASLGGGSRLEKQSLTHGSSGYINSTGSTRGHASTSSYRRAHSPASTLPNSPGSTFERKTHVTRHAYEGSSSGNSSPEYPRKEFASSSTRGRSQTRESEIRVRLQSASPSTRWTELDDVKRLL 165
col    1 MDVTKKNKRDGTEVTERIVTETVTTRLTSLPPKGGTSNGYAKTASLGGGSRLEKQSLTHGSSGYINSTGSTRGHASTSSYRRAHSPASTLPNSPGSTFERKTHVTRHAYEGSSSGNSSPEYPRKEFASSSTRGRSQTRESEIRVRLQSASPSTRWTELDDVKRLL 165
col    1 MDVTKKNKRDGTEVTERIVTETVTTRLTSLPPKGGTSNGYAKTASLGGGSRLEKQSLTHGSSGYINSTGSTRGHASTSSYRRAHSPASTLPNSPGSTFERKTHVTRHAYEGSSSGNSSPEYPRKEFASSSTRGRSQTRESEIRVRLQSASPSTRWTELDDVKRLL 165

col  166 KGSRSASVSPTRNSSNTLPIPKKGTVETKIVTASSQSVSGTYDATILDANLPSHVWSSTLPAGSSMGTYHNNMTTQSSSLLNTNAYSAGSVFGVPNNMASCSPTLHPGLSTSSSVFGMQNNLAPSLTTLSHGTTTTSTAYGVKKNMPQSPAAVNTGVSTSAACTT 330
col  166 KGSRSASVSPTRNSSNTLPIPKKGTVETKIVTASSQSVSGTYDATILDANLPSHVWSSTLPAGSSMGTYHNNMTTQSSSLLNTNAYSAGSVFGVPNNMASCSPTLHPGLSTSSSVFGMQNNLAPSLTTLSHGTTTTSTAYGVKKNMPQSPAAVNTGVSTSAACTT 330
col  166 KGSRSASVSPTRNSSNTLPIPKKGTVETKIVTASSQSVSGTYDATILDANLPSHVWSSTLPAGSSMGTYHNNMTTQSSSLLNTNAYSAGSVFGVPNNMASCSPTLHPGLSTSSSVFGMQNNLAPSLTTLSHGTTTTSTAYGVKKNMPQSPAAVNTGVSTSAACTT 330

col  331 SVQSDDLLHKDCKFLILEKDNTPAKKEMELLIMTKDSGKVFTASPASIAATSFSEDTLKKEKQAAYNADSGLKAEANGDLKTVSTKGKTTTADIHSYGSSGGGGSGGGGGVGGAGGGPWGPAPAWCPCGSCCSWWKWLLGLLLTWLLLLGLLFGLIALAEEVRKL 495
col  331 SVQSDDLLHKDCKFLILEKDNTPAKKEMELLIMTKDSGKVFTASPASIAATSFSEDTLKKEKQAAYNADSGLKAEANGDLKTVSTKGKTTTADIHSYGSSGGGGSGGGGGVGGAGGGPWGPAPAWCPCGSCCSWWKWLLGLLLTWLLLLGLLFGLIALAEEVRKL 495
col  331 SVQSDDLLHKDCKFLILEKDNTPAKKEMELLIMTKDSGKVFTASPASIAATSFSEDTLKKEKQAAYNADSGLKAEANGDLKTVSTKGKTTTADIHSYGSSGGGGSGGGGGVGGAGGGPWGPAPAWCPCGSCCSWWKWLLGLLLTWLLLLGLLFGLIALAEEVRKL 495

col  496 KARVDELERIRRSILPYGDSMDRIEKDRLQGMAPAAGADLDKIGLHSDSQEELWMFVRKKLMMEQENGNLRGSPGPKGDMGSPGPKGDRGFPGTPGIPGPLGHPGPQGPKGQKGSVGDPGMEGPMGQRGREGPMGPRGEAGPPGSGEKGERGAAGEPGPHGPPGV 660
col  496 KARVDELERIRRSILPYGDSMDRIEKDRLQGMAPAAGADLDKIGLHSDSQEELWMFVRKKLMMEQENGNLRGSPGPKGDMGSPGPKGDRGFPGTPGIPGPLGHPGPQGPKGQKGSVGDPGMEGPMGQRGREGPMGPRGEAGPPGSGEKGERGAAGEPGPHGPPGV 660
col  496 KARVDELERIRRSILPYGDSMDRIEKDRLQGMAPAAGADLDKIGLHSDSQEELWMFVRKKLMMEQENGNLRGSPGPKGDMGSPGPKGDRGFPGTPGIPGPLGHPGPQGPKGQKGSVGDPGMEGPMGQRGREGPMGPRGEAGPPGSGEKGERGAAGEPGPHGPPGV 660

col  661 PGSVGPKGSSGSPGPQGPPGPVGLQGLRGEVGLPGVKGDKGPMGPPGPKGDQGEKGPRGLTGEPGMRGLPGAVGEPGAKGAMGPAGPDGHQGPRGEQGLTGMPGIRGPPGPSGDPGKPGLTGPQGPQGLPGTPGRPGIKGEPGAPGKIVTSEGSSMLTVPGPPGP 825
col  661 PGSVGPKGSSGSPGPQGPPGPVGLQGLRGEVGLPGVKGDKGPMGPPGPKGDQGEKGPRGLTGEPGMRGLPGAVGEPGAKGAMGPAGPDGHQGPRGEQGLTGMPGIRGPPGPSGDPGKPGLTGPQGPQGLPGTPGRPGIKGEPGAPGKIVTSEGSSMLTVPGPPGP 825
col  661 PGSVGPKGSSGSPGPQGPPGPVGLQGLRGEVGLPGVKGDKGPMGPPGPKGDQGEKGPRGLTGEPGMRGLPGAVGEPGAKGAMGPAGPDGHQGPRGEQGLTGMPGIRGPPGPSGDPGKPGLTGPQGPQGLPGTPGRPGIKGEPGAPGKIVTSEGSSMLTVPGPPGP 825

col  826 PGAMGPPGPPGAPGPAGPAGLPGHQEVLNLQGPPGPPGPRGPPGPSIPGPPGPRGPPGEGLPGPPGPPGSFLSNSETFLSGPPGPPGPPGPKGDQGPPGPRGHQGEQGLPGFSTSGSSSFGLNLQGPPGPPGPQGPKGDKGDPGVPGALGIPSGPSEGGSSSTMY 990
col  826 PGAMGPPGPPGAPGPAGPAGLPGHQEVLNLQGPPGPPGPRGPPGPSIPGPPGPRGPPGEGLPGPPGPPGSFLSNSETFLSGPPGPPGPPGPKGDQGPPGPRGHQGEQGLPGFSTSGSSSFGLNLQGPPGPPGPQGPKGDKGDPGVPGALGIPSGPSEGGSSSTMY 990
col  826 PGAMGPPGPPGAPGPAGPAGLPGHQEVLNLQGPPGPPGPRGPPGPSIPGPPGPRGPPGEGLPGPPGPPGSFLSNSETFLSGPPGPPGPPGPKGDQGPPGPRGHQGEQGLPGFSTSGSSSFGLNLQGPPGPPGPQGPKGDKGDPGVPGALGIPSGPSEGGSSSTMY 990

col  991 VSGPPGPPGPPGPPGPPGSISSSGQEIQQYISEYMQSDSIRSYLSGVQGPPGPPGPPGPPGPVTTITGETFDYSELASHVVSYLRTSGYGVSLFSSSISSEDILAVLQRDDVRQYLRQYLMGPRGPPGPPGASGDGSLLSLDYAELSSRILSYMSSSGISIGLPGPPGPPG 1155
col  991 VSGPPGPPGPPGPPGPPGSISSSGQEIQQYISEYMQSDSIRSYLSGVQGPPGPPGPPGPPGPVTTITGETFDYSELASHVVSYLRTSGYGVSLFSSSISSEDILAVLQRDDVRQYLRQYLMGPRGPPGPPGASGDGSLLSLDYAELSSRILSYMSSSGISIGLPGPPGPPG 1155
col  991 VSGPPGPPGPPGPPGPPGSISSSGQEIQQYISEYMQSDSIRSYLSGVQGPPGPPGPPGPPGPVTTITGETFDYSELASHVVSYLRTSGYGVSLFSSSISSEDILAVLQRDDVRQYLRQYLMGPRGPPGPPGASGDGSLLSLDYAELSSRILSYMSSSGISIGLPGPPGPPG 1155

col 1156 LPGTSYEELLSLLRGSEFRGIVGPPGPPGPPGIPGNVWSSISVEDLSSYLHTAGLSFIPGPPGPPGPPGPRGPPGVSGALATYAAENSDSFRSELISYLTSPDVRSFIVGPPGPPGPQGPPGDSRLLSTDASHSRGSSSSSHSSSVRRGSSYSSSMSTGGGGAGS 1320
col 1156 LPGTSYEELLSLLRGSEFRGIVGPPGPPGPPGIPGNVWSSISVEDLSSYLHTAGLSFIPGPPGPPGPPGPRGPPGVSGALATYAAENSDSFRSELISYLTSPDVRSFIVGPPGPPGPQGPPGDSRLLSTDASHSRGSSSSSHSSSVRRGSSYSSSMSTGGGGAGS 1320
col 1156 LPGTSYEELLSLLRGSEFRGIVGPPGPPGPPGIPGNVWSSISVEDLSSYLHTAGLSFIPGPPGPPGPPGPRGPPGVSGALATYAAENSDSFRSELISYLTSPDVRSFIVGPPGPPGPQGPPGDSRLLSTDASHSRGSSSSSHSSSVRRGSSYSSSMSTGGGGAGS 1320

col 1321 LGAGGAFGEAAGDRGPYGTDIGPGGGYGAAAEGGMYAGNGGLLGADFAGDLDYNELAVRVSESMQRQGLLQGMAYTVQGPPGQPGPQGPPGISKVFSAYSNVTADLMDFFQTYGAIQGPPGQKGEMGTPGPKGDRGPAGPPGHPGPPGPRGHKGEKGDKGDQVYA 1485
col 1321 LGAGGAFGEAAGDRGPYGTDIGPGGGYGAAAEGGMYAGNGGLLGADFAGDLDYNELAVRVSESMQRQGLLQGMAYTVQGPPGQPGPQGPPGISKVFSAYSNVTADLMDFFQTYGAIQGPPGQKGEMGTPGPKGDRGPAGPPGHPGPPGPRGHKGEKGDKGDQVYA 1485
col 1321 LGAGGAFGEAAGDRGPYGTDIGPGGGYGAAAEGGMYAGNGGLLGADFAGDLDYNELAVRVSESMQRQGLLQGMAYTVQGPPGQPGPQGPPGISKVFSAYSNVTADLMDFFQTYGAIQGPPGQKGEMGTPGPKGDRGPAGPPGHPGPPGPRGHKGEKGDKGDQVYA 1485

col 1486 GRRRRRSIAVKP 1497
col 1486 GRRRRRSIAVKP 1497
col 1486 GRRRRRSIAVKP 1497
```

#### collagen type XXIII – $(\alpha 1)_3$

```
col    1 MGPGERAGGGDAGKGNAAGGGGGGRSATTAGSRAVSALCLLLSVGSAAACLLLGVQAAALQGRVAALEEERELLRRAGPPGALDAWAEPHLERLLREKLDGLAKIRTAREAPSECVCPPGPPGRRGKPGRRGDPGPPGQSGRDGYPGPLGLDGKPGLPGPKGEK 165
col    1 MGPGERAGGGDAGKGNAAGGGGGGRSATTAGSRAVSALCLLLSVGSAAACLLLGVQAAALQGRVAALEEERELLRRAGPPGALDAWAEPHLERLLREKLDGLAKIRTAREAPSECVCPPGPPGRRGKPGRRGDPGPPGQSGRDGYPGPLGLDGKPGLPGPKGEK 165
col    1 MGPGERAGGGDAGKGNAAGGGGGGRSATTAGSRAVSALCLLLSVGSAAACLLLGVQAAALQGRVAALEEERELLRRAGPPGALDAWAEPHLERLLREKLDGLAKIRTAREAPSECVCPPGPPGRRGKPGRRGDPGPPGQSGRDGYPGPLGLDGKPGLPGPKGEK 165

col  166 GAPGDFGPRGDQGQDGAAGPPGPPGPPGARGPPGDTGKDGPRGAQGPAGPKGEPGQDGEMGPKGPPGPKGEPGVPGKKGDDGTPSQPGPPGPKGEPGSMGPRGENGVDGAPGPKGEPGHRGTDGAAGPRGAPGLKGEQGDTVVIDYDGRILDALKGPPGPQGPPG 330
col  166 GAPGDFGPRGDQGQDGAAGPPGPPGPPGARGPPGDTGKDGPRGAQGPAGPKGEPGQDGEMGPKGPPGPKGEPGVPGKKGDDGTPSQPGPPGPKGEPGSMGPRGENGVDGAPGPKGEPGHRGTDGAAGPRGAPGLKGEQGDTVVIDYDGRILDALKGPPGPQGPPG 330
col  166 GAPGDFGPRGDQGQDGAAGPPGPPGPPGARGPPGDTGKDGPRGAQGPAGPKGEPGQDGEMGPKGPPGPKGEPGVPGKKGDDGTPSQPGPPGPKGEPGSMGPRGENGVDGAPGPKGEPGHRGTDGAAGPRGAPGLKGEQGDTVVIDYDGRILDALKGPPGPQGPPG 330

col  331 PPGIPGAKGELGLPGAPGIDGEKGPKGQKGDPGEPGPAGIKGEAGEMGLSGLPGADGLKGEKGESASDSLQESLAQLIVEPGPPGPPGPPGPMGLQGIQGPKGLDGAKGEKGASGERGPSGLPGPVGPPGLIGLPGTKGEKGRPGEPGLDGFPGPRGEKGDRSER 495
col  331 PPGIPGAKGELGLPGAPGIDGEKGPKGQKGDPGEPGPAGIKGEAGEMGLSGLPGADGLKGEKGESASDSLQESLAQLIVEPGPPGPPGPPGPMGLQGIQGPKGLDGAKGEKGASGERGPSGLPGPVGPPGLIGLPGTKGEKGRPGEPGLDGFPGPRGEKGDRSER 495
col  331 PPGIPGAKGELGLPGAPGIDGEKGPKGQKGDPGEPGPAGIKGEAGEMGLSGLPGADGLKGEKGESASDSLQESLAQLIVEPGPPGPPGPPGPMGLQGIQGPKGLDGAKGEKGASGERGPSGLPGPVGPPGLIGLPGTKGEKGRPGEPGLDGFPGPRGEKGDRSER 495

col  496 GEKGERGVPGRKGVKGQKGEPGPPGLDQPCPVGPDGLPVPGCWHK 540
col  496 GEKGERGVPGRKGVKGQKGEPGPPGLDQPCPVGPDGLPVPGCWHK 540
col  496 GEKGERGVPGRKGVKGQKGEPGPPGLDQPCPVGPDGLPVPGCWHK 540
```

#### collagen type XXV – $(\alpha 1)_3$

```
col    1 MLLKKHAGKGGGREPRSEDPTPAEQHCARTMPPCAVLAALLSVVAVVSCLYLGVKTNDLQARIAALESAKGAPSIHLLPDTLDHLKTMVQEKVERLLAQKSYEHMAKIRIAREAPSECNCPAGPPGKRGKRGRRGESGPPGQPGPQGPPGPKGDKGEQGDQGPRM 165
col    1 MLLKKHAGKGGGREPRSEDPTPAEQHCARTMPPCAVLAALLSVVAVVSCLYLGVKTNDLQARIAALESAKGAPSIHLLPDTLDHLKTMVQEKVERLLAQKSYEHMAKIRIAREAPSECNCPAGPPGKRGKRGRRGESGPPGQPGPQGPPGPKGDKGEQGDQGPRM 165
col    1 MLLKKHAGKGGGREPRSEDPTPAEQHCARTMPPCAVLAALLSVVAVVSCLYLGVKTNDLQARIAALESAKGAPSIHLLPDTLDHLKTMVQEKVERLLAQKSYEHMAKIRIAREAPSECNCPAGPPGKRGKRGRRGESGPPGQPGPQGPPGPKGDKGEQGDQGPRM 165

col  166 VFPKINHGFLSADQQLIKRRLIKGDQGQAGPPGPPGPPGPRGPPGDTGKDGPRGMPGVPGEPGKPGEQGLMGPLGPPGQKGSIGAPGIPGMNGQKGEPGLPGAVGQNGIPGPKGEPGEQGEKGDAGENGPKGDTGEKGDPGSSAAGIKGEPGESGRPGQKGEPGL 330
col  166 VFPKINHGFLSADQQLIKRRLIKGDQGQAGPPGPPGPPGPRGPPGDTGKDGPRGMPGVPGEPGKPGEQGLMGPLGPPGQKGSIGAPGIPGMNGQKGEPGLPGAVGQNGIPGPKGEPGEQGEKGDAGENGPKGDTGEKGDPGSSAAGIKGEPGESGRPGQKGEPGL 330
col  166 VFPKINHGFLSADQQLIKRRLIKGDQGQAGPPGPPGPPGPRGPPGDTGKDGPRGMPGVPGEPGKPGEQGLMGPLGPPGQKGSIGAPGIPGMNGQKGEPGLPGAVGQNGIPGPKGEPGEQGEKGDAGENGPKGDTGEKGDPGSSAAGIKGEPGESGRPGQKGEPGL 330

col  331 PGLPGLPGIKGEPGFIGPQGEPGLPGLPGTKGERGEAGPPGRGERGEPGAPGPKGKQGESGTRGPKGSKGDRGEKGDSGAQGPRGPPGQKGDQGATEIIDYNGNLHEALQRITTLTVTGPPGPPGPQGLQGPKGEQGSPGIPGMDGEQGLKGSKGDMGDPGMTGE 495
col  331 PGLPGLPGIKGEPGFIGPQGEPGLPGLPGTKGERGEAGPPGRGERGEPGAPGPKGKQGESGTRGPKGSKGDRGEKGDSGAQGPRGPPGQKGDQGATEIIDYNGNLHEALQRITTLTVTGPPGPPGPQGLQGPKGEQGSPGIPGMDGEQGLKGSKGDMGDPGMTGE 495
col  331 PGLPGLPGIKGEPGFIGPQGEPGLPGLPGTKGERGEAGPPGRGERGEPGAPGPKGKQGESGTRGPKGSKGDRGEKGDSGAQGPRGPPGQKGDQGATEIIDYNGNLHEALQRITTLTVTGPPGPPGPQGLQGPKGEQGSPGIPGMDGEQGLKGSKGDMGDPGMTGE 495

col  496 KGGIGLPGLPGANGMKGEKGDSGMPGPQGPSIIGPPGPPGPHGPPGPMGPHGLPGPKGTDGPMGPHGPAGPKGERGEKGAMGEPGPRGPYGLPGKDGEPGLDGFPGPRGEKGDLGEKGEKGFRGVKGEKGEPGQPGLDGLDAPCQLGPDGLPMPGCWQK 654
col  496 KGGIGLPGLPGANGMKGEKGDSGMPGPQGPSIIGPPGPPGPHGPPGPMGPHGLPGPKGTDGPMGPHGPAGPKGERGEKGAMGEPGPRGPYGLPGKDGEPGLDGFPGPRGEKGDLGEKGEKGFRGVKGEKGEPGQPGLDGLDAPCQLGPDGLPMPGCWQK 654
col  496 KGGIGLPGLPGANGMKGEKGDSGMPGPQGPSIIGPPGPPGPHGPPGPMGPHGLPGPKGTDGPMGPHGPAGPKGERGEKGAMGEPGPRGPYGLPGKDGEPGLDGFPGPRGEKGDLGEKGEKGFRGVKGEKGEPGQPGLDGLDAPCQLGPDGLPMPGCWQK 654
```

#### multiplexin collagens

collagen type XV – (α1)$_3$

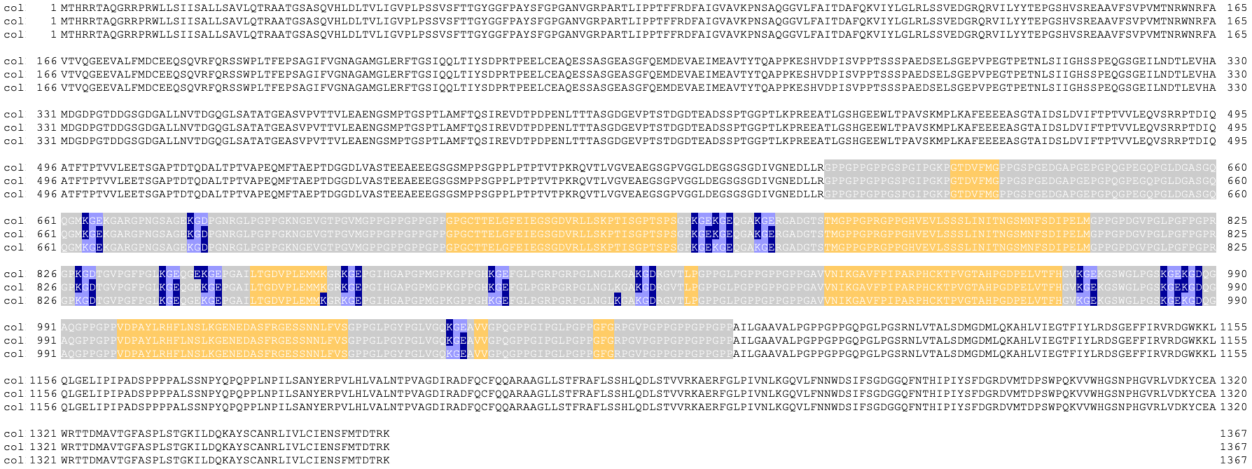

collagen type XVIII – (α1)$_3$

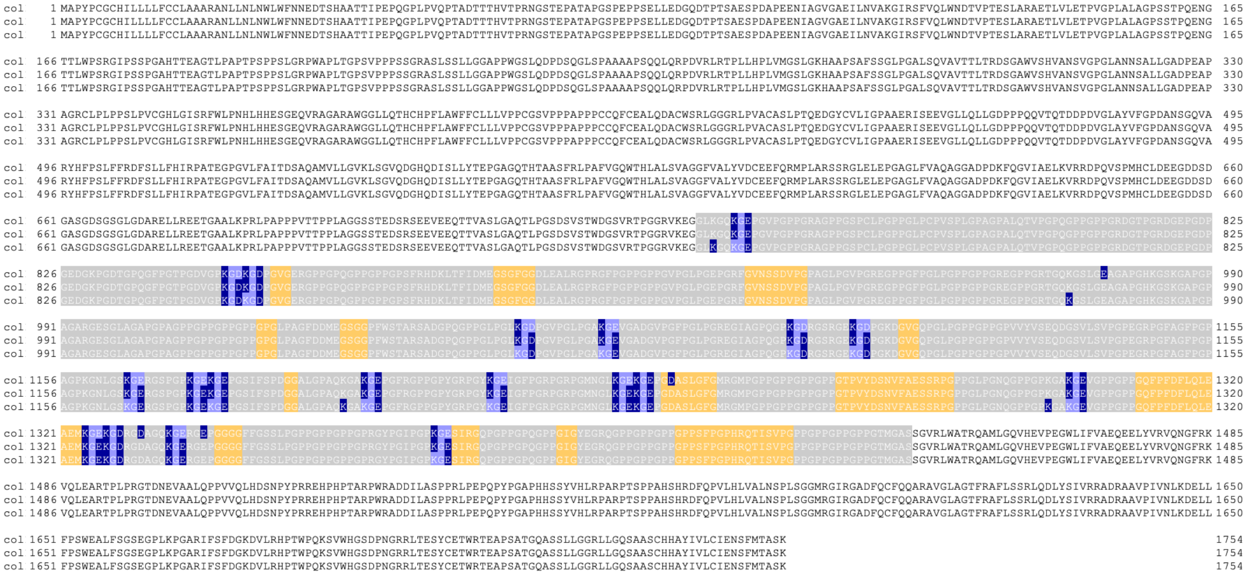

**Supplementary figure S1.** Schematic illustration of the triple-helical domains (gray), KGE or KGD triplets (light purple), salt-bridges (dark blue) and interruptions (yellow) in the aligned triple helical stoichiometries of human collagens.

**Supplementary table S2. Abundance of salt-bridges and interruptions in human collagens.** The number of axial lysine-aspartate and lysine-glutamate salt-bridges present in the twenty-eight known human collagens in the native chain alignment and when intentionally misaligned by 3 residues of chain B or C. The collagen assemblies where the native chain alignment is unknown and thus the number salt-bridges cannot be obtained are indicated with dash (-). Collagen assemblies containing a single uninterrupted triple-helical domain are indicated with asterik (*).

| Hierarchical structure | Type of Collagen | Stoichiometry | Salt-bridges | Salt-bridges (middle chain misaligned) | Salt-bridges (trailing chain misaligned) | interruptions | interruptions flanked by salt-bridge knots |
|---|---|---|---|---|---|---|---|
| **Fibers** | I | $(\alpha1)_3^{a}$ | 33 | 22 | 20 | 4 | 2 |
| | I | $(\alpha1)_2\alpha2^{a}$ | 30 | 18 | 18 | 7 | 1 |
| | II | $(\alpha1)_3^{a}$ | 36 | 28 | 25 | 5 | 1 |
| | III | $(\alpha1)_3^{a}$ | 42 | 17 | 32 | 19 | 9 |
| | V | $(\alpha1)_3^{a}$ | 59 | 47 | 48 | 8 | 3 |
| | V | $(\alpha1)_2\alpha2^{a}$ | 44 | 45 | 36 | 13 | 7 |
| | V | $\alpha1\alpha2\alpha3^{a}$ | - | - | - | - | - |
| | XI | $\alpha1\alpha2\alpha3^{a}$ | - | - | - | - | - |
| | XI | $\alpha1\alpha1(V)\alpha3^{a}$ | - | - | - | - | - |
| | XXIV | $(\alpha1)_3^{a}$ | 40 | 31 | 31 | 6 | 4 |
| | XXVII | $(\alpha1)_3^{a}$ | 39 | 28 | 29 | 4 | 2 |
| **Networks** | IV | $(\alpha1)_3\alpha1$ | 129 | 92 | 81 | 36 | 23 |
| | IV | $(\alpha5)_2\alpha6$ | 100 | 58 | 62 | 29 | 18 |
| | IV | $\alpha3\alpha4\alpha5$ | 61 | 56 | 49 | 35 | 24 |
| | VIII | $(\alpha1)3^{a}$ | 22 | 2 | 15 | 14 | 5 |
| | VIII | $(\alpha2)3^{a}$ | 18 | 1 | 10 | 12 | 5 |
| | VIII | $(\alpha1)_2\alpha2^{a}$ | - | - | - | | |
| | VIII | $\alpha1(\alpha2)_2^{a}$ | - | - | - | | |
| | X | $(\alpha1)_3$ | 18 | 1 | 10 | 10 | 4 |
| **Fibril-associated collagen with interrupted triple-helices (FACIT)** | IX | $\alpha1\alpha2\alpha3$ | 19 | 12 | 19 | 8 | 3 |
| | XII | $(\alpha1)_3$ | 12 | 10 | 7 | 6 | 4 |
| | XIV | $(\alpha1)_3$ | 11 | 9 | 9 | 7 | 4 |
| | XVI | $(\alpha1)_3$ | 84 | 63 | 60 | 24 | 17 |
| | XIX | $(\alpha1)_3$ | 70 | 42 | 44 | 11 | 9 |
| | XX | $(\alpha1)_3$ | 10 | 6 | 7 | 1 | 1 |
| | XXI | $(\alpha1)_3$ | 35 | 17 | 28 | 2 | |
| | XXII | $(\alpha1)_3$ | 66 | 45 | 50 | 12 | 7 |
| **Microfibrils** | VI | $\alpha3\alpha2\alpha1^{a}$ | 30 | 18 | 21 | 13 | 5 |
| | VI | $\alpha5\alpha2\alpha1^{a}$ | 29 | 15 | 20 | 11 | 6 |
| | VI | $\alpha6\alpha2\alpha1^{a}$ | 33 | 19 | 22 | 11 | 5 |
| | XXVIII | $(\alpha1)_3^{a}$ | 60 | 22 | 37 | 18 | 13 |
| **Anchoring-fibrils** | VII | $(\alpha1)_3$ | 141 | 85 | 97 | 22 | 17 |
| **Transmembrane collagens** | XIII | $(\alpha1)_3$ | 57 | 32 | 45 | 6 | 5 |
| | XVII | $(\alpha1)_3$ | 33 | 18 | 24 | 7 | 2 |
| | XXIII | $(\alpha1)_3$ | 43 | 24 | 32 | 4 | 4 |
| | XXV | $(\alpha1)_3$ | 65 | 45 | 43 | 6 | 4 |
| **Multiplexins** | XV | $(\alpha1)_3$ | 35 | 16 | 25 | 9 | 6 |
| | XVIII | $(\alpha1)_3$ | 49 | 26 | 28 | 14 | 8 |
| Total | | | 1553 | 970 | 1084 | 394 | 228 |

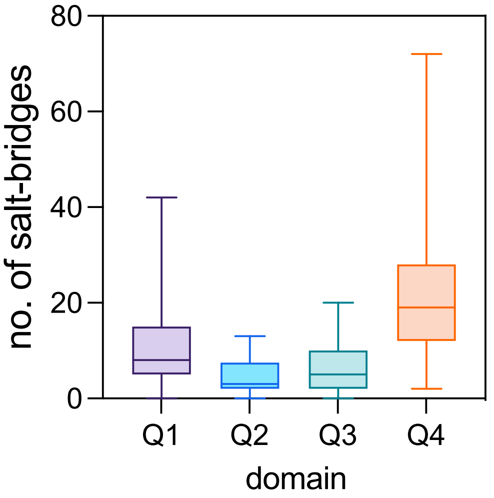


**Supplementary figure S2.** Distribution of salt-bridges in the four quarters (Q1-Q4) of the triple-helical domain of human collagens. Each quarter contains an equal number of triplets. Q1 represents the N-terminal most quarter.

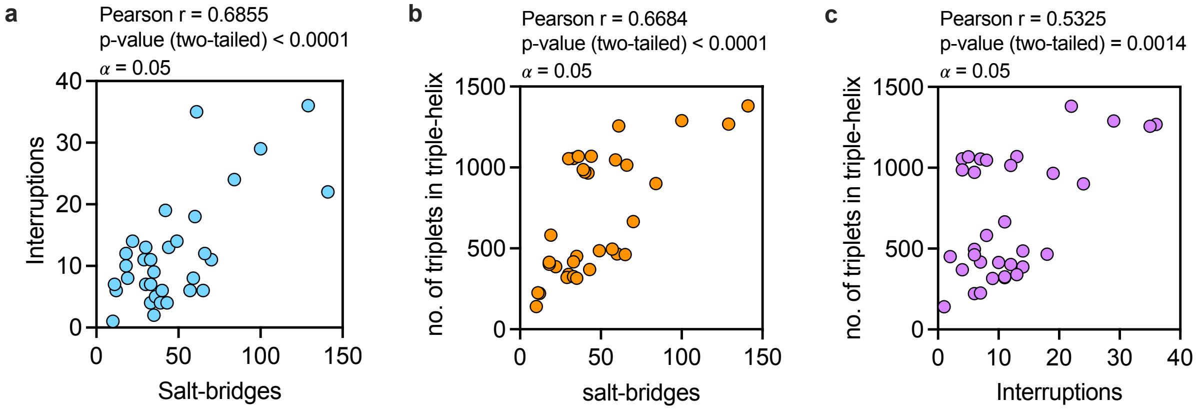


**Supplementary figure S3.** Correlation between the number of interruptions, salt-bridges and the length of triple-helical domain observed in the aligned collagen sequences show in supplementary figure S1.

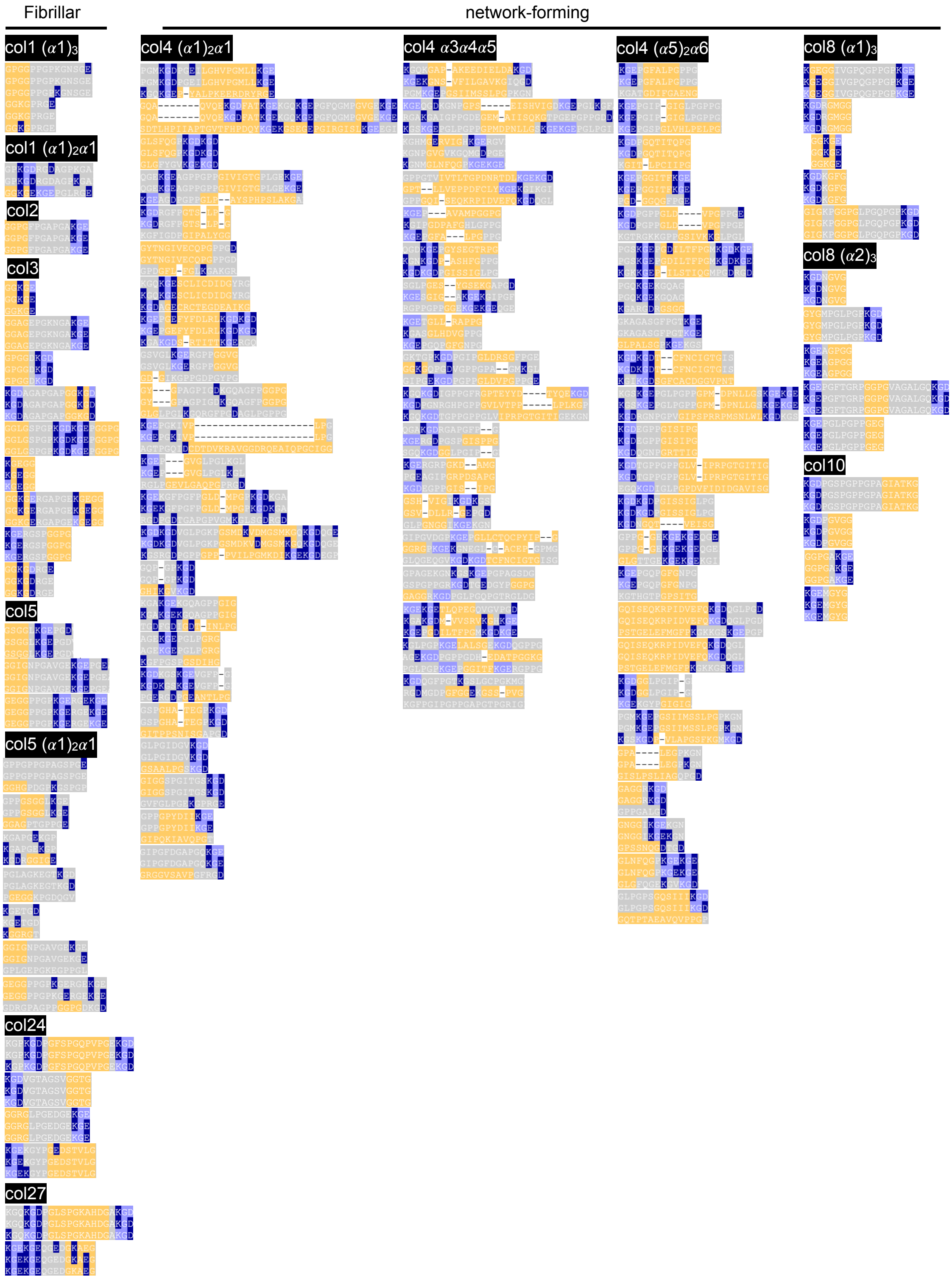
Fibrillar
network-forming
col1 (α1)3
col1 (α1)2α1
col2
col3
col5
col5 (α1)2α1
col24
col27
col4 (α1)2α1
col4 α3α4α5
col4 (α5)2α6
col8 (α1)3
col8 (α2)3
col10

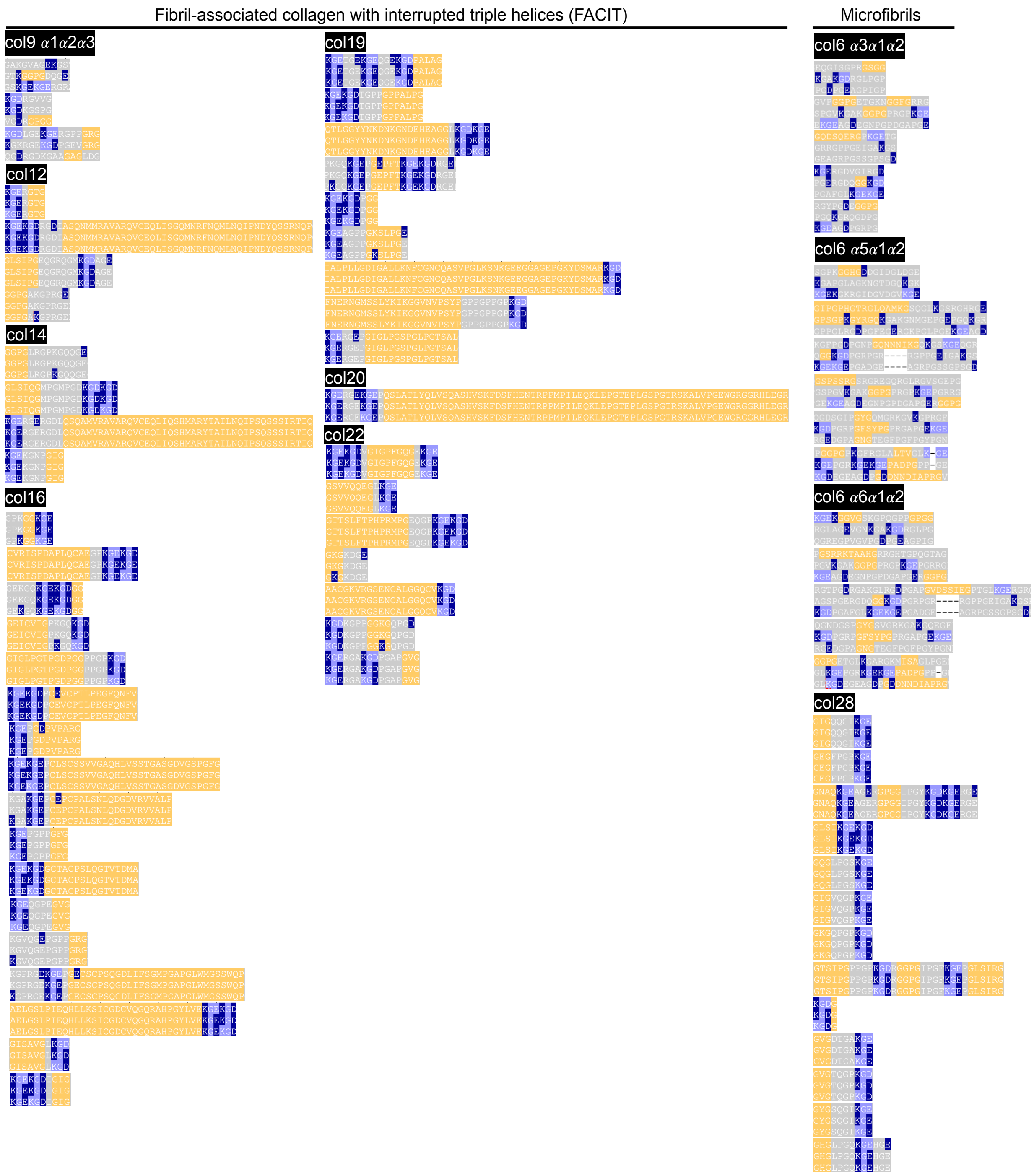
Fibril-associated collagen with interrupted triple helices (FACIT)
Microfibrils
col9 α1α2α3
col12
col14
col16
col19
col20
col22
col6 α3α1α2
col6 α5α1α2
col6 α6α1α2
col28

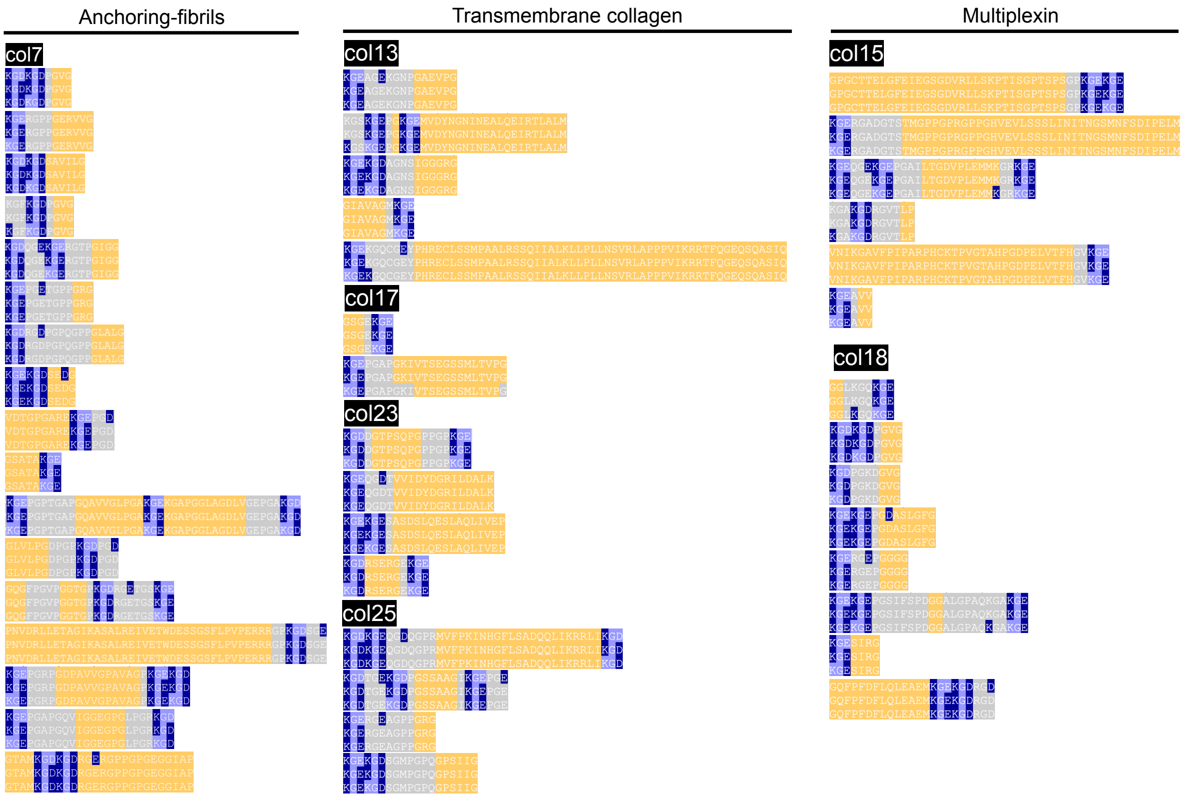


**Supplementary figure S4.** Interruptions in human collagens are frequently flanked by salt-bridge knots. KGE and KGD triplets (light blue) and lysine – aspartate/glutamate salt-bridges (dark blue) observed surrounding the interruptions (orange) in aligned triple-helical domains of human collagens.

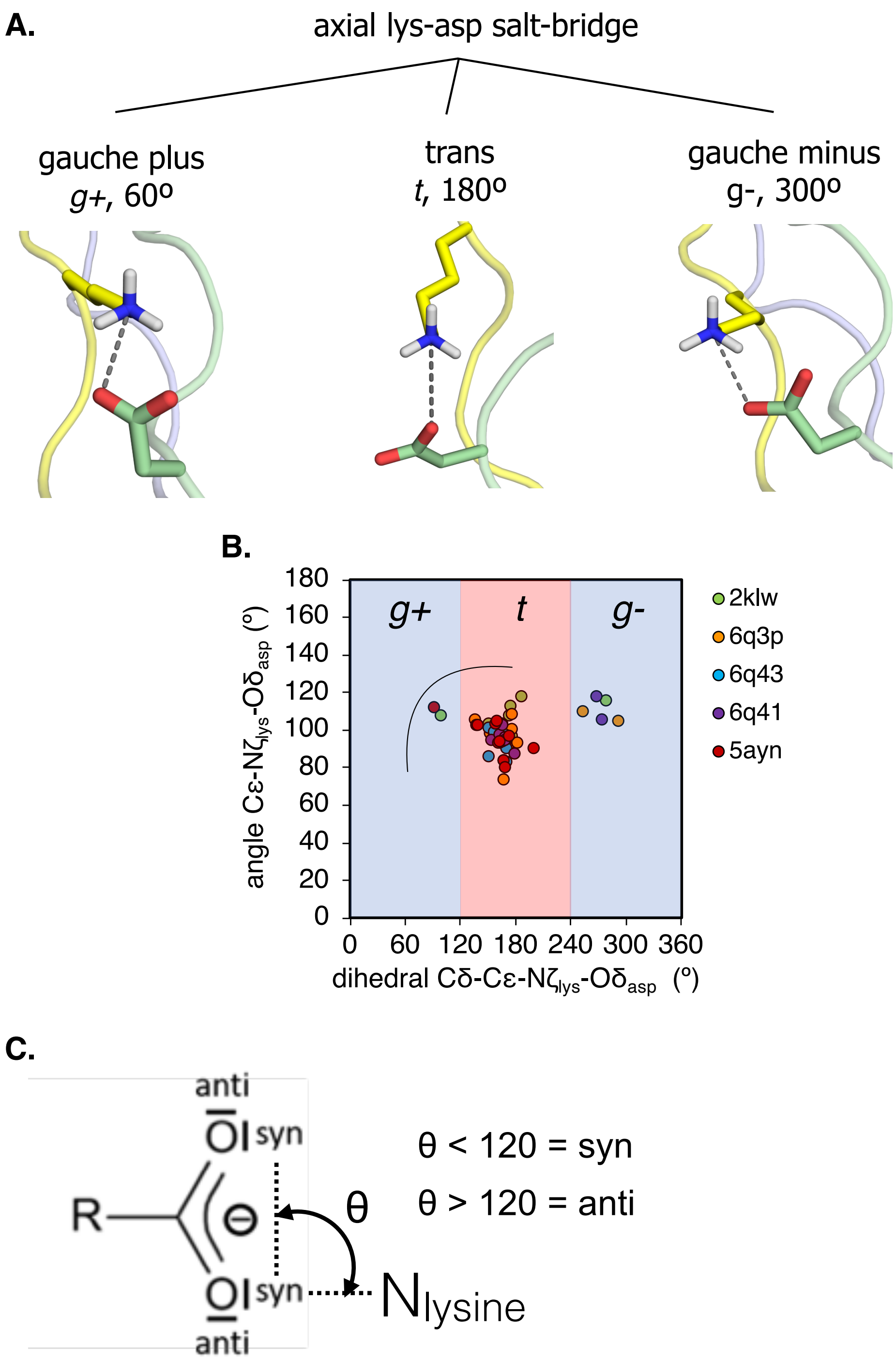


**Supplementary figure S5.** (A) Geometrical preference of the lysine-aspartate/glutamate axial salt-bridges parameterized by the angle defined by the carboxylate oxygen and the Cε-Nζ atoms of lysine and the dihedral angle defined by the carboxylate oxygen and Cδ-Cε-Nζ atoms of lysine (A) and the angle of these interactions observed in the previously published crystal structures of collagen peptides deposited under pdb codes 2klw, 6q3p, 6q43, 6q41 and 5ayn (B). Criteria for classification of lys – asp/glu salt-bridges *syn* or *anti* (**supporting data 3**).

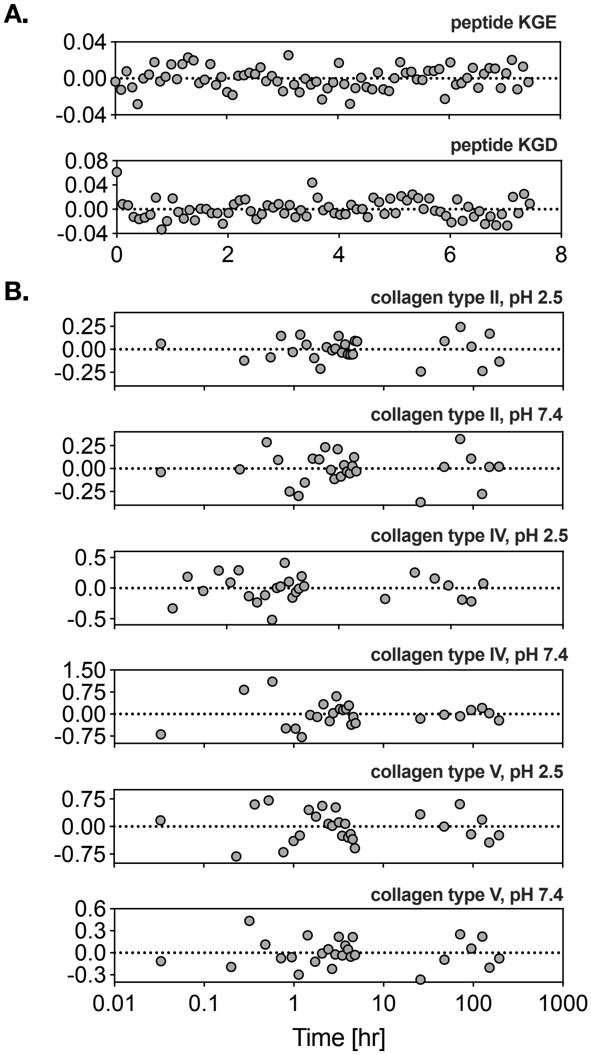


**Supplementary figure S6.** Residuals for monoexponential fit of the unfolding kinetics of collagen peptides KGE and KGD (A) and biexponential fit of the unfolding kinetics of native collagens type II, IV and V (B).

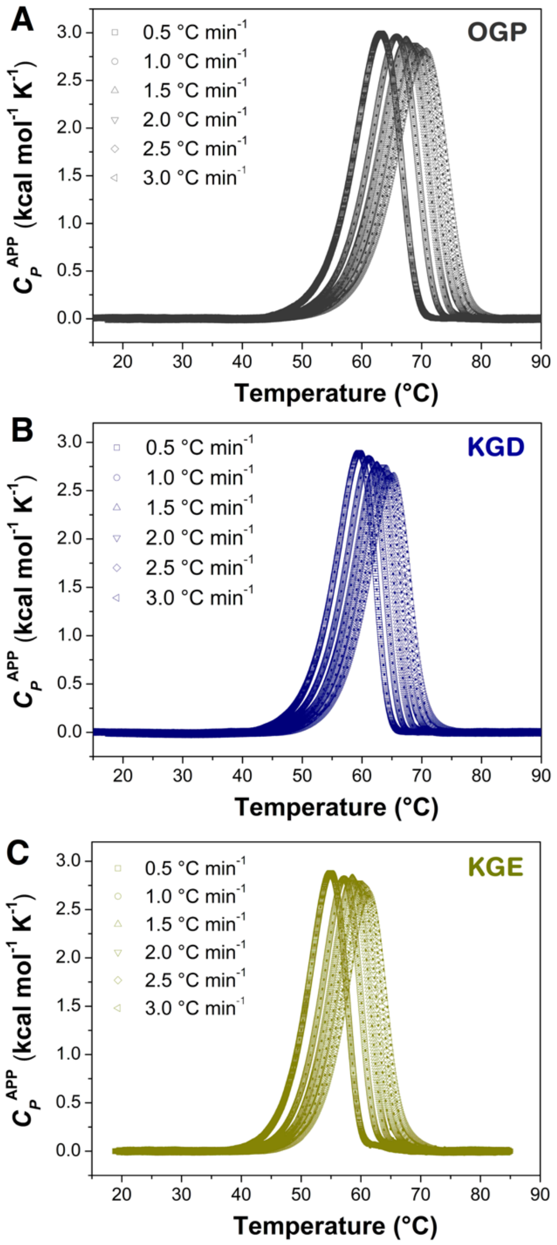


**Supplementary figure S7.** Thermal unfolding experiments by DSC for the different peptides. DSC endotherms at different scan rates (0.5-3.0 ºC/min) were collected for GPO (A), KGD (B), and KGE (C). Lines represent the best fit to a two-state irreversible model. In all experiments, physical and chemical baselines were subtracted and peptide concentration was 0.4 mg/mL in buffer 10 mM sodium phosphate pH 7.0.

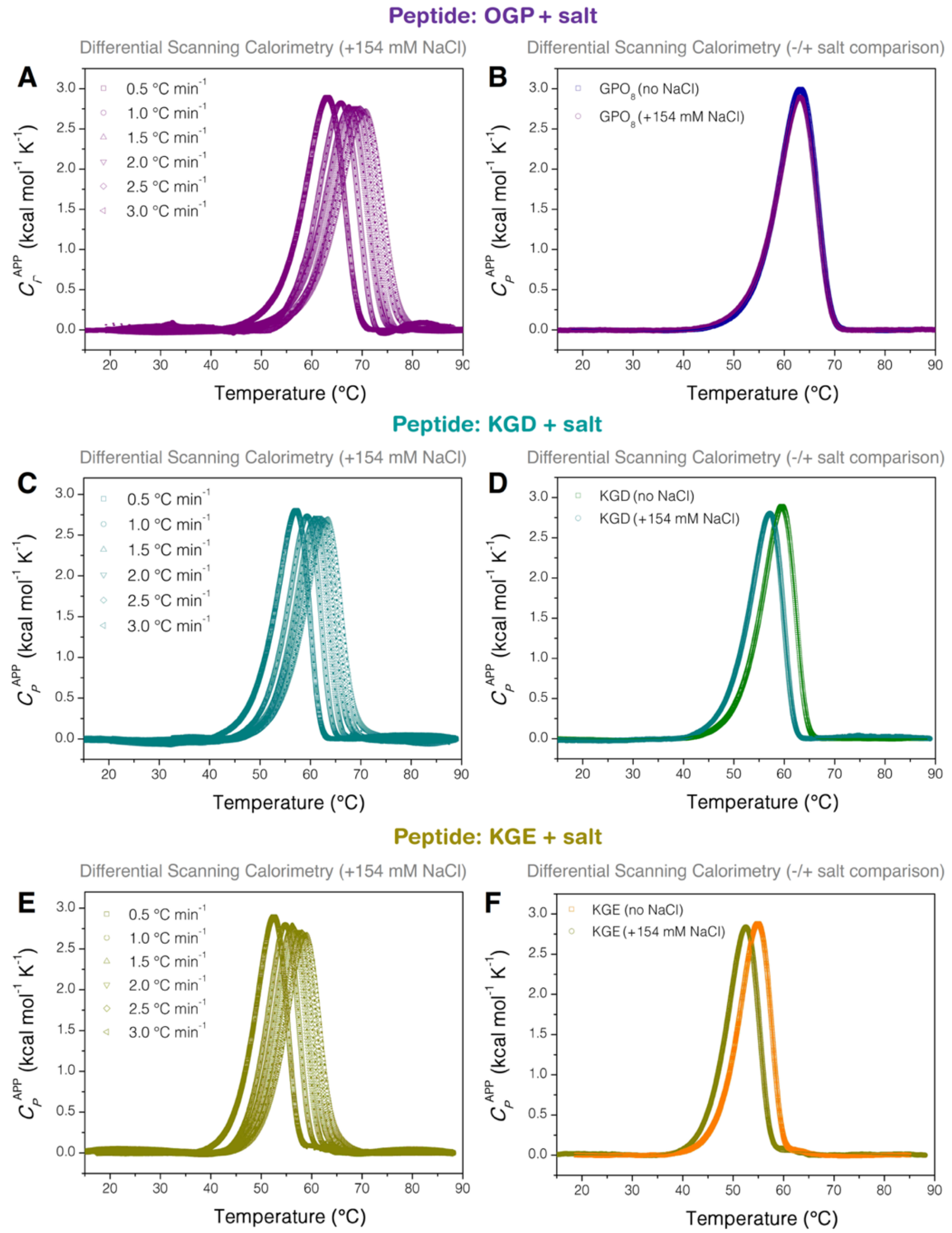


**Supplementary figure S8.** Effect of salt in the thermal-unfolding kinetic stability for the different peptides. DSC endotherms at different scan rates (0.5-3.0 ºC/min) were collected for GPO (A), KGD (C), and KGE (E) in the presence of 10 mM sodium phosphate pH 7.0 and 154 mM NaCl. Lines represent the best fit to a two-state irreversible model. Panels B), D) and F) show a comparison of the thermal-unfolding profiles in the absence and presence of salt at a constant scan rate (0.5 ºC/min). In all endotherms, physical and chemical baselines were substracted and peptide concentration was 0.4 mg/mL.

**Supplementary table S3.** Melting temperature ($T_m$) and activation energy ($E_a$) for the thermal unfolding of peptides OGP, KGD and KGE with and without physiological concentration of sodium chloride. Please see methods for details of experiments and data fitting.

| Peptide | Condition | $T_m$ (°C) | $E_a$ (kcal mol$^{-1}$) |
|---|---|---|---|
| OGP | No NaCl | 67.6 ± 2.5 | 59.7 ± 1.2 |
| | +154 mM NaCl | 67.5 ± 2.4 | 59.5 ± 1.1 |
| | Difference | 0.06 ± 0.06 | 0.73 ± 0.55 |
| KGD | No NaCl | 62.6 ± 1.9 | 70.8 ± 1.3 |
| | +154 mM NaCl | 60.7 ± 2.1 | 68.1 ± 1.2 |
| | Difference | 1.91 ± 0.25 | 2.64 ± 0.06 |
| KGE | No NaCl | 58.9 ± 2.1 | 73.3 ± 0.8 |
| | +154 mM NaCl | 56.3 ± 2.0 | 70.6 ± 0.7 |
| | Difference | 2.50 ± 0.19 | 3.28 ± 0.06 |

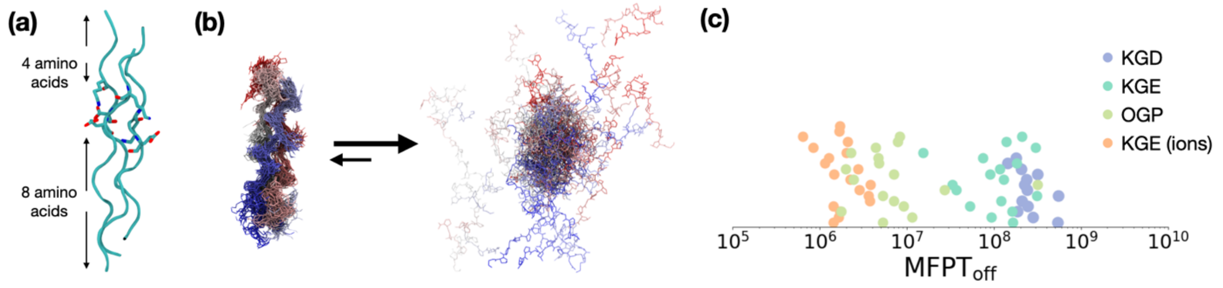


**Supplementary figure S9: Overview of the molecular dynamics (MD) simulation protocol.** **(a)** Simulated collage fragment, containing 4 amino acids before the triplet and 8 after. **(b)** The unfolding process aimed to target during the simulations. **(c)** The unfolding MFPT results for the simulations after bootstrapping classified by triplet.

**Supplementary table S4.** Total pathogenic missense mutations and those that are present within the salt-bridge footprint as aggregated from Clinvar[1], Leiden Open Variation Database (LOVD)[2] and Alport Syndrome Database[3].

| Collagen chain | Pathogenic mutations | Pathogenic mutations in salt-bridge footprint |
|---|---|---|
| col1α1 | 344 | 68 |
| col1α2 | 317 | 29 |
| col2α1 | 219 | 53 |
| col3α1 | 301 | 68 |
| col4α1 | 54 | 28 |
| col4α2 | 9 | 2 |
| col4α3 | 197 | 43 |
| co4α4 | 135 | 22 |
| col4α5 | 416 | 160 |
| col4α6 | - | - |
| col5α1 | 65 | 11 |
| col7α1 | 173 | 77 |
| col8α1 | - | - |
| col8α2 | - | - |
| col10α1 | 5 | - |
| col11α1 | 42 | - |
| col12α1 | 3 | - |
| col13α1 | 1 | - |
| col14α1 | - | - |
| col15α1 | 1 | - |
| col16α1 | - | - |
| col17α1 | 4 | - |
| col18α1 | - | - |
| col19α1 | - | - |
| col20α1 | - | - |
| col21α1 | - | - |
| col22α1 | - | - |
| col23α1 | - | - |
| col24α1 | 1 | 1 |
| col25α1 | 3 | 3 |
| col26α1 | - | - |
| col27α1 | 4 | - |
| col28α1 | - | - |
| **Total** | **2294** | **565** |

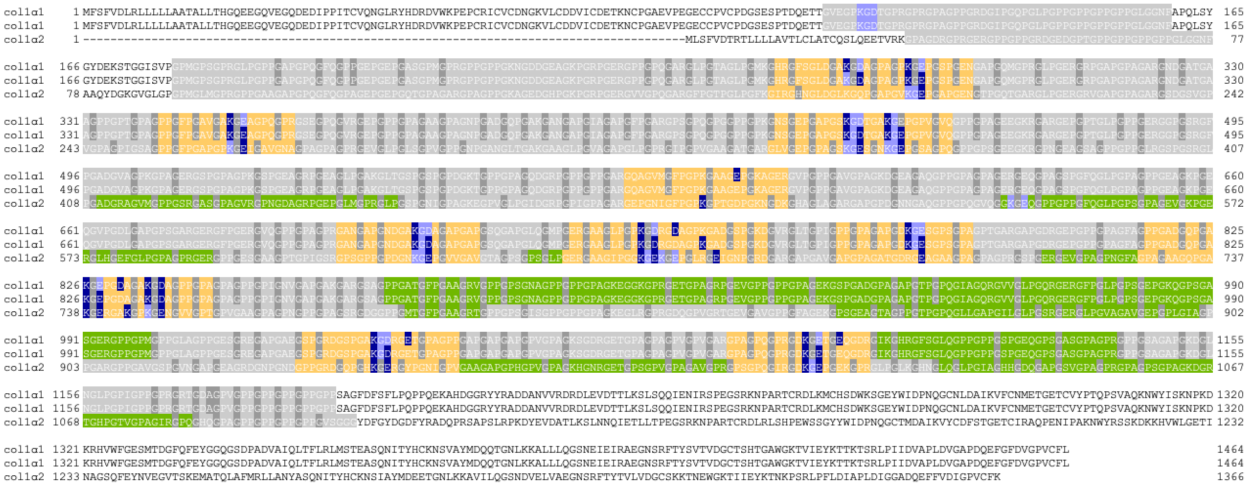

**Supplementary figure S10. Lethal mutations in the salt-bridge footprint of collagen type I.** The triple-helical domain of collagen type I (gray) indicating the previously observed lethal regions[4] (green), actual lethal mutations obtained from LOVD database (dark gray)[2], salt-bridges (purple) and their footprints (yellow).

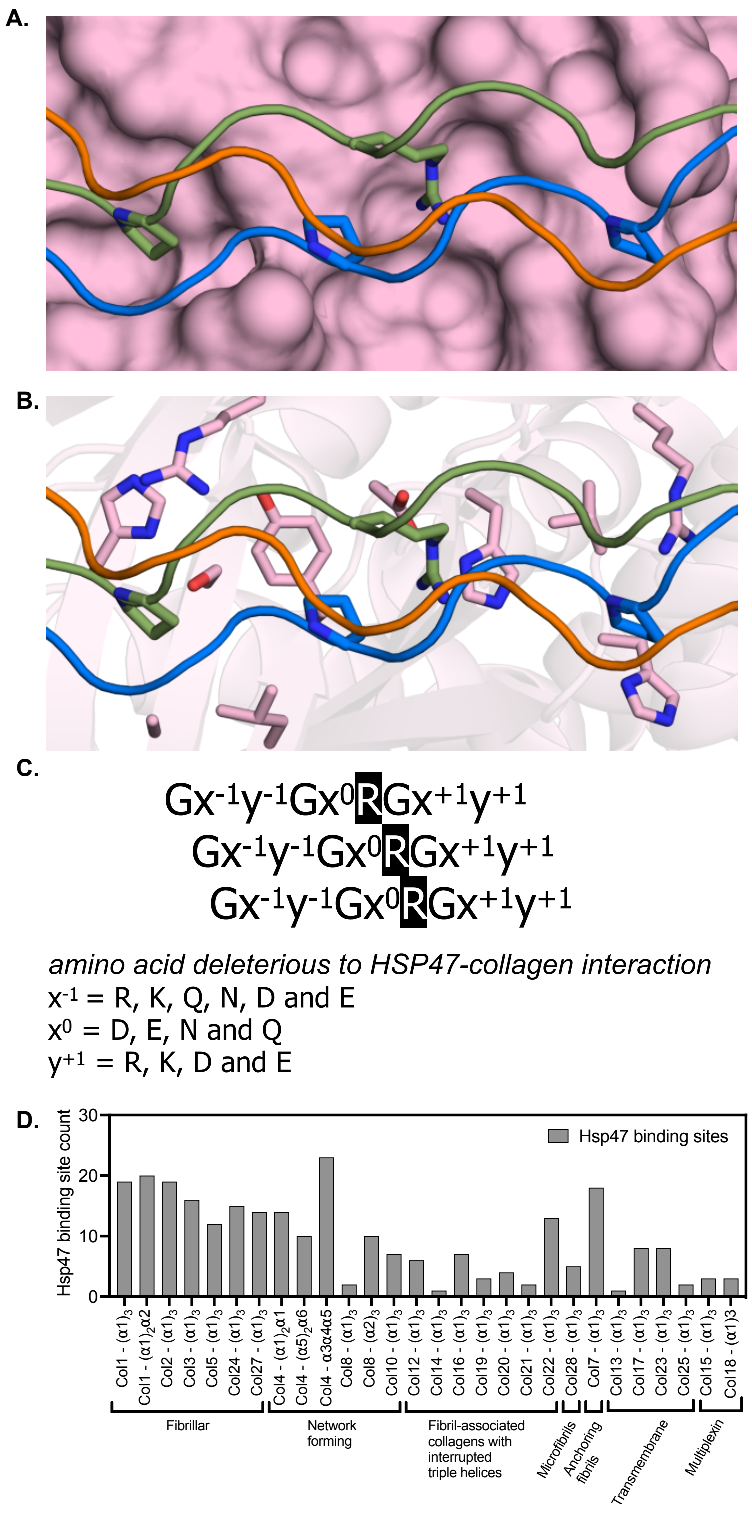


**Supplementary figure S11. Identification of Hsp47 binding sites in human collagen.** The binding interface of collagen with HSP47 shown as surface (A) and cartoon (B) representation. Interfacial residues identified as important for the interaction are shown as sticks (pdb: 7bdu). We obtained the sequence constraint show in **C** via analysis of the crystal structure and the peptide library scan used previously by Abraham et al for identifying potential HSP47 binding sites across the different collagen types[5]. Abraham et al used a peptide library to determine which amino acids in the minimal GxyGxRGxy motif minimize or abrogate Hsp47 recognition. In order to describe the sequence tolerance, the five positions (excluding glycine) are denoted as $Gx^{-1}y^{-1}Gx^{0}RGx^{+1}y^{+1}$. Their findings suggested that in $y^{-1}$ position, alanine results in medium affinity (1 ≤ IC50 < 10 μM), methionine results in low affinity (10 ≤ IC50 < 100 μM) binding while arginine, glutamine, lysine and glutamate abrogate binding. As shown in the peptide-Hsp47 crystal structure in **A** and **B**, sidechain of the $y^{-1}$ residue sits in a shallow hydrophobic pocket. Thus, polar residues are not well tolerated. We expect aspartate and asparagine to be similarly deleterious in this position. In the $x^{0}$ position, glutamate and asparagine abrogate binding. The sidechain of $x^{0}$ residue is positioned such that it can interact with the arginine that binds D385 of Hsp47. Thus, aspartate, glutamate, asparagine and glutamine are also expected to be deleterious as they can compete with Hsp47 for interaction with the primary Arg. In the $y^{+1}$ position, glutamate, aspartate or arginine are shown to abrogate binding. The deleterious effect of arginine is not suprising given its proximity to R228 and H274 residues of Hsp47. The glutamate and aspartate are deleterious likely because they can form salt-bridge to R228 or H274 and prevent the conformational rearrangement of the loop containing these residues as required to accommodate the bulky hydrophobic amino acid in the $x^{+1}$ position. Focusing on the deleterious amino acids, a sequence where $x^{-1}$ = R, K, Q, N, D and E, $x^{0}$ = D, E, N and Q or $y^{+1}$ = R, K, D and E would likely abrogate Hsp47-collagen interaction. Hsp47 only recognizes folded triple helices and not single polypeptides. Thus, we searched aligned triple helices show in supplementary figure S2 for matches excluding the amino acid combinations that abrogate binding. Low and medium affinity amino acids are intentionally included as high intracellular concentration of Hsp47 render these binding sites viable. As shown in **D** the 265 sequence motifs across all collagen types that could recognize Hsp47 are overwhelmingly present in the fibrillar and network-forming collagens. Tellingly, collagen types XIII and XIV contain only one binding site while types XXI, XXV, VIII, XV, XVIII and XIX contain three or less binding sites. The single binding site found for type XIV also supports its previously observed weak affinity for Hsp47.

References

1. Landrum, M. J. *et al.* ClinVar: improving access to variant interpretations and supporting evidence. *Nucleic Acids Res.* **46**, D1062–D1067 (2018).
2. Fokkema, I. F. A. C. *et al.* LOVD v.2.0: The next generation in gene variant databases. *Hum. Mutat.* **32**, 557–563 (2011).
3. Crockett, D. K. *et al.* The Alport syndrome COL4A5 variant database. *Hum. Mutat.* **31**, E1652-7 (2010).
4. Marini, J. C. *et al.* Consortium for osteogenesis imperfecta mutations in the helical domain of type I collagen: Regions rich in lethal mutations align with collagen binding sites for integrins and proteoglycans. *Hum. Mutat.* **28**, 209–221 (2007).
5. Abraham, E. T. *et al.* Collagen's primary structure determines collagen:HSP47 complex stoichiometry. *J. Biol. Chem.* **297**, 101169 (2021).
